# Inflammatory pain resolution by mouse serum-derived small extracellular vesicles

**DOI:** 10.1101/2024.02.16.578759

**Authors:** Zhucheng Lin, Xuan Luo, Jason R. Wickman, Deepa Reddy, Richa Pande, Yuzhen Tian, Vivian Triana, Jingyun Lee, Cristina M. Furdui, Desmond Pink, Ahmet Sacan, Seena K. Ajit

## Abstract

Chronic pain is a significant public health issue. Current treatments have limited efficacy and significant side effects, warranting research on alternative strategies for pain management. One approach involves using small extracellular vesicles (sEVs) to transport beneficial biomolecular cargo to aid pain resolution. Exosomes are 30-150 nm sEVs that can carry RNAs, proteins, and lipid mediators to recipient cells via circulation. Exosomes can be beneficial or harmful depending on their source and contents. To investigate the short and long-term effects of mouse serum-derived sEVs in pain modulation, sEVs from naïve control or spared nerve injury (SNI) model donor mice were injected intrathecally into naïve recipient mice. Basal mechanical thresholds transiently increased in recipient mice. This effect was mediated by opioid signaling as this outcome was blocked by naltrexone. Mass Spectrometry of sEVs detected endogenous opioid peptide leu-enkephalin. A single prophylactic intrathecal injection of sEVs two weeks prior to induction of the pain model in recipient mice delayed mechanical allodynia in SNI model mice and accelerated recovery from inflammatory pain after complete Freund’s adjuvant (CFA) injection. ChipCytometry of spinal cord and dorsal root ganglion (DRG) from sEV treated mice showed that prophylactic sEV treatment reduced the number of natural killer (NK) and NKT cells in spinal cord and increased CD206+ anti-inflammatory macrophages in (DRG) after CFA injection. Further characterization of sEVs showed the presence of immune markers suggesting that sEVs can exert immunomodulatory effects in recipient mice to promote the resolution of inflammatory pain. Collectively, these studies demonstrate multiple mechanisms by which sEVs can attenuate pain.

## Introduction

Current drugs designed to manage chronic pain display low efficacy and many side effects, which warrants developing alternative therapeutic strategies. Though many studies have investigated soluble pain mediators including neurotransmitters, cytokines, chemokines, and small peptides, the roles of extracellular vesicles (EVs) are less explored^1^. EVs are heterogeneous populations of vesicles released from cells into the extracellular space^2^. EVs transfer cargo molecules between cells and thereby impact the function of recipient cells. EVs are categorized based on vesicle size, density, morphological features, cargo, and biogenesis, into three major types: exosomes, microvesicles, and apoptotic bodies. Exosomes are 30-150 nm in diameter and derived from multivesicular bodies. Microvesicles are generated from the outward budding and fission of the plasma membrane and range in size from 150-1000 nm. Apoptotic EVs can be up to 5000 nm and are released by cells undergoing apoptosis. Thus, EVs range in diameter from 30 nm to greater than 1000 nm and can be detected in all body fluids and conditioned culture media^3^. Since exosomes and microvesicles overlap in size, the term small extracellular vesicles (sEVs) describes all vesicles less than 200 nm in diameter, as recommended by the International Society of Extracellular Vesicles^4^. Cells use sEVs to deliver cargoes of proteins, lipids, and nucleic acids, thereby communicating with other cells over long distances^2^. sEV contents vary depending on cellular source as well as physiological or pathological conditions of the donor cells. In addition to the variable contents under different states, sEVs can have numerous effects on different recipient cell types^5^.

sEVs have been studied for their utility as biomarkers and therapeutics as well as mediators of intercellular communication and signaling^6, 7^. Several studies have investigated the role of sEVs in neurological disorders including pain^1, 8, 9^. For example, sEVs from mesenchymal stem cells could attenuate neuropathic pain in a rodent pain model and were proposed as a novel strategy to attenuate nerve injury-induced pain^8, 10^. In osteoarthritis-induced inflammatory pain, intraarticular administration of bone marrow stem cell-derived exosomes attenuated both thermal and mechanical allodynia in rats^11^. Our studies in mice have shown that macrophage-derived sEVs can alleviate inflammatory pain from complete Freund’s adjuvant (CFA)^12, 13^ together suggesting that sEVs can attenuate different types of pain. Most similar studies investigated exosomes derived from various types of cultured cells.

Our recent proteomics study using sEVs purified from mouse serum four weeks after spared nerve injury (SNI) showed the presence of commonly expressed sEV proteins and differential expression of both anti and pro-inflammatory molecules compared to control sEVs from the serum of naïve and sham-operated mice^14^. For example, the pro-inflammatory complement 5a (C5a) protein was significantly upregulated in sEVs from SNI model mice^14^. Another study reported downregulation of miRNAs that suppress inflammation in plasma-derived EVs from the rat spinal nerve ligation model of neuropathic pain^15^. Based on changes in pronociceptive C5a^16^ and miRNAs^15^, we hypothesized that the protein and miRNAs in sEVs released under neuropathic pain could sensitize mice to pain. However, our studies using sEVs from SNI model and naïve mice show that sEVs from mouse serum help to resolve pain in recipient mice. These outcomes demonstrate the immunomodulatory effects of sEVs whereby administration of sEVs altered immune cell composition in spinal cord and dorsal root ganglia (DRG) of recipient mice. We also report a comprehensive multi-modal analysis of different aspects of sEV characterization, miRNAs, and protein markers in addition to pain behavior testing in recipient mice.

## Methods

### Mice

All the studies were performed following the NIH guidelines and the protocols were approved by the Institutional Animal Care and Use Committee of Drexel University College of Medicine. For behavior studies, we used 8- to 10-week-old male and female C57BL/6 mice purchased from the Jackson Laboratory (Bar Harbor, ME). All mice were housed with a 12-h light/dark cycle and provided with food and water *ad libitum*.

### Rodent models of pain

Spared nerve injury (SNI) model mice were generated as follows^17^. Mice were anesthetized using 2% isoflurane before and during surgery. The surgical procedure included ligation and transection of the tibial and common peroneal nerves of the left hind paw, but the adjacent sural and saphenous nerves were left intact. For sham control surgery, animals underwent the same procedure, but without nerve transection. Complete Freund’s adjuvant (CFA) model mice were generated by injecting 20 µL of 50% emulsified CFA (Sigma-Aldrich, St. Louis, MO) in saline into the right hind paw.

### Behavioral testing

Mice were habituated in the testing room two-to-three days before baseline tests. The experimenter was blinded to the treatment conditions. Mechanical allodynia was assessed by von Frey method and thermal pain threshold was evaluated by the Hargreaves method as described^13^. Mice were habituated in the dynamic weight bearing (DWB) enclosure (Bioseb, Pinellas Park, FL) for 10 min one day prior to baseline. Following a 5-min habituation, we recorded and analyzed posture, weight distribution, and paw surface area of each of the four paws, allowing for precise tracking of postural and weight shifts^18, 19^. The ratio of body weight distribution on hind paws was measured at multiple time points after injections.

### Isolation of sEVs from mouse serum

A combination of differential ultracentrifugation, ultrafiltration, and size-exclusion chromatography (SEC) was used to isolate sEVs from mouse serum. All centrifugations were performed at 4°C. Briefly, two weeks after sham or SNI surgeries, sham control, SNI model and age-matched naïve mice were anesthetized by isoflurane. Whole blood was collected in 1.7 mL microcentrifuge tubes without anticoagulant and incubated undisturbed at room temperature for 45 min to allow clot formation. The supernatant serum was collected after centrifugation at 2,000 × g for 10 min and stored at -80 °C until use. The serum sample was diluted with an equal volume of DPBS without calcium and magnesium then centrifuged for 30 min at 2,000 × g to pellet cell debris. The supernatants were then centrifuged at 12,000 × g for 45 min and suspension was filtered through a 0.22 μm syringe filter. Samples were then diluted to 4 mL with DPBS and transferred to 100 K Amicon Ultra Centrifugal Filters (Sigma-Aldrich), followed by centrifugation at 5,000 × g for 30 min. The concentrated sEVs were diluted to 500 µL with DPBS and purified by SEC using qEVoriginal 35 nm Legacy columns (iZON, Medford, MA) following the manufacture’s manual. Four EV-rich fractions (7-10, 0.5 mL each) were pooled and after ultracentrifugation at 110,000 × g for 70 min (Optima TLX ultracentrifuge with TLA 100.4 rotor, Beckman Coulter, Inc.), the pellets were resuspended in DPBS and stored at -80 °C until further use.

### Isolation of sEVs from cell supernatant

RAW 264.7 macrophage cells (ATCC) were cultured to 70% confluency in DMEM supplemented with 10% exo-depleted FBS and 100 U/mL pen-strep in the presence or absence of 1 μg/mL LPS (Sigma-Aldrich) at 37°C with 5% CO_2_. Conditioned media was collected at 24 hours for sEV isolation. Briefly, media was centrifuged at 500 x g for 10 minutes at 4°C followed by 12,000 x g for 35 minutes at 4°C. The supernatant was filtered using 0.22 μm syringe filter and supernatant concentrated to a final volume of 500 µL with 100 kDa concentrator (Amicon Millipore Sigma) by centrifugation at 5,000 x g at 4°C for 30 minutes in a fixed angle rotor (Beckman Coulter). The qEV column washed with 1x PBS was then loaded with 500 µL of concentrated supernatant followed by 2.5 mL PBS to collect 3 mL void volume. After the column flow stopped, 2 mL of PBS was added to collect EV zone sample. The EV volume (2 mL) was then centrifuged for 70 minutes at 110,000 x g at 4°C and sEV pellet resuspended in DPBS.

### Nanoparticle tracking analysis (NTA)

Brownian motion of sEV particles from mouse serum were visualized and their size distribution and concentration measured using NanoSight NS300 (Malvern Instruments). Briefly, sEV samples were diluted in DPBS and placed into sample chamber by syringe pump. The combination of shutter speed and gain were set to obtain 30 s videos which were analyzed by the NanoSight NTA 3.1.54 software.

### Western blotting

Cell or sEV samples were re-suspended in RIPA buffer (Sigma-Aldrich) with protease inhibitors (Thermo Fisher Scientific). Protein concentrations were measured by Micro BCA Protein Assay kit (Thermo Scientific) for sEVs or DC protein assay (Bio-Rad, CA, USA) for cell lysates. Equal amounts of protein samples were mixed with Laemmli SDS reducing sample buffer (Thermo Scientific), loaded on 10% Tris-Glycine Gel (Thermo Fisher Scientific) and the gel was run at 125 V for 90 min. Proteins were transferred to 0.4 µm PVDF membrane at 25 V for 90 min, followed by blocking the PVDF membrane with Odyssey Tris-Buffered Saline (TBS) blocking buffer (LICOR Biosciences, NE, USA) for 1 h at room temperature. The membranes were incubated with primary antibodies in TBS with 0.1% TWEEN 20 (TBST) and 10% (v/v) blocking buffer on shaker overnight at 4 °C. The blots were washed with TBST thrice, then the blots were incubated with secondary antibodies in TBST at room temperature for 1 h on the shaker. The membranes were washed with TBST thrice and protein detected by LI-COR Image Studio Software (LI-COR Biosciences). Primary antibodies used were: mouse anti-CD81 (1:500, #sc-166029, Santa Cruz Biotechnology), rabbit anti-Hsp70 (1:2000, #ab94368, Abcam), rabbit anti-albumin (1:2000, #16475-1-ap, Proteintech). Secondary antibodies used were 680RD donkey anti-mouse IgG (#926-68072, LI-COR Biosciences), and 800CW donkey anti-rabbit IgG (#926-32213, Li-COR Biosciences).

### Preparation of sEV samples for transmission electron microscopy (TEM)

sEV samples were prepared as previously described^12^. The grids were incubated with primary antibody rabbit anti-CD81 (1:100, #SAB3500454, Sigma) for 30 min at room temperature. Unbound antibodies were removed using washing buffer (0.1% BSA in 0.1 M PB) and grids were incubated with 6 nm gold-conjugated donkey-anti-rabbit IgG (1:25, #25104, Electron Microscopy Sciences) for 20 min.

### Labeling of sEVs

The labeling studies were performed as described^20^. Purified sEVs were diluted in 1 mL of diluent C (Sigma-Aldrich) and mixed with 3 µL PKH26 dye. An equal volume of PBS as that of the sEV suspension was used for dye control. After incubation for 5 min at room temperature in dark, 2 mL of 1% BSA in DPBS was added to absorb excess dye. Samples were ultracentrifuged at 110,000 × g for 70 min. Labeled sEV pellets were resuspended in DPBS and centrifuged again under 110,000 × g for 70 min to remove any remaining unlabeled dye. Ten µg of labeled sEVs were used in each intrathecal injection.

### Intrathecal sEV administration

Eight to ten-week-old C57BL/6 mice were used for intrathecal injections. All injections were performed with a Hamilton syringe and 30-gauge needles. After identification of injection site, the needle was inserted into the tissue to the intervertebral space between L4 and L5, and a successful puncture resulted in a tail flick. Then 10-15 μL of solution was slowly injected.

### Perfusion, tissue fixation and immunohistochemistry

Mice were anesthetized by intraperitoneal injection of 100 mg/kg body weight of ketamine and 10 mg/kg body weight of xylazine. Briefly, intracardial perfusion of mice with 0.9% saline to flush out blood was followed by prechilled 4% paraformaldehyde (PFA) in phosphate buffer for 5 min to fix mouse tissues. Dissected L4-L5 spinal cord and DRG were immersed in 4% PFA in PB at 4°C for 24 h. Cryopreservation was done with 30% sucrose in PB at 4°C for at least 24h. DRG or spinal cord tissues were embedded in O.C.T. (optimal cutting temperature) compound and frozen at -20°C for 20 min. Using cryostat Microm HM550 (Thermo Scientific), tissues were sectioned at 30 μm and washed thrice with 0.3% Triton in PB for 5 min. Then sections were blocked with 5% normal goat serum (NGS) and 0.3% Triton in PB for 2 hours at room temperature. Sections were incubated with primary antibodies with 5% NGS and 0.3% Triton in PB on shaker overnight at 4°C. Sections were washed thrice with 0.3% Triton in PB then incubated with secondary antibodies with 5% NGS in PB at room temperature for 2 h on the shaker. Sections were washed with PB thrice and mounted on a clean slide (Superfrost Plus Gold). Coverslips with DAPI-mounting solution (Abcam, Catalog# ab104139) were used to mount and slides were placed in the dark overnight. Slides were imaged using Olympus FV3000 microscope. Primary antibodies used were: Anti-MAP2A (1:500, #M9942, Sigma-Aldrich), GFAP (1:1000, #ZRB2383, Sigma-Aldrich), Iba1 (1:2000, #019-19741, Wako Chemicals). Secondary antibodies used were donkey anti-rabbit IgG Alexa Fluor 546 (1:500, Invitrogen), or goat anti-mouse IgG Alexa Fluor 488 (1:500, Invitrogen).

### Whole-transcriptome microRNA profiling and bioinformatics analysis

Serum-derived sEVs from naïve, sham, and SNI mice (n=4) were diluted in a 1:1 ratio with HTG Biofluids Lysis Buffer (HTG Molecular Diagnostics, Inc., Tucson, AZ), and stored at -80°C. All samples (25 µL/sample) were run and sequenced with the HTG EdgeSeq miRNA Whole Transcriptome Assay on an Illumina MiSeq Sequencer to measure the expression of 2,083 miRNA transcripts. Reads of each miRNA were normalized to transcripts per kilobase million (TPM) then log2-transformed. We used principal component analysis for the preliminary analysis to identify any outliers. The correlation between pairs of genes across samples and the correlation between pairs of samples were analyzed by hierarchical clustering method using the Pearson correlation coefficient. Low-expressed miRNAs were filtered out if a miRNA had a TPM value <1 in all samples. For the presence–absence analysis, a miRNA was considered present in a group if it had a TPM value of ≥1 in every replicate of that group. Differential expression analysis was performed between two experimental groups using a permutation test and differentially expressed genes were defined as having a Type I error *p*-value <0.01 and an absolute fold change ≥2. Gene targets of differentially expressed miRNAs were identified using TargetScan with an aggregate P_CT_ >0.9 ^21^. Enriched Gene Ontology (GO) biological processes and Kyoto Encyclopedia of Genes and Genomes (KEGG) pathways were identified from the predicted targets using the Database for Annotation, Visualization and Integrated Discovery (DAVID) Bioinformatics Resources (2021 Update). Significant annotations (FDR <0.01) were summarized to maintain <80% overlap among them.

### Mass spectrometry

Five µg of sEV suspension was subjected to extraction procedure. Briefly, sample volume was adjusted to 50 µL with PBS and 50 µL of hypertonic buffer (2.2 M sucrose, 1 mM MgCl2, 10 mM Tris-HCl, pH 7.4) was added and vortexed. Electroporation was performed with the mixture using the Multiporator (Eppendorf, Hamburg, Germany; 500 V, 200 µsec, 5 cycles). Samples were purified using the Pierce C18 Tips (Cat# 87784, Thermo Scientific, Waltham, MA, USA) and then prepared in 5% acetonitrile with 1% formic acid for LC-MS/MS analysis. Samples were analyzed on a LC-MS/MS system which consisted of an Orbitrap Velos Pro Mass Spectrometer (Thermo Scientific, Waltham, MA, USA) and a Dionex Ultimate-3000 nano-UPLC system (Thermo Scientific, Waltham, MA, USA). An Acclaim PepMap 100 (C18, 5 μm, 100 Å, 100 μm x 2 cm) trap column and an Acclaim PepMap RSLC (C18, 2 μm, 100 Å, 75 μm x 50 cm) analytical column were employed for peptide separation. Data was acquired using parallel reaction monitoring (PRM) acquisition by collecting peptide specific product ions (MS2) from collision-induced dissociation (CID) of parent ions targeted in the survey scan (MS1). Target ions were as follows; for β-endorphin 688.1740 m/z (z=5), Met-enkephalin 574.2330 m/z (z=1), Leu-enkephalin 556.2766 m/z (z=1), nociceptin 603.6675 m/z (z=3), endomorphin-1 612.2817 m/z (z=1), endomorphin-2 573.2708 m/z (z=1), and dynorphin A 537.5551 m/z (z=4). Peptides were confirmed by comparing their peak retention time and fragment ion profile to the ones of authentic peptide standards. β-endorphin (#LS-H14937-1), met-enkephalin (#LS-H4423-25), leu-enkephalin (#LS-H4826-25), nociception (#LS-H5242-1), endomorphin-1 (#LS-H13168-1), and endomorphin-2 (#LS-H13038-1) were purchased from LifeSpan BioSciences (Seattle, WA, USA), and dynorphin A (#18169) was purchased from Cayman Chemical (Ann Arbor, MI, USA). Skyline (MacCoss Lab Software, University of Washington, Seattle, WA, USA) was used for target ion detection, peak feature extraction, and area calculation for quantitation. Leu-enkephalin was also measured using EIA kit (#FEK-024-21) (Phoenix Peptide, CA) following manufactures protocol. Measurements were performed in duplicate using pooled (n=10 per group) serum and sEV samples that had been processed on C18 columns (#RK-SEPCOL-1), concentrated in a speed-vac then lyophilized overnight and resuspended in 100 µL.

### Administration of drugs

Morphine (Cayman Chemical 15464) and naltrexone (Sigma-Aldrich, 1453504) were diluted in sterile PBS. For *in vivo* studies, naltrexone was administered intraperitoneally with a dose of 10 mg/kg 2h 30 min before 0.05 mg/mL morphine solution was injected intrathecally at a volume of 10 μL per mouse 30 min prior to behavior test.

### ChipCytometry

Mice were perfused with ice-cold 0.9% saline. Fresh L4-L5 spinal cord and ipsilateral DRG were dissected and snap-frozen in cryo mold containing O.C.T. compound in dry ice. Frozen tissues were stored at -80°C until sectioning. To obtain cryosections, fresh frozen tissues were sectioned at 5-7 µm each at -20°C on coated coverslips immediately followed by acetone/ethanol fixation and chip preparation at 4°C according to the manufacturer’s protocol. Chips were shipped at 4°C to Canopy Biosciences for antibody probing and imaging. Images were acquired using ZellScanner and ScanApp and analyzed using ZKWapp. Following antibodies were used: FITC anti-mouse CD206 (1:100, #141703, BioLegend), PE anti-mouse NeuN (1:3000, #NBP1-92693PE, Novus), FITC anti-human/mouse CD45R (1:500, #11-0452-82, Invitrogen), PE anti-mouse CD3ε (1:500, #100308, BioLegend), PerCP-Cy5.5 anti-mouse CD4 (1:300, #100540, BioLegend), BUV395 anti-mouse CD8α (1:200, #565968, BD Biosciences), PE Ki67 (1:100, #12160, Cell Signaling Technology), PerCP-Cy5.5 anti-mouse/human CD11b (1:200, #101227, BioLegend), FITC anti-mouse MHCII (1:500, #11-5321-82, Invitrogen), PE anti-mouse CD69 (1:200, #104508, BioLegend), PE anti-mouse FOXP3 (1:50, #12-5773-82, Invitrogen), PerCP-Cy5.5 anti-mouse F4/80 (1:500, #123126, BioLegend), PE anti-mouse CD335 (1:100, #137604, BioLegend), FITC anti-mouse CD45 (1:1500, #11-0451-82, Invitrogen), PE anti-mouse CD19 (1:100, #557399, BD Biosciences), and PE anti-mouse Gr-1 (1:1000, #108407, BioLegend). DNA was stained with Hoechst 33342 (1:5000, #H21492, Thermo Fisher). Experimental design and the 16-plex panel of markers used are shown in Supplementary Tables 4, 5.

### ExoView single vesicle assay

ExoView-Exoflex chip with capture antibody^22^ against EV tetraspanin proteins CD9, CD81, and custom targets CD45 and F4/80 (NanoView Biosciences Inc.) was used to study sEVs from RAW 264.7 cell culture media, naïve and SNI model mice. ExoView assay was performed according to manufacturer’s protocol. Briefly, samples were diluted four times in incubation buffer, then 35 μL sample was incubated on the ExoView chip overnight (16 hours) at room temperature. Chips were then washed and stained for 1 h with detector antibody cocktail. Tetraspanin antibodies were obtained from NanoView Biosciences. CD45 (#103102) and F4/80 (# 123101) antibodies were purchased from Biolegend Inc. Following staining, chips were washed, dried, and imaged with the R100 reader. NanoViewer Analysis software 3.04 was used to calculate the particle count and colocalization for each capture spot.

### Microflow cytometry and NTA for sEVs

All microflow cytometry samples were assayed on a calibrated Apogee A60 MicroPlus with three angles of light scatter collection. The instrument was cleaned with a 1% bleach solution, and calibrated daily as per vendor protocols and using a bead mixture (Apogee 1524 bead mixture) to standardize instrument fluorescence and counting properties; calibration was checked using a monitoring bead mixture (Apogee 1527 bead mixture). Instrument cleanliness was assessed daily, prior to any sample assay, using both sheath and also diluent (PBS) blanks. Samples were diluted in PBS prior to analysis. Samples incubated with CFDA-SE to stain membranes were treated with or without Triton X-100 prior to assay as internal vesicle controls. Samples were assayed for 120 seconds at 3.01 uL/min.

For NTA performed at Nanostics, all samples were analyzed using a NanoSight LM10 system equipped with a 405 nm laser (NanoSight, Amesbury, U.K.; Malvern Panalytical). Prior to sample analysis, a series of NIST traceable polystyrene beads (3000 series, Thermo Scientific, Waltham, MA, U.S.A.) were assayed to ensure that the instrument was working under optimal functionality. Beads were diluted in 10 mM KCl buffer filtered through a 0.1 µm filter (Acrodisc® 25mm w/0.1µm, Cat#4611, Pall Corporation) to achieve a concentration of 20–90 particles/frame and injected into the sample chamber with sterile syringes. EV samples were diluted in PBS to achieve a similar concentration of 20–90 particles/frame and injected into the sample chamber with sterile syringes. Five technical replicates from each sample were measured, each run for 1 min. The temperature during sample acquisition was recorded to accurately determine particle concentration. The unit was cleaned between samples and instrument cleanliness was checked prior to execution by using diluent.

### Detection of sEV surface epitopes by standard flow cytometry

The mouse MACSPlex exosome kit (Miltenyi Biotec, 130-122-211) was used to detect EV specific markers using BD LSRFortessa (BD Biosciences) following the manufacturer’s protocol. Briefly, 10 µg of sEVs were diluted to 120 μL using the MACSPlex Buffer. The sEVs were incubated with 15 μL of MACSPlex exosome capture beads overnight at room temperature using an orbital shaker (450 rpm). The following day, 500 μL of MACSPlex buffer was added and the samples were centrifuged at 3,000 × g for 5 minutes. After aspirating the supernatant, 15 μL of MACSPlex exosome detection reagent CD9, CD63, or CD81 cocktail was added to each tube and incubated for an hour using an orbital shaker as mentioned above. After washing and centrifugation, the tubes were incubated for 15 minutes at room temperature on orbital shaker. MACSPlex exosome capture beads with EVs were resuspended by pipetting and subjected to flow cytometry. sEVs were analyzed for various markers such as CD9, CD63, CD81, CD45, CD11b, CD11c, MHC-I and MHC-II, isotype control (IgG), CD19, CD25, CD49b expression. Observed (Mean Fluorescence Intensity) MFI was normalized to the highest observed MFI across the groups. Weighted expression of samples is represented in the form of heat map.

### Statistical analysis

Data are presented as mean ± SEM. Group differences of sEV particle concentration and size in NTA were compared by the one-way analysis of variance (ANOVA) and the multiple comparisons between means were tested by the *post hoc* Bonferroni method. All behavior data were analyzed using two-way ANOVA, and pairwise comparisons between means were tested by the *post hoc* Bonferroni test. The differences between groups were considered significant when the *p*-value was less than 0.05. GraphPad Prism software (version 10) was used for all statistical analysis.

## Results

### sEV characterization

We first confirmed that SNI model mice developed mechanical pain hypersensitivity compared to naïve and sham surgery mice (**Fig. 1a**). Two weeks after surgery, we collected serum samples from all three groups of mice. Nanoparticle tracking analysis (NTA) analysis of sEVs showed there was no statistically significant differences in particle concentrations from naïve, sham and SNI model mice. The concentrations were 8.7 ± 1.6 x 10^8^, 6.0 ± 1.0 x 10^8^ and 5.6 ± 0.7 x 10^8^ particles/mL respectively (**Fig. 1b**). sEVs from SNI model mice were slightly smaller than the two control groups (**Fig. 1c**). NTA showed the mean diameters were 121.1 ± 4.1 nm for naïve sEVs, 115.7 ± 5.2 nm for sham sEVs and 101.1 ± 2.4 nm for SNI sEVs respectively (**Fig. 1c**). sEV particle size distribution and diameter range was 50-150 nm, which is a typical diameter range for exosomes^23^ (**Fig. 1d**). Additional NTA studies done at Nanostics are shown in **Fig. 10**. We also imaged sEVs via TEM and demonstrated they are positive for the sEV marker CD81 via immunogold labeling (**Fig. 1e**). Western blotting of sEVs and sEV-depleted serum from different groups of mice confirmed that sEVs expressed the exosome marker protein CD81 but not the negative control protein albumin (**Fig. 1f**). These results indicate that sEVs isolated from the serum meet the MISEV guidelines^4^ for purity.

**Fig. 1.**
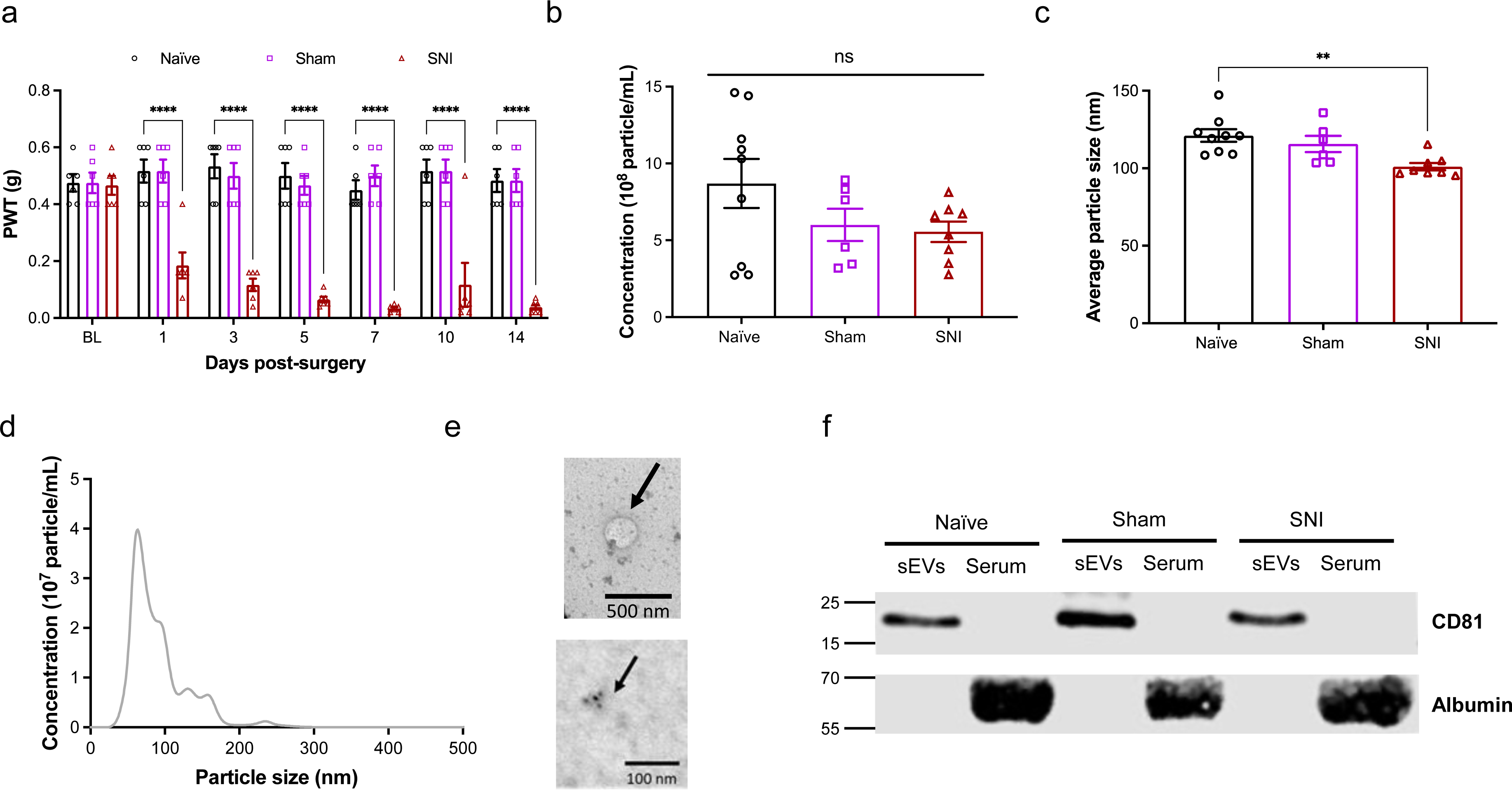
Characterization of mouse serum-derived sEVs. **a** Confirmation of mechanical hypersensitivity in male mice after SNI. Baseline testing (BL) was followed by SNI or sham surgery and mice were tested for mechanical sensitivity for 14 days. Animals were sacrificed after two weeks and sEVs isolated from their serum were used in all studies. Data shown are mean ± standard error of the mean, n=6. Statistical analysis was determined by two-way repeated-measures ANOVA followed by Tukey posttest ** p <0.01, *** p <0.001, **** p <0.0001. **b** Nanoparticle tracking analysis (NTA) showing the mean of concentrations of sEVs in serum of naïve, sham control and SNI model mice respectively. **c** The sEV particle size distribution shows particle sizes of sEVs from SNI model mice were smaller compared to the control groups. The mean diameters were 121.1 ± 4.1 nm for naïve, 115.7 ± 5.2 nm for sham and 101.1 ± 2.4 nm for SNI sEVs. One-way ANOVA followed by Tukey posttest ** p <0.01. **d** Particle size distribution and diameter range for sEVs from naïve mice **e** TEM images showing sEV morphology and size (top) and immunogold labeling for CD81 in sEVs (bottom) from SNI model. **f** Western blot for sEV markers and sEV-depleted serum from different groups of mice confirming the presence of exosome marker protein CD81. Negative marker albumin was absent in sEVs but present in the serum.

### miRNA profiling of sEVs

Our previous study used sEVs collected at four weeks post-injury and profiled by qPCR for ∼750 miRNA using Taqman low density array^24^. Here we used two weeks post-injury sEVs and measured their expression of 2,083 human miRNA transcripts using HTG EdgeSeq miRNA Whole Transcriptome Assay. More importantly, here we used ultracentrifugation combined with size-exclusion chromatography to isolate sEVs and improve their purity. Using these improved samples, we tested if miRNA cargo differed in sEVs from the serum of SNI model, sham control and naïve mice. One of the naïve sEV samples was considered an outlier based on the principal component analysis and hierarchical clustering of all 12 samples (**Fig. 2a**) and excluded from further analyses. Several miRNAs were differentially expressed (Supplementary Table 1), as summarized in a Venn diagram showing unique and overlapping miRNAs in sEVs from the serum of SNI, sham, and naive control mice (**Fig. 2b**). A total of 1584 miRNAs were present in all three groups. Each group contained exclusive miRNAs, including 40 in sEVs from naive control mice, 81 in sham surgery control mice, and 57 in sEVs from SNI model mice. We documented a subset of differentially expressed miRNAs with the highest fold change between naïve and SNI model mice (**Fig. 2c**) as well as a subset of the most highly expressed miRNAs in all serum derived sEVs. The complete list is included in Supplementary Table 2. Our miRNA analysis shows that SNI injury alters the composition of miRNAs within sEVs, suggesting changes to upstream sEV packaging. The genes predicted to be targeted by the miRNAs that were differentially expressed in SNI vs. Naïve groups were enriched in numerous signaling pathways, including CAMP, MAPK, and Insulin signaling (**Fig. 2d**). The full list of significantly enriched pathways for each experimental group is available in Supplementary Table 3.

**Fig. 2.**
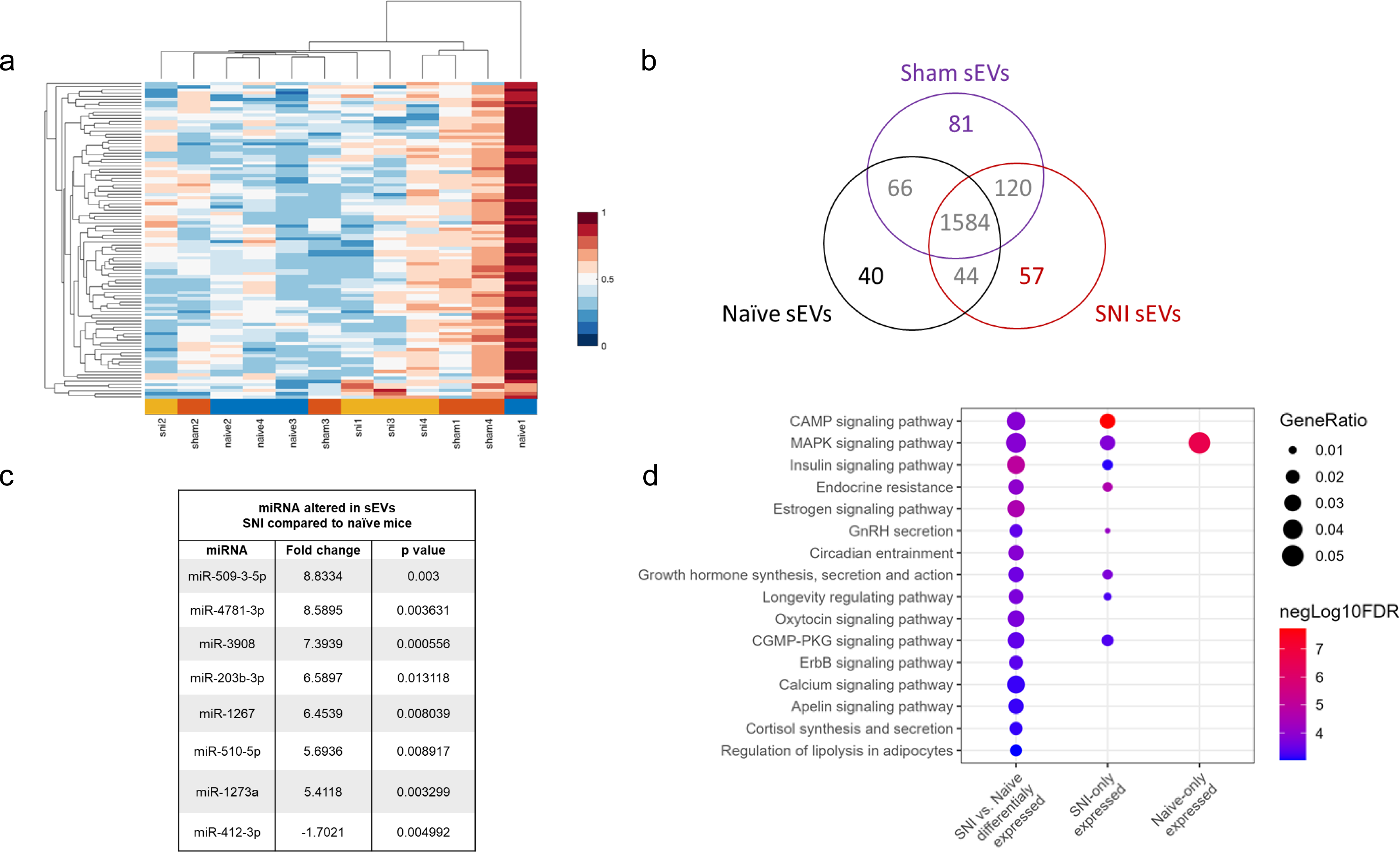
miRNAs in sEV samples. **a** Principle component analysis for miRNAs in serum-derived sEVs from naïve, sham control, and two weeks post SNI model mice. One of the naïve sEV samples (ID: naive1) was considered as an outliner and was excluded from further analyses. Hierarchical clustering analysis of overall miRNA expression level with higher expression denoted by red and lower by blue shown and more related terms are closely grouped together. Pairwise distances between miRNAs shown for SNI model, sham and naïve control mice (n=3-4). **b** Venn diagram showing presence/absence analysis of miRNAs in sEVs from serum. HTG EdgeSeq/RNA-Seq detected unique and overlapping miRNAs in sEVs purified from the serum of SNI, sham, and naive control mice (n=4) two weeks after surgery. **c** A subset of differentially expressed miRNAs with the highest fold change in sEVs from SNI model compared to naïve control. **d** Gene Ontology (GO) terms and KEGG pathways enriched for the genes targeted by the miRNAs. Annotations are filtered to reduce redundancy, with no more than 80% overlap in the target genes. Only the annotations (FDR<=0.01) resulting from the miRNAs that are significantly differentially expressed between SNI vs. naive conditions are shown. Additional annotations from miRNAs expressed explicitly in SNI or naive conditions are available in the supplementary files.

### sEV uptake in spinal cord and DRG

We next labeled sEVs from naïve mice with the fluorescent dye PKH26 as described^20^ to confirm that sEVs are taken up by the spinal cord and DRG. Ten µg of labeled sEVs were intrathecally injected into each naïve mouse, and 6 h later, we collected the spinal cord and L4-L5 DRG tissues. Tissues were stained for markers of neurons (MAP2A, green), astrocytes (GFAP, green), and microglia/macrophages (IBA1, green), while nuclei were stained with DAPI (blue). PKH26 dye alone served as a negative control for sEV staining. As expected, labeled sEVs (red) were taken up by different cells in both spinal cord and DRG (Supplementary Fig 1).

### Serum derived sEVs can increase basal pain threshold in naïve mice

Previous *in vivo* studies have demonstrated that exogenous sEVs from specific cell lines can regulate inflammation and development of pain^13, 25^. Our proteome profiling studies found that sEVs from SNI mice had increased levels of C5a and ICAM-1 compared to naïve control mice. As these proteins are proinflammatory and have known roles in pain^26, 27^, we hypothesized that serum-derived sEVs from donor mice could induce or promote pain hypersensitivity in naïve recipient mice. Thus, we collected sEVs from naïve control or SNI model mice, intrathecally injected naïve mice with a specific sEV prep or PBS control, and measured pain thresholds for up to 7 d post-injection (**Fig. 3a**). All the studies were performed using sEVs purified from the serum of mice 2 weeks post SNI surgery as well as age and sex matched naïve control mice.

**Fig. 3.**
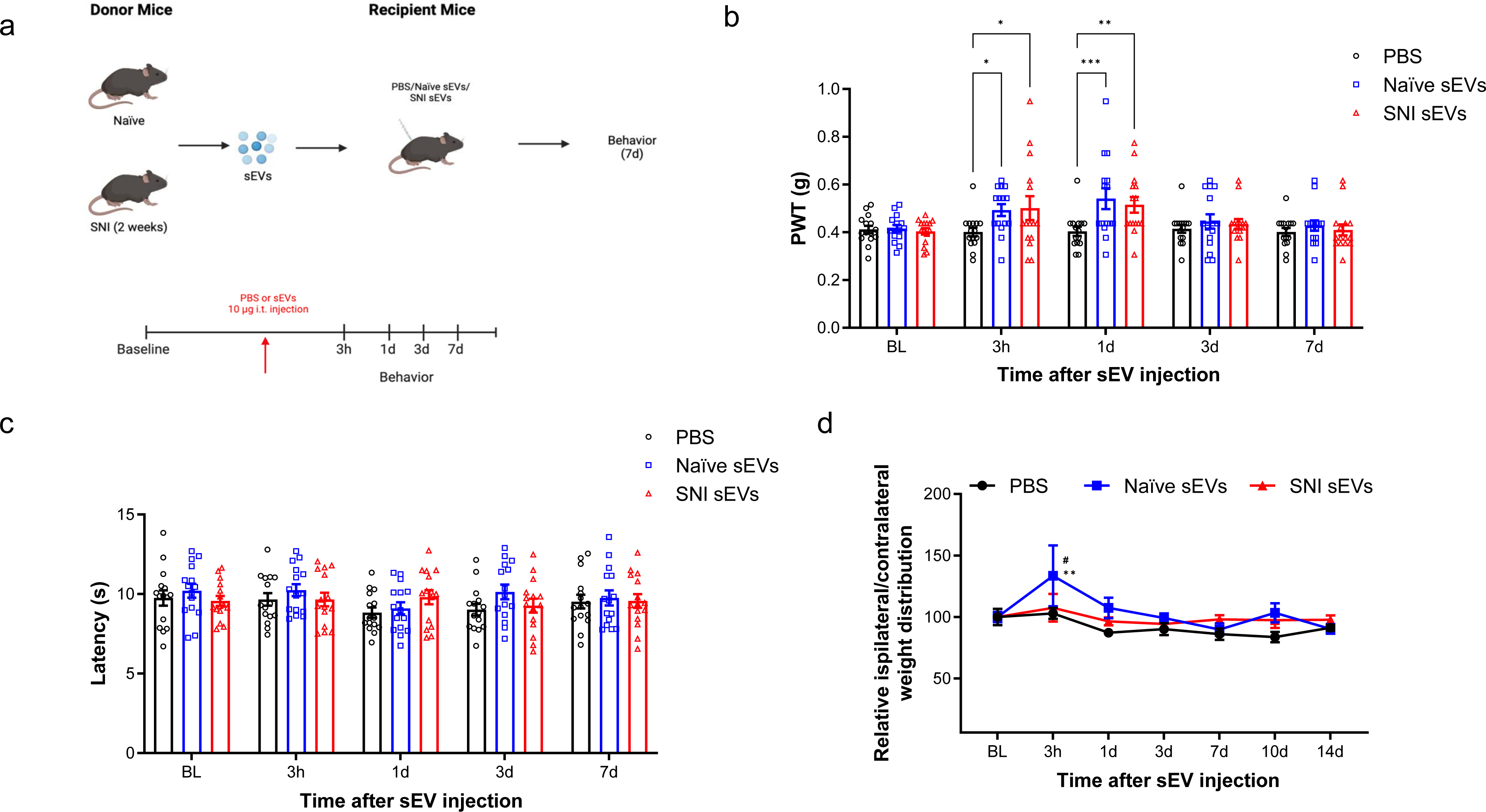
Intrathecal injection of 10 µg sEVs transiently increased basal mechanical but not the thermal threshold. **a** Schematic of *in vivo* experiment. sEVs purified from serum of male C57BL/6 mice 2 weeks post SNI surgery or naïve mice were administrated intrathecally into another group of naïve male recipient mice followed by behavior tests. **b** Mechanical threshold was measured by von Frey filaments before and after the single intrathecal injection of sEVs. The recipient mice showed an increase in basal mechanical threshold (n=15). **c** sEVs did not impact basal thermal pain threshold in mice. Thermal threshold was measured by Hargreaves test before and after sEV injection. The recipient mice showed no significant difference (n=6). Data shown are mean ± SEM. Statistical analysis was determined by two-way repeated-measures ANOVA followed by Bonferroni posttest * p <0.05, ** p <0.01, *** p <0.001. **d** Treatment with sEVs from naïve mice changed basal weight distribution of hind paws in recipient mice. The ratio of body weight distribution on two rear paws (RL/RR) were measured by dynamic weight bearing system at different time points after sEVs injection. Statistical analysis determined by two-way repeated-measures ANOVA followed by Bonferroni posttest ** p <0.01 PBS vs. naïve sEVs, # p<0.05 naïve sEVs vs SNI sEVs (n=5).

Dose response studies showed that low dose (1 µg) sEVs did not change the basal mechanical pain threshold (Supplementary Fig 2), but a higher 10 µg dose surprisingly increased this threshold. Specifically, 10 µg of naïve or SNI sEVs from male donor mice transiently increased basal mechanical threshold at 3 h and 1 day post injection (**Fig. 3b**) in male recipient mice. However, 10 µg sEVs did not alter thermal sensitivity (**Fig. 3c**). There were also no significant changes in female mice injected with sEVs from female SNI and naïve donors (Supplementary Fig 3). Male mice intrathecally injected with sEVs from naïve male mouse donors also showed changes in basal weight distribution of hind paws in a dynamic weight bearing assessment (**Fig. 3d**). Though these results may be influenced by the proinflammatory sEV cargo including ICAM1 and C5a, our behavior results on mechanical threshold partially disproved the hypothesis that serum-derived sEVs could induce/promote pain hypersensitivity. In fact, the sEV-mediated increase in mechanical threshold suggests that serum sEVs may transiently modulate basal pain thresholds. This led us to consider another hypothesis that sEVs transient effects on basal thresholds could be due to their endogenous opioid peptide cargo.

### Detection of endogenous opioid cargo in serum-derived sEVs

We assessed the presence of a subset of opioid peptides with LC-MS/MS analysis focusing on predominant endogenous opioids that bind μ, δ, and κ opioid receptors. We evaluated β-endorphin, met-enkephalin, leu-enkephalin, nociceptin, endomorphin-1, endomorphin-2 and dynorphin A 1-17 in sEVs from serum of both naïve and SNI model mice. Leu-enkephalin, a pentapeptide (Tyr-Gly-Gly-Phe-Leu or YGGFL) was identified in all sEVs by observing fragment peaks of 397 m/z and 425 m/z as previously reported^28^ (**Fig 4a**). We also observed low level of the characteristic fragment peaks of nociceptin, for which the peak intensity was close to background level (Supplementary Fig 4). Additional studies are needed to determine the predominant products from nociceptin. Eluent samples collected from sEVs isolation were also assessed by LC-MS/MS, which identified leu-enkephalin and nociception (data not shown). This observation indicated leu-enkephalin and nociceptin are also present in free form in the serum. The presence of leu-enkephalin in sEVs was orthogonally validated using a competitive enzyme immunoassay (**Fig 4b**) showing leu-enkephalin was enriched in sEVs compared to the serum levels. We also observed that leu-enkephalin in sEVs from the serum of female mice was higher than that of male mice.

**Fig. 4.**
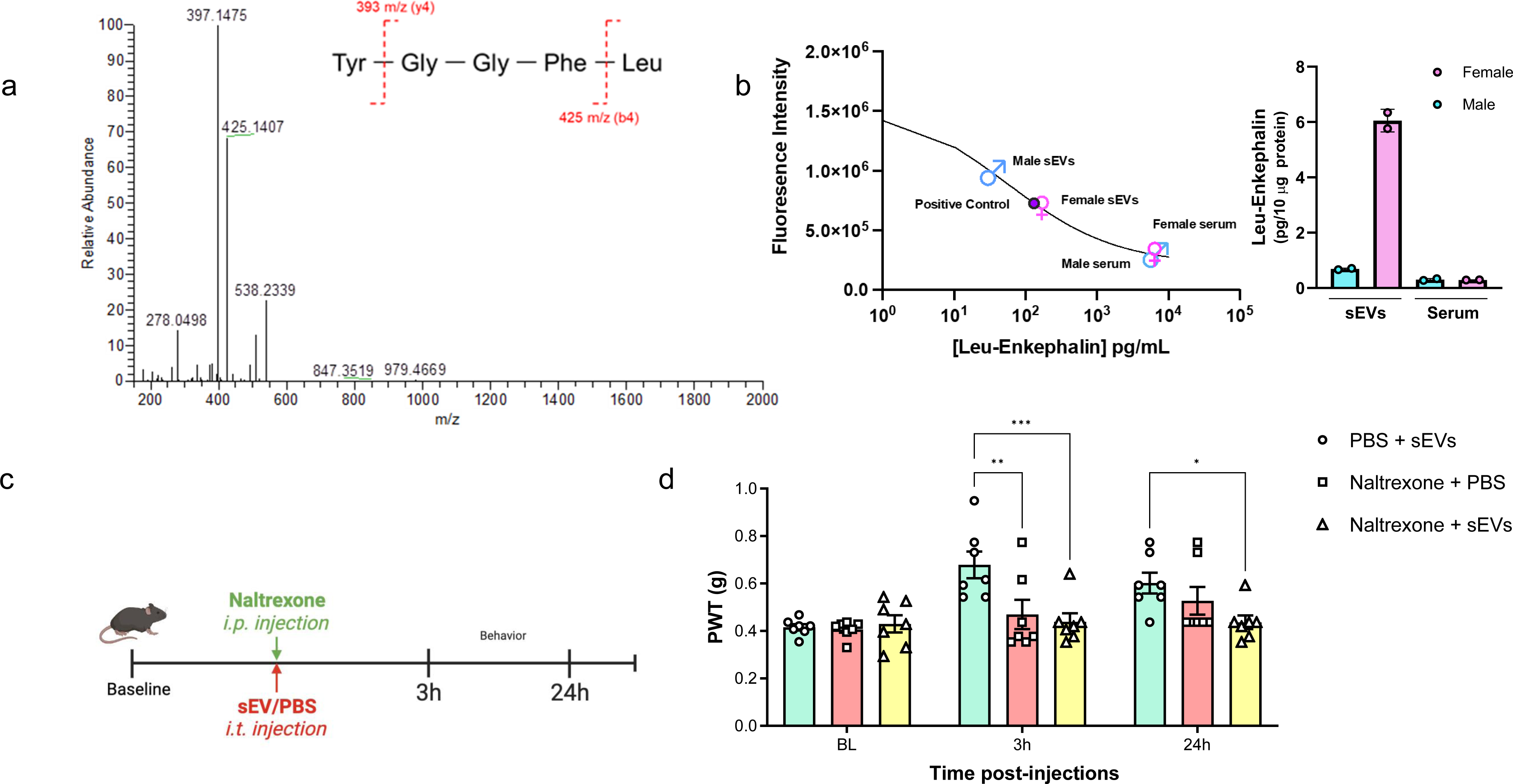
Detection of leu-enkephalin in sEVs and reversal of short-term analgesia induced by naïve sEVs using naltrexone. **a** Liquid chromatography with tandem mass spectrometry of serum sEVs samples. Representative MS/MS spectrum showing the peaks of mass-to-charge ratios (m/z) at 397 and 425 derived from the collision-induced dissociation of leu-enkephalin in sEVs. **b** Confirmation of leu-enkephalin in sEVs by enzyme immunoassay. Levels of leu-enkephalin in serum-derived sEVs and serum from naïve male and female mice. Both sEV and serum samples were pooled from n=5 mice of each sex. The standard curve for leu-enkephalin is also shown. **c** Schematic of *in vivo* experiment in mice administrated with naltrexone (i.p., 10mg/kg) and naïve sEVs (i.t., 10 µg/mice) at the same time, and behavior testing 3h and 24h post-administration. **D** The group receiving naïve sEVs and PBS had higher mechanical threshold than the group receiving both naltrexone and sEVs at 3h post-injection. Similarly, for the 24h time point, the group receiving sEV and naltrexone had higher PWT compared to the group injected with sEV and PBS (n=7). Data shown are mean ± standard error of the mean. Statistical analysis was determined by two-way repeated-measures ANOVA followed by Bonferroni test * p <0.05, ** p <0.01, *** p <0.001.

### Endogenous opioid cargo in sEVs contributes to short-term analgesia

Based on our observation that leu-enkephalin is present in sEVs, we investigated whether non-selective opioid receptor antagonist naltrexone can reverse sEV-induced analgesia. Before initiating the studies using sEVs, we confirmed that naltrexone blocked morphine analgesia at 3h and 24h post-treatment, the two time points that were significant in our sEV experiments. As shown in the schematic of our *in vivo* experiment (Supplementary Fig 5a) naltrexone (10 mg/kg) was administered intraperitoneally into naïve 9-week-old male mice, and after 2h 30min or 24h post injection, 0.5 µg morphine was injected intrathecally. Behavior studies were performed 30 min after morphine injection. As shown in Supplementary Fig 5a, naltrexone itself did not affect the pain threshold and the group that received only morphine showed an increase in basal mechanical threshold. However, naltrexone reversed morphine-induced short-term analgesic effect post 3h naltrexone injection. For 24h time point, we injected naltrexone on day 1, then conducted behavior testing on day 2, before and after morphine injection. As shown in Supplementary Fig 5b, pretreatment with naltrexone decreased morphine analgesia even at 24h post naltrexone injection. After confirmation of naltrexone efficacy, we investigated whether naltrexone could reverse sEV-induced short-term analgesia. We only used sEV from naïve mice and not SNI model mice since both contained leu-enkephalin based on LC-MS-MS studies and sEVs from both sources had very similar short-term analgesic effect. We injected mice with naltrexone (10 mg/kg) and sEVs (10 µg/mice) at the same time and tested pain thresholds 3h and 24h post injection (**Fig. 4c**). At 3 h post injection, the group that received sEVs and PBS had increased mechanical threshold, but naltrexone reversed this effect (**Fig. 4d**). For the 24h time point, the group that received sEVs and naltrexone had lower sensory threshold compared to the group injected with sEVs and PBS. This suggests naltrexone reversed analgesia conferred by sEVs. Collectively, our studies showed that leu-enkephalin is present in naïve and SNI sEVs and naltrexone reversed analgesic effect of sEVs obtained from the serum of naïve mice.

### Serum derived sEVs have a long-term prophylactic role in the resolution of chronic pain

We next investigated if sEVs had long term effects on pain threshold under chronic pain states in addition to their short-term effects. Since we used the term “short-term” for a time window of less than 48 h earlier, we will use “long-term” for observations made two weeks post sEV injection. We chose SNI mouse model of chronic neuropathic pain and CFA mouse model of persistent inflammatory pain for *in vivo* studies.

### The effect of serum-derived sEVs on recipient SNI model mice

First, we investigated how sEVs from both naïve control mice and SNI model mice control pain in SNI model recipient mice. Recipient mice were injected intrathecally with sEVs or PBS two weeks prior to SNI surgery. After surgery, they underwent behavioral testing for 21 days (**Fig. 5a**). A 10 µg dose of naïve or SNI sEVs increased the mechanical threshold 1- and 3-days post-surgery, compared to PBS control (**Fig. 5b**). Pretreatment with sEVs did not prevent mechanical hypersensitivity in the SNI model, but both sEV types slightly delayed the development of SNI-induced pain hypersensitivity in the initial stages. However, there were no differences after the third day. There was also no significant difference for thermal threshold between groups (**Fig. 5c**).

**Fig. 5.**
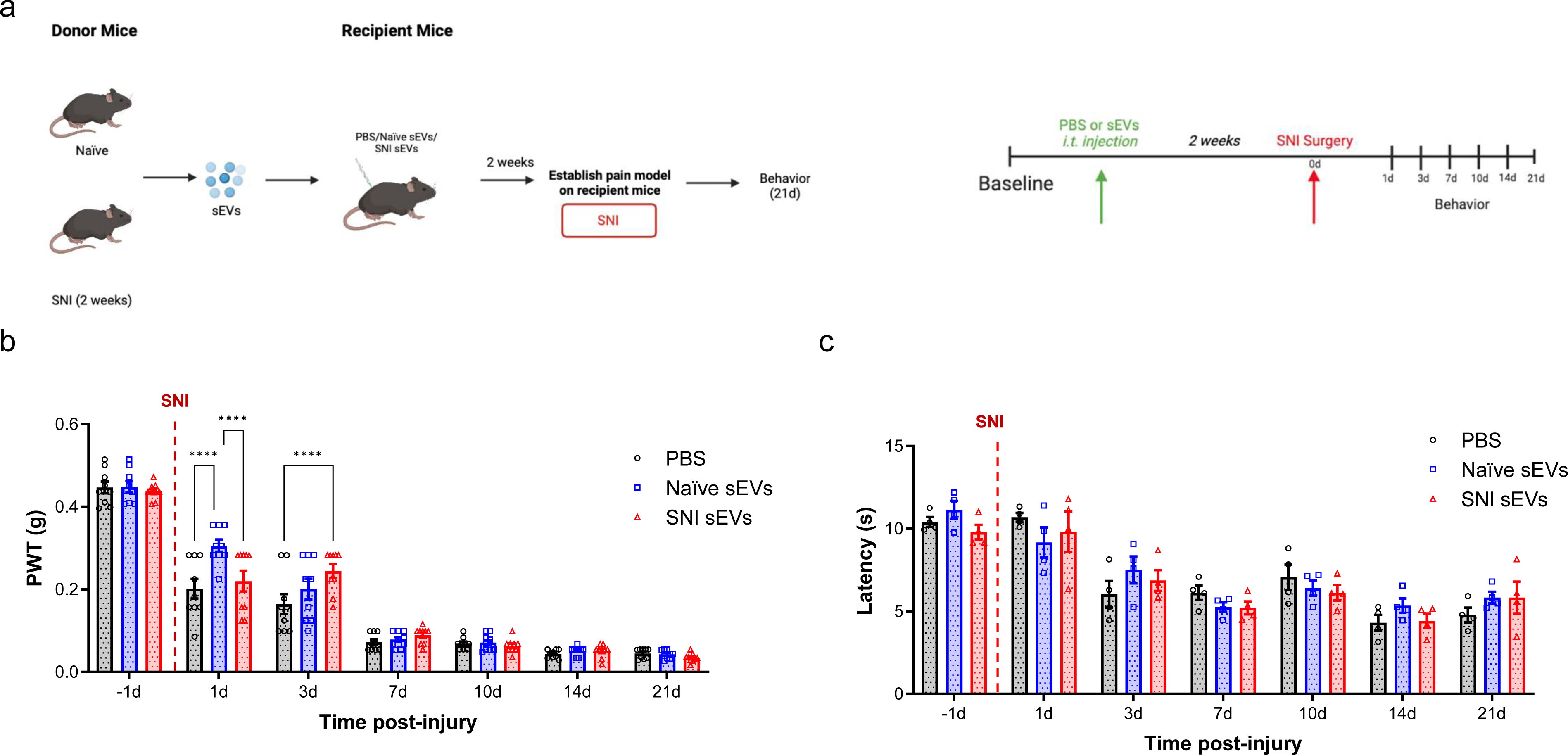
The pretreatment with 10 µg of serum-derived sEVs caused a transient delay in the development of mechanical allodynia in SNI model. **a** Schematic of experiment showing the serum-derived sEVs from naïve and SNI model donor mice two weeks after injury injected intrathecally into another group of naïve recipient mice. After two weeks, the SNI surgery was performed on these recipient mice followed by behavior tests to assess the effect of sEVs on SNI-induced pain. **b** Mechanical hypersensitivity measured by von Frey filaments. The recipient mice that received 10 µg naïve or SNI sEVs then underwent SNI surgery showed increased mechanical threshold at 1d and 3d compared to PBS control (n=9). These results indicate sEVs from both naïve and SNI model can induce a transient delay in SNI-induced mechanical hypersensitivity in the initial stages. **c** There was no significant differences in thermal hypersensitivity as measured by Hargraves after SNI surgery. Data shown are mean ± SWM. Statistical analysis was determined by two-way repeated-measures ANOVA followed by Bonferroni test, **** p < 0.0001.

### The effect of serum-derived sEVs on recipient CFA model mice of inflammatory pain

We next investigated if sEVs could attenuate CFA induced inflammatory pain. We previously found that an intrathecal injection of 1 µg sEVs from RAW 264.7 macrophage cells reversed mechanical allodynia induced by CFA^13^. For this study, we performed a dose response by injecting 1 and 10 µg serum-derived sEVs into recipient male mice two weeks before injecting them with CFA (**Fig. 6a**). While 1 µg sEVs did not produce any significant differences (Supplementary Fig 6), 10 µg sEVs significantly decreased CFA-induced mechanical hypersensitivity after 14 days (**Fig. 6b**). Multiple comparison between different time points shows that pretreatment with sEVs from either naïve or SNI model mice accelerated recovery i.e., reached baseline for mechanical threshold faster than mice that received PBS injection (**Fig. 6b**). Specifically, mice that received sEVs returned to the baseline threshold 14 days after CFA injection, but the control group injected with PBS required 21 days for recovery. These results show that 10 µg serum-derived sEVs have a prophylactic effect that can enhance recovery in CFA model. sEVs did not affect thermal sensitivity in male mice (**Fig. 6c**), and we also did not observe significant changes in comparable studies in female mice (Supplementary Fig 7).

**Fig. 6.**
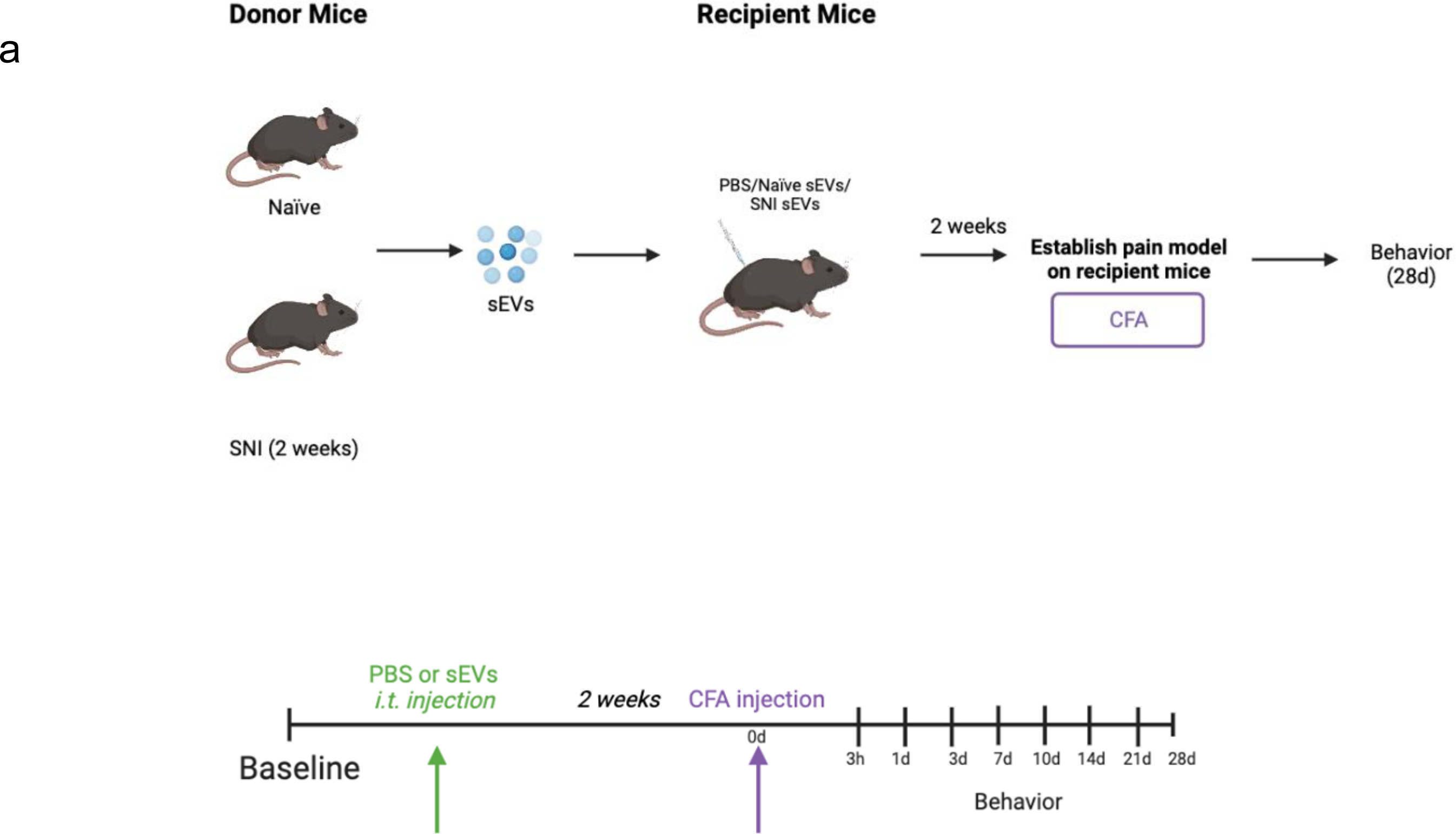
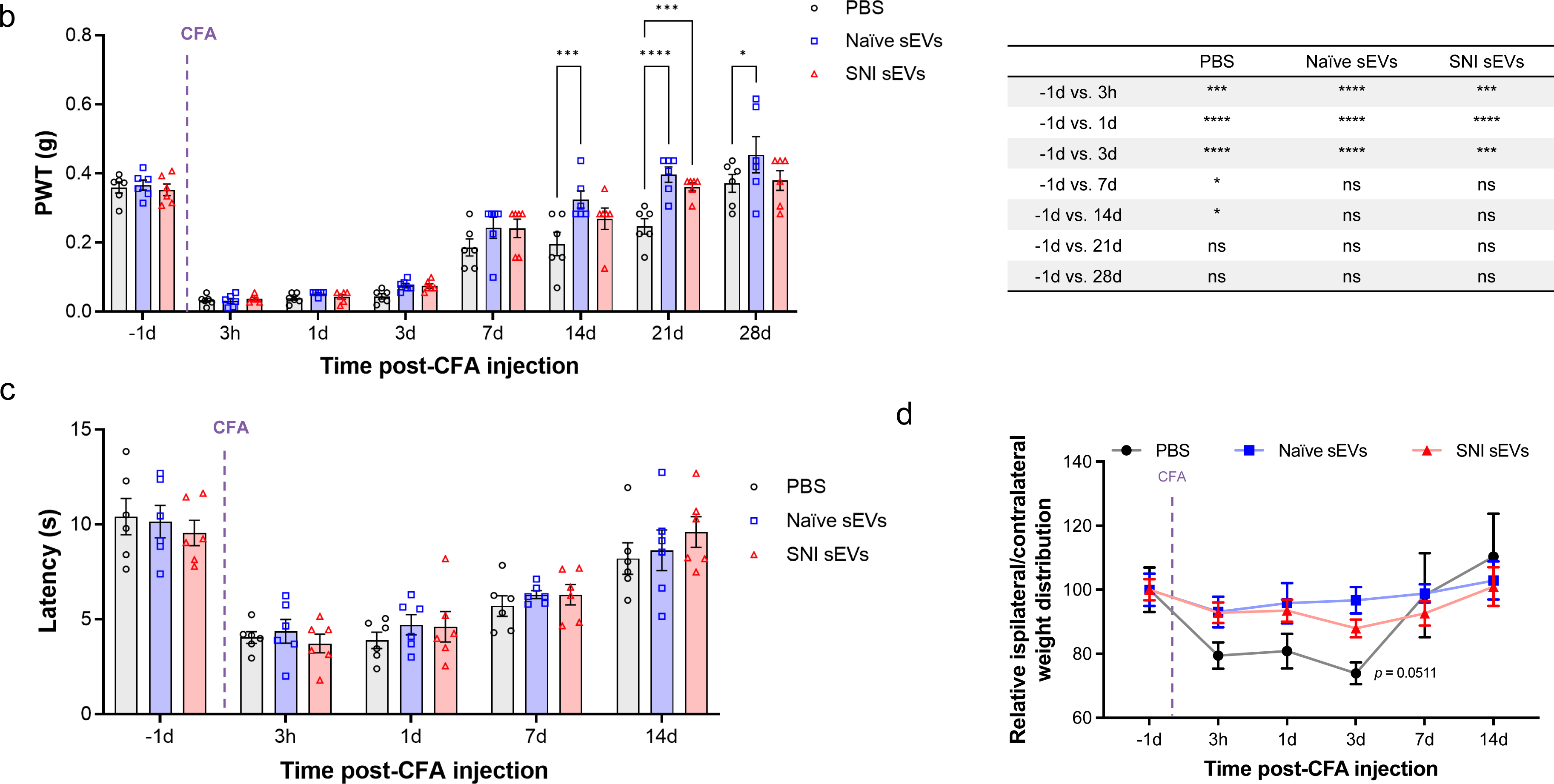
Prophylactic intrathecal injections of 10 µg sEVs promoted resolution of CFA induced mechanical hypersensitivity. **a** Schematic representation of experimental design. The sEVs from male C57BL/6 naïve and SNI model donor mice two weeks after injury were injected intrathecally into nine weeks old male C57BL/6 recipient mice. CFA model was established in these recipient mice 2 weeks post sEVs injection. **b** Mechanical allodynia in CFA model mice. Mice were prophylactically injected with 10 µg sEVs or PBS two weeks prior to CFA. PWT were measured at different time points after CFA injection (n=6). Ten µg sEV accelerated recovery from CFA-induced mechanical hypersensitivity compared with PBS control group. Multiple comparisons between different time points in CFA model are shown on the right. **c** There were no significant differences in thermal hypersensitivity as measured by Hargreaves test. Data shown are mean ± SEM. Multiple comparation between different time points in CFA model is shown in adjacent table. Statistical analysis was determined by two-way repeated-measures ANOVA followed by Bonferroni test * p <0.05, *** p < 0.001, ### p < 0.001, **** p < 0.0001. **d** Pretreatment with naive sEVs reverse CFA-induced weight distribution changes on rear paws. sEVs from male C57BL/6 naïve mice were injected intrathecally into nine weeks old male C57BL/6 recipient mice. CFA was injected into rear right (RR) paw of these recipient mice 2 weeks post sEV injection. The ratio of body weight distribution on two rear paws (RL/RR) were measured by dynamic weight bearing system at different time points after CFA injection (n=5). Ten µg naïve sEV significantly reversed CFA-induced increasing ratio of body weight distribution (RL/RR) on paws after three days compared to PBS control group. Though sEVs from SNI model had a similar trend, it was not significant. The significance was observed on day 3 between SNI sEVs and PBS control. Data shown are mean ± SEM. Statistical analysis was determined by two-way repeated-measures ANOVA followed by Bonferroni posttest * p <0.05.

### The effect of serum-derived sEVs on spontaneous pain

Since both von Frey and Hargraves tests are evoked measures of pain hypersensitivity, we evaluated spontaneous pain in CFA model mice with a DWB system that measures how they distribute weight on all four paws. Previous studies have reported CFA injection will change weight distribution on rear paws compared with control^29, 30^. We observed reduction in ipsilateral hind paw weight bearing after CFA injection (**Fig. 6d**), but the groups pretreated with serum sEVs from either naïve or SNI model mice displayed a much smaller change of weight distribution. The relative ratio of weight between left and right paw in naïve sEV-treated group was lower than control group at 3 days post CFA injection. The ratios of all groups returned to baseline after 7 days post CFA. This suggests sEVs from naïve mice can reverse CFA-induced weight distribution changes on hind paws.

### Immune cell alterations in spinal cord and DRG in sEV treated CFA model recipient mice

We next used ChipCytometry to investigate changes in immune cell populations in the spinal cord and DRG of CFA model mice treated with sEVs or PBS. At 7- and 14-days post CFA treatment, we collected tissue samples from the PBS treated control group and from mice injected with sEVs from naïve or SNI model donor mice. We also included a group treated with sEVs from RAW 264.7 macrophage cells for comparative studies. For the quantification of cellular phenotypes flow cytometry-like hierarchical gating strategy was followed until all population gates were quantified^31^. We used a custom algorithm to quantify cell populations and markers in individual sections (two sections from two mice/group). Cells were normalized to the corresponding tissue surface area, and several immune cell subtypes showed low or no expression in the regions analyzed.

**Fig. 7a** shows an example of image overlay in SC representing how NK cells were identified by a combination of markers. As expected, the immune cells probed comprised of only a subset of all cells detected. **Fig. 7b** shows the number of immune cells 14 days post CFA and **Fig.7c** shows the percentage of CD45^+^ cells. The bar graphs (**Fig. 7d-l**) show the mean ± SE of total and various immune cells. ChipCytometry enabled us to compare the differences between CFA injected ipsilateral side of the spinal cord to contralateral. For better visualization of how sEV treatment affected immune cells, we used pie charts of cell type distributions (**Fig. 7m-o**). CFA model mice displayed a much larger portion of NK and NKT cells in spinal cord, and the proportion was much larger on the ipsilateral side. However, sEV treatment completely reversed these outcomes. In mice that received RAW 264.7 sEVs, neutrophils increased in ipsilateral side.

**Fig. 7.**
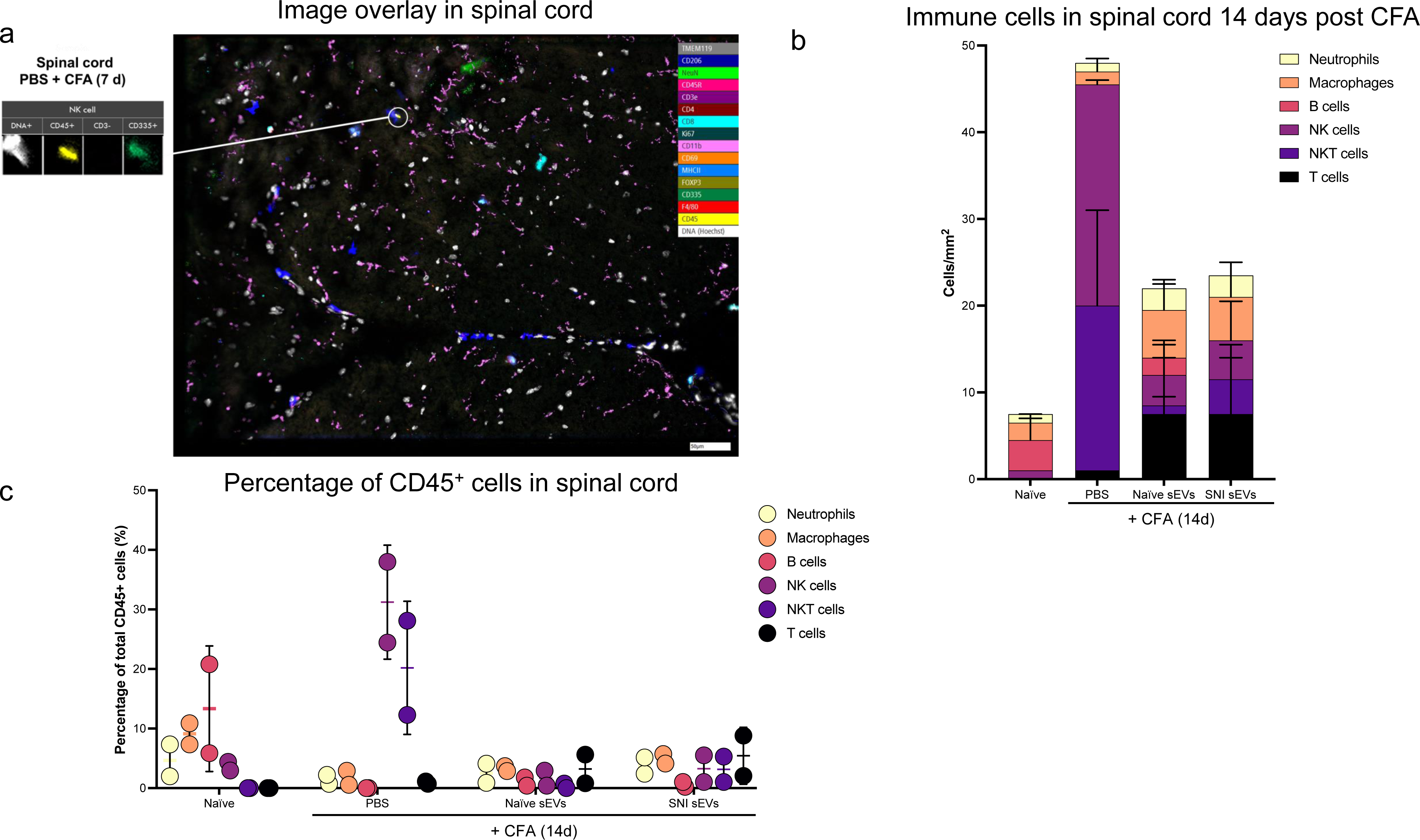
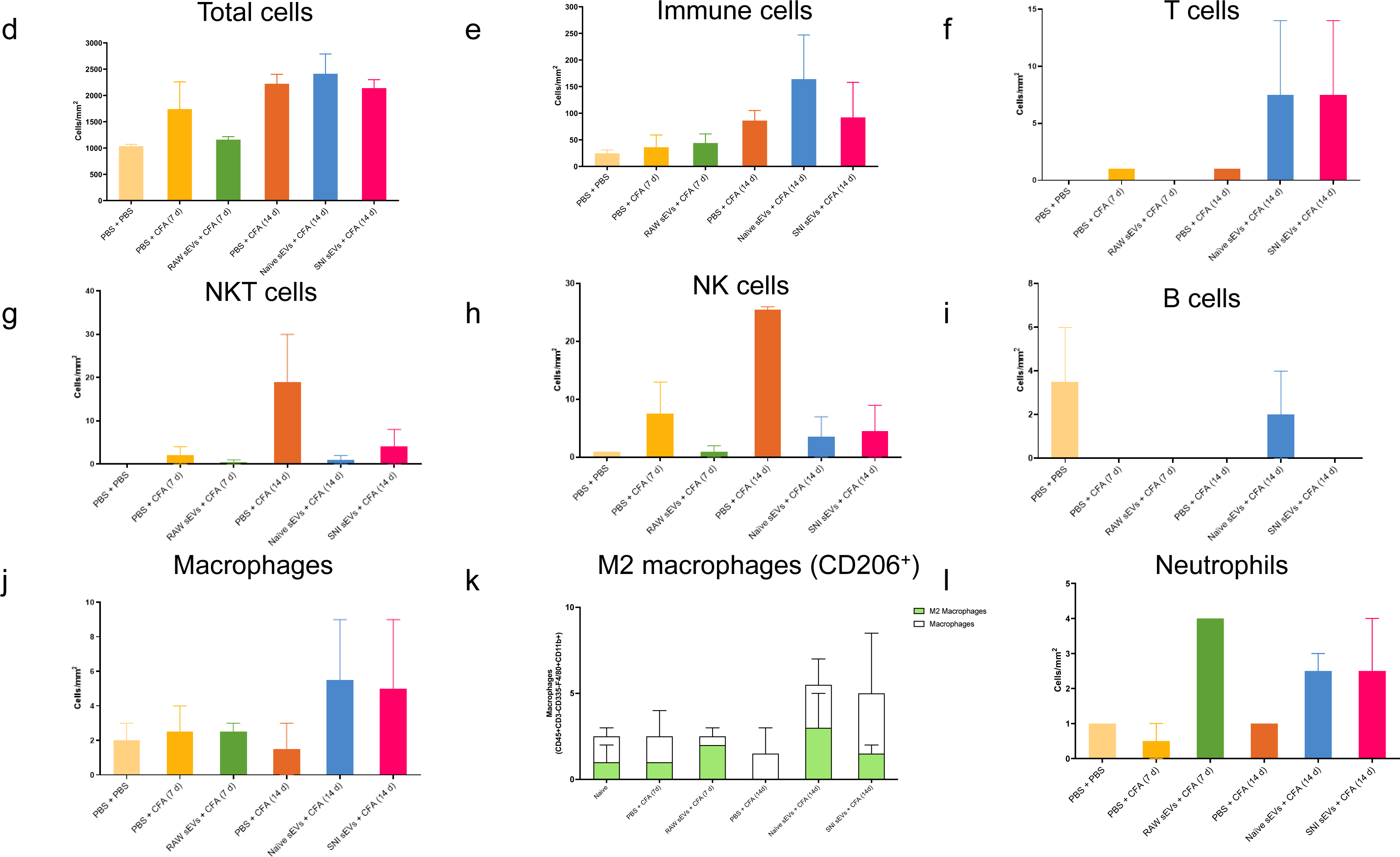
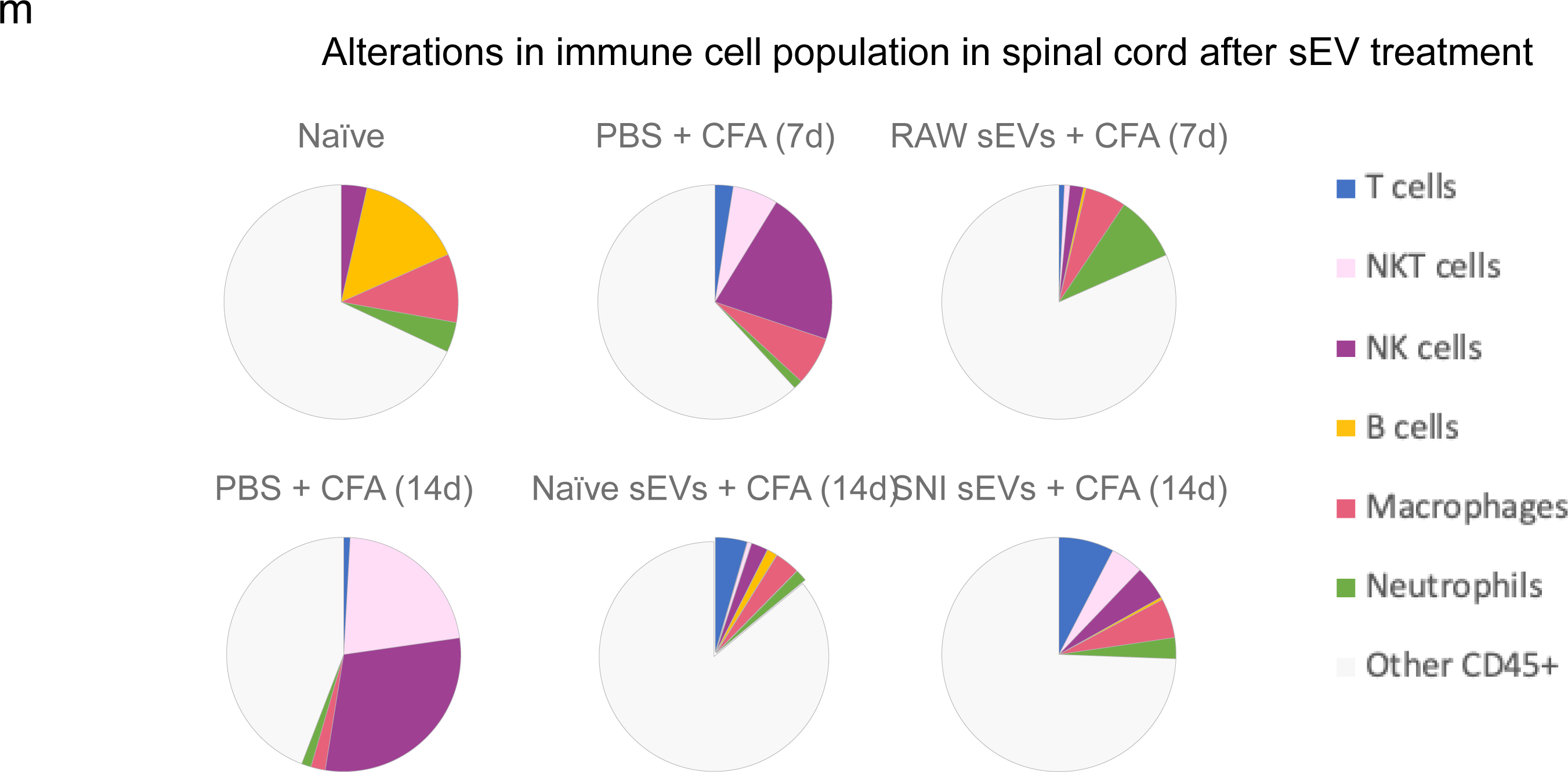
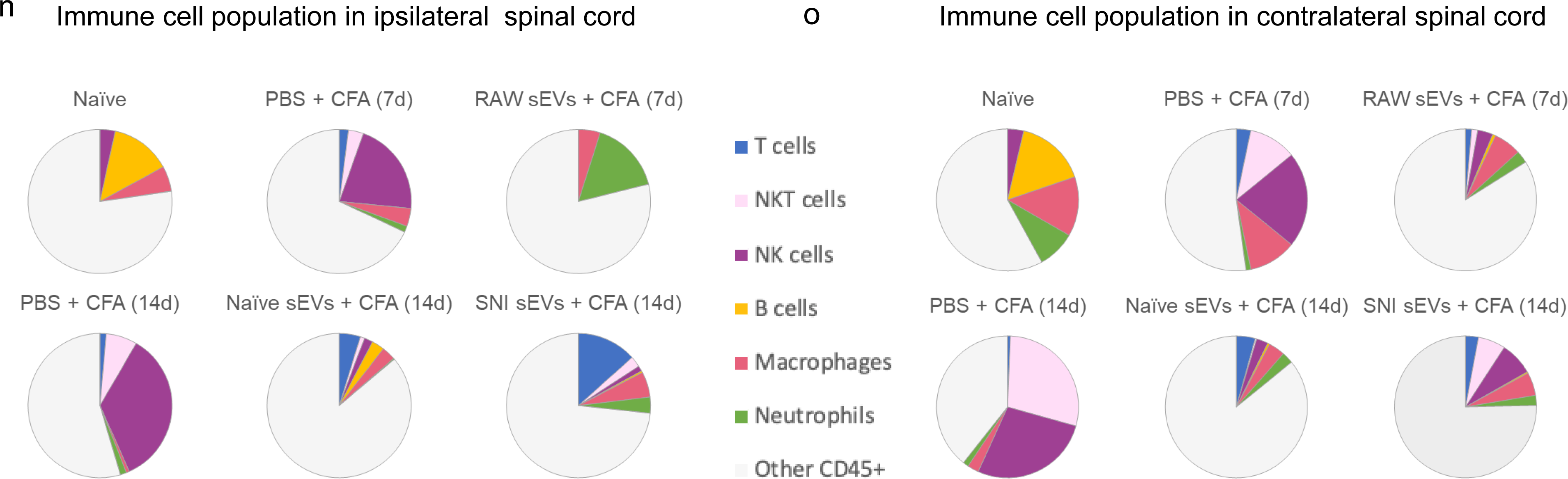
Alterations in immune cell population in 5-7 µM sections spinal cord after sEV treatment determined using ChipCytometry. **a** An example of image overlay in spinal cord. **b** Immune cells in spinal cord 14 days post CFA **c** Percentage of CD45^+^ cells in spinal cord. Bar graphs showing **d** total cells **e** Immune cells **f** T cells **g** NKT cells **h** NK cells **I** B cells **j** macrophages **k** M2 macrophages (CD206^+^) **l** Neutrophils Pie chart representation of alterations in immune cell population after sEV treatment in **m** the whole **n** in ipsilateral **o** in contralateral spinal cord. Experimental groups are naïve control, 7 days post CFA, 7 days post CFA mice + sEVs from RAW 264.7 macrophage cells, 14 days post CFA, 14 days post CFA + sEVs from naïve donor mice, 14 days post CFA + sEVs from SNI model n=2.

We analyzed the DRG samples similarly (**Fig. 8a-b**) and found different immune cell compositions. F**ig. 8c** shows the percentage of CD45^+^ cells and the bar graphs representing different cells are shown in **Fig. 8d-l**. As sEV treatment increased B cell numbers, we examined additional B cell and myeloid markers, CD19 and Gr1 respectively, using the re-probing feature of ChipCytometry. CD19 staining was weak in all samples and most Gr1 signal did not colocalize with CD45, suggesting most Gr1^+^ cells were not neutrophils. Additionally, CFA treated samples showed lower numbers of macrophages at 14 days post-CFA compared to 7 days, and macrophage numbers were increased by both naïve and SNI sEV treatment. We further gated on an M2 or anti-inflammatory macrophage marker CD206^+^ to identify the macrophage subpopulations. M2 macrophages are shown in green (**Fig. 8k**) and neither M2 nor total macrophages were readily detected in DRG 14 days post CFA. However, almost all the macrophages detected in mice treated with sEVs from RAW 264.7 cells were M2. Overall, administration of sEVs increased the M2 population. **Fig. 8m** shows CD4/CD8 over all T cells. **Fig. 8n** is the pie chart representation of alterations in immune cell population after sEV treatment in DRG.

**Fig. 8.**
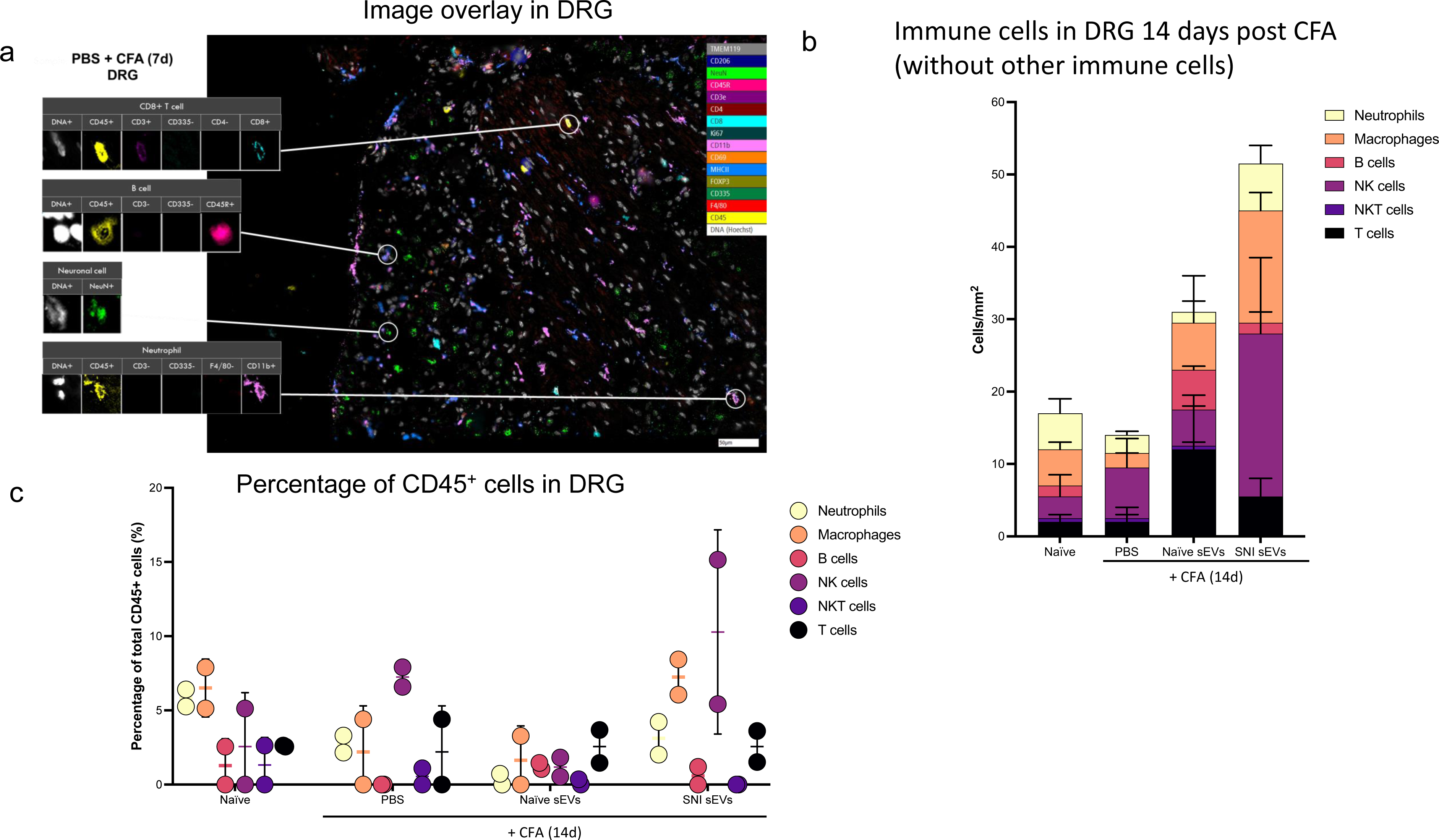
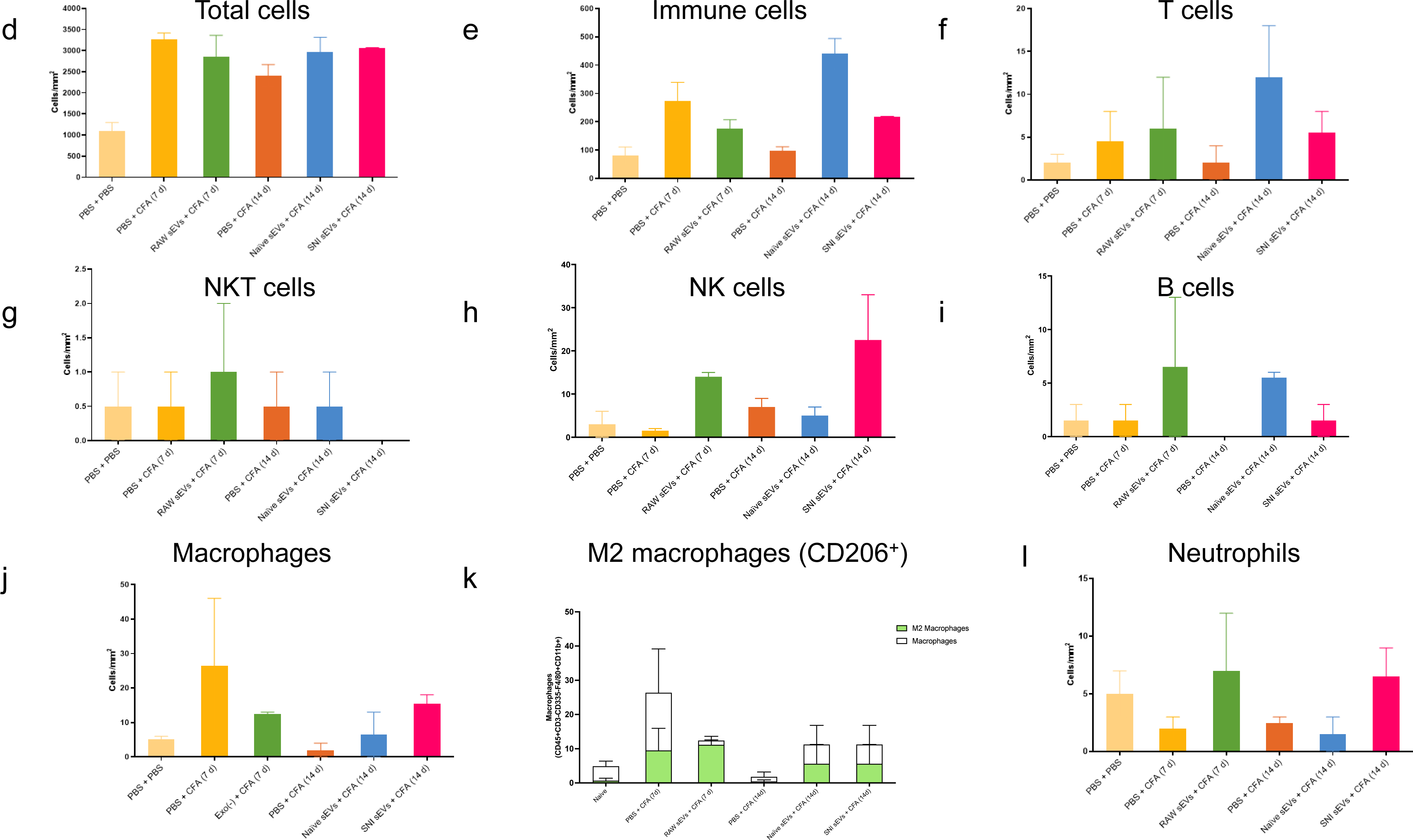
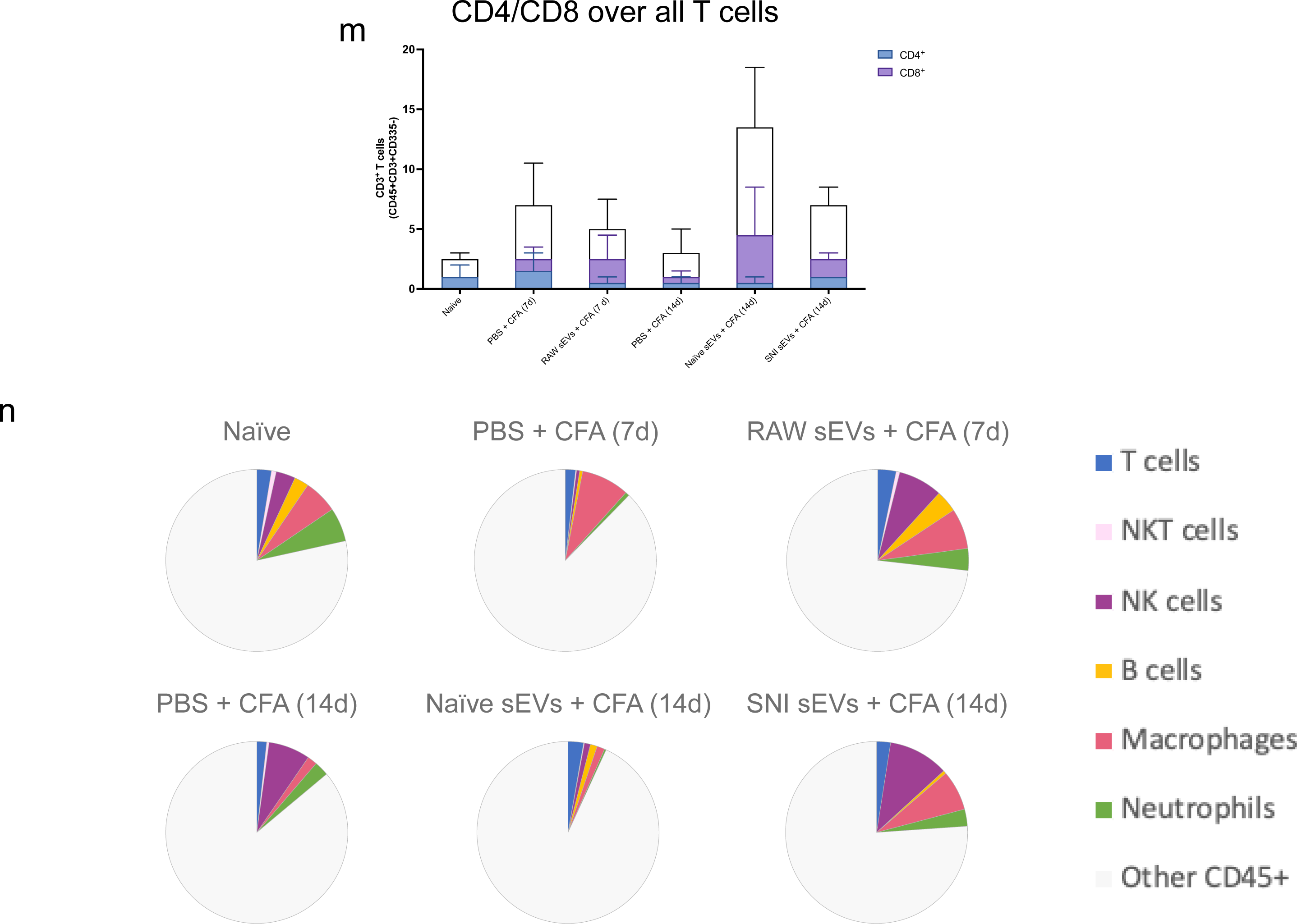
Alterations in immune cell population in 5-7 µm sections of DRG after sEV treatment determined using ChipCytometry. **a** An example sample image overlay in DRG. **b** CD45^+^ immune cells in DRG 14 days post CFA with additional specified markers in the immune pane**. c** Percentage of CD45^+^ cells in DRG. Bar graphs showing **d** total cells **e** Immune cells **f** T cells **g** NKT cells **h** NK cells **I** B cells **j** macrophages **k** M2 macrophages (CD206^+^) **l** Neutrophils **m** CD4/CD8 over all T cells. **n** Pie chart representation of alterations in immune cell population after sEV treatment in DRG n=2.

### Characterization of sEVs for immune markers

sEVs released by specific immune cells encapsulate properties of the donor cells releasing them. For example, dendritic cells (DCs) and exosomes released by DCs have similar membrane proteins and biological properties^32^. We investigated if some of the same immune cell markers were enriched in sEVs. The ExoView platform enables high-resolution multidimensional characterization of EVs combining size, count, and protein expression measurements at the level of single vesicle. EVs expressing specific proteins bind to the corresponding antibodies on the ExoView chip. ExoView analyzes EVs using visible light interference for size measurements and fluorescence for protein profiling. We used the ExoView Exoflexprotein microarray chip which has antibodies against commonly expressed EV tetraspanin proteins CD9 and CD81, and allowed for custom capture antibodies to be added to the chip, CD45 and F4/80, hematopoietic and macrophage markers, respectively. Since sEVs from macrophages resolved CFA-induced inflammatory pain faster than vehicle^13^, we investigated if these sEVs express macrophage markers and designed a custom assay using CD45, F4/80 to specifically capture the subpopulation of sEVs derived from macrophage. We also used custom detector antibodies against these same markers to evaluate distributions of these markers on sEVs captured by anti-CD9, CD81, CD45, or F4/80. Capture antibodies specific to EV proteins immobilize EVs on the chip and we confirmed that the sEVs from mouse serum express these markers (**Fig. 9a**). **Fig. 9c** and **9d** show that serum-derived sEVs not only carry the exosome tetraspanin marker proteins but a subset of sEVs also express macrophage/monocytic markers, suggesting these sEVs could be one of the contributing factors that helps attenuate inflammatory hypersensitivity. The particle sizes for naïve and SNI model mice serum were below 100 nm (**Fig. 9b**). Representative images are shown **in Fig. 9e**. All chips also include appropriate isotype negative controls to test for non-specific binding (Supplementary Fig. 9a-b).

**Fig. 9.**
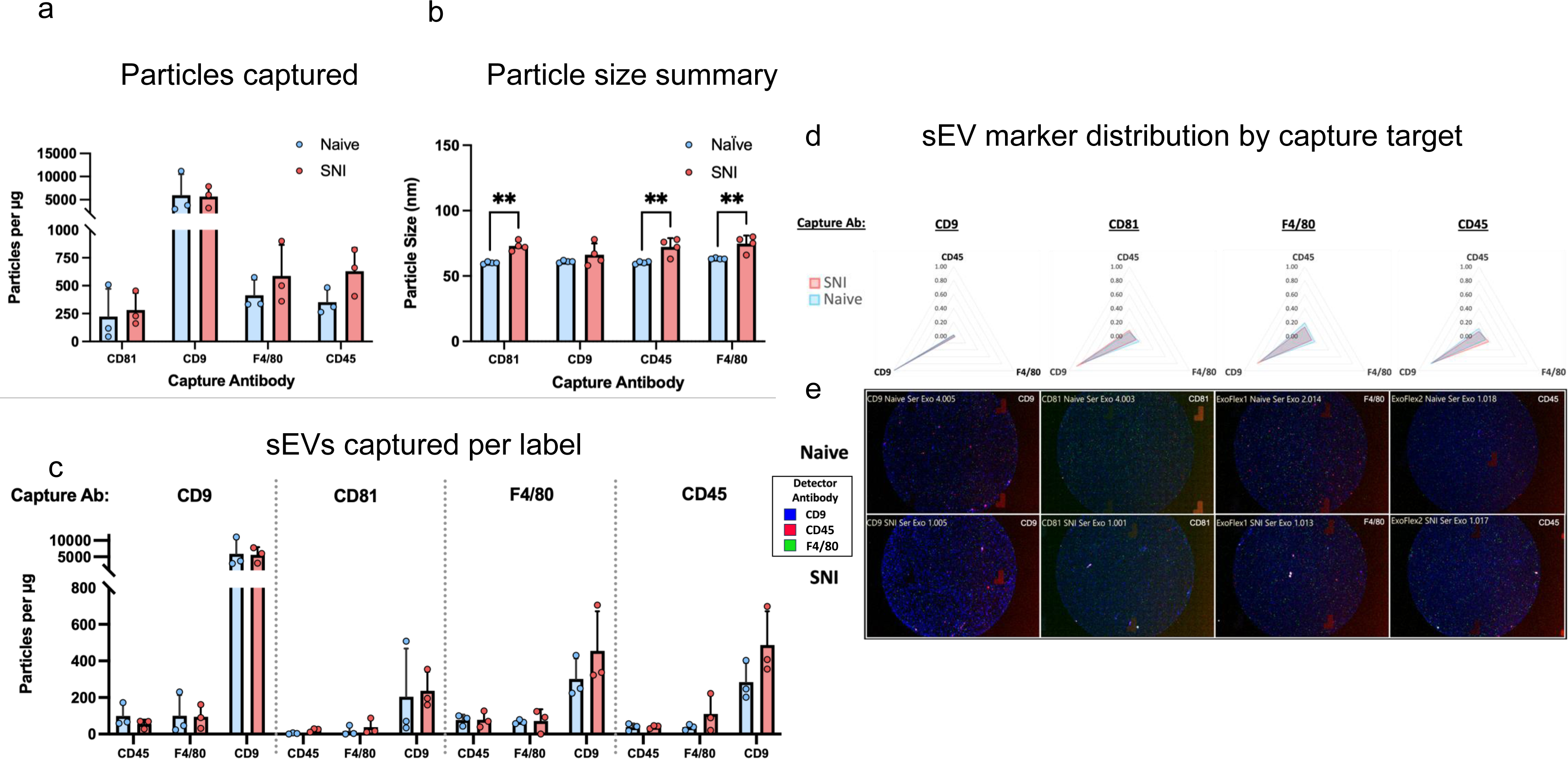
Comparison of tetraspanin composition, leukocyte and macrophage markers and characteristics of sEV derived from naïve and SNI model mice. SEVs were captured using specific antibody-coated spots against CD81, CD9, F4/80 and CD45 and analyzed using the ExoView R100 platform. **a** Total fluorescent particle counts of sEVs on tetraspanin capture spots and mouse IgG isotype control analyzed using fluorescent antibodies against CD81/CD9/F4/80 and CD45. **b** Size distribution of sEVs on tetraspanin capture spots analyzed using the SP-IRIS mode of the ExoView R100 platform. **c** sEVs captured per label. Plotted is the mean ± SEM of three independent biological replicates.

### EV characterization by microflow cytometry

In addition to NTA to describe the hydrodynamic diameter (**Fig. 10a**), additional characterization of sEVs was performed using μFlow. While a modeling algorithm was not used to translate the light scatter intensities to standard units, we sought to provide an estimation of the relative light scatter intensities based on reference beads. In these studies, the light scatter intensity of detected vesicles range was ≥ 83nm polystyrene bead to 480nm polystyrene bead or 1300nm silica bead. Most vesicles were detected in the range of light scatter intensities between 83nm – 110nm polystyrene (**Fig. 10b**). Of note, we are not claiming any standard size because we have not confirmed the refractive index of these sEVs using a standard curve of refractive index oils or other orthogonal measure; in addition, there are now evolving estimates of the presumed refractive index (∼1.35 – >1.42) ^33, 34, 35, 36, 37^ of biological EVs which confounds the presumed sEV sizes using available algorithms. To confirm the vesicles were lipid membrane composed, samples were treated with detergent to confirm solubilization of the vesicles (**Fig. 10c**). We saw that in the presence of Triton, sEVs profiles were dramatically altered, interpreted as a general decrease in relative intensity and concentration of the CFSE positive vesicles when disrupted by the detergent, further confirming their properties as sEVs. Since we use protein concentration for sEV delivery, we determined how protein concentration correlated to particle number. We have previously found that 1 µg sEVs protein equals to 1 x 10^9^ particles according to NTA^13^. Using sEVs from RAW 264.7 cells, we compared the NTA versus µFlow Cytometry plotted with protein concentration and found a good correlation with a similar estimate of 5.89 x 10^8^ particles per µg by NTA (**Fig. 10d**).

**Fig. 10.**
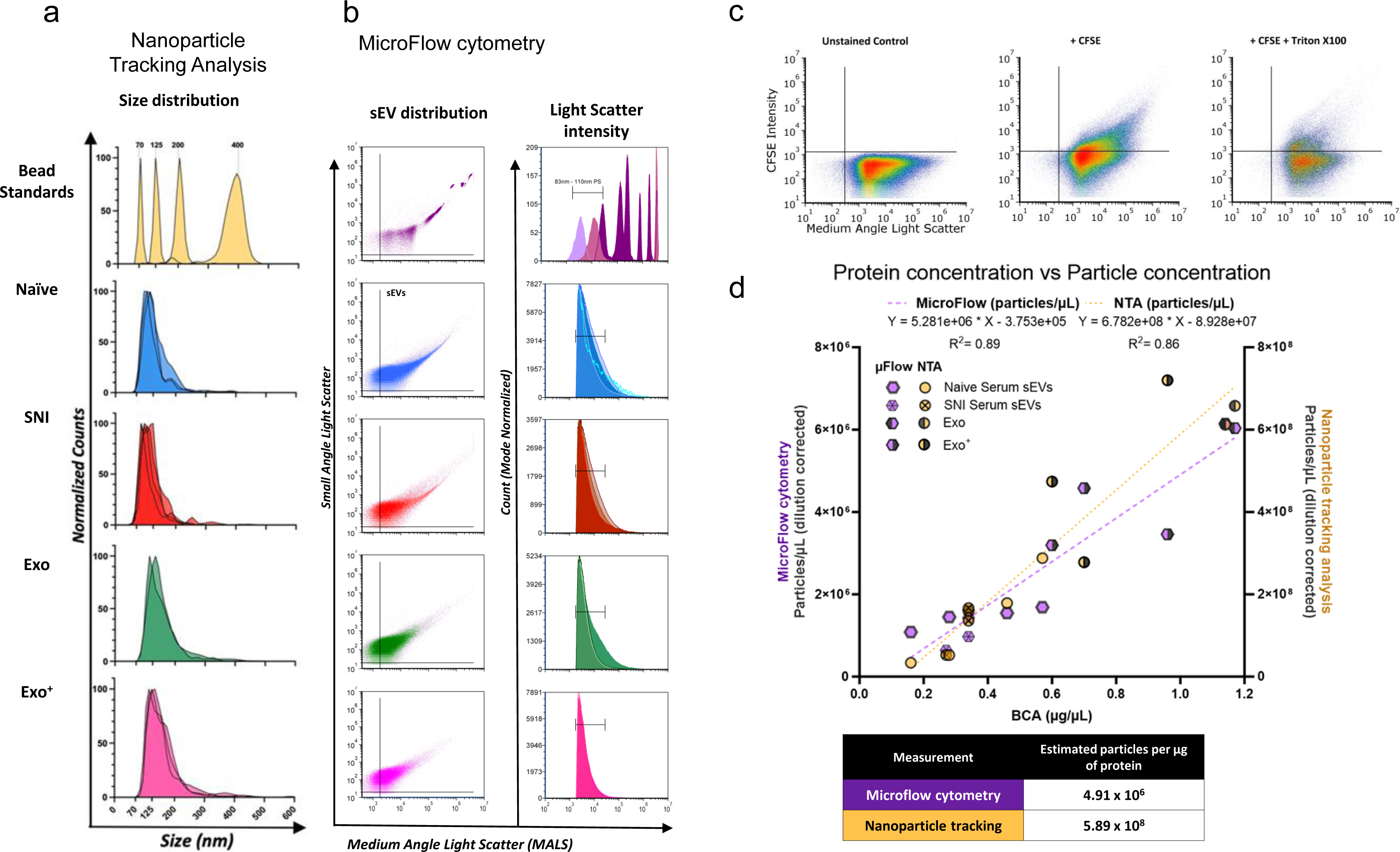
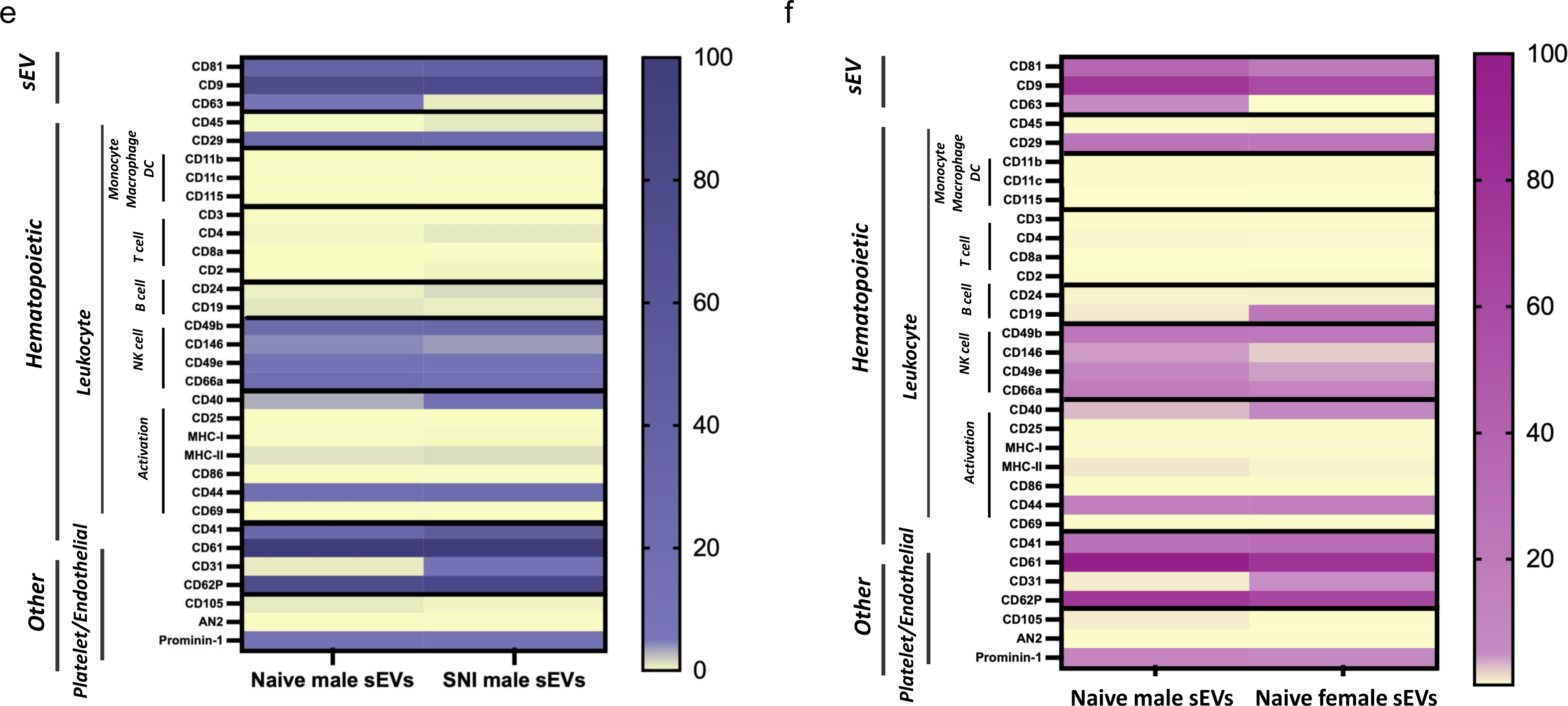
Characterization of serum-derived sEVs and macrophage derived sEVs. **a** Nanoparticle tracking analysis (NTA) size distributions NIST bead standards (yellow), serum-derived sEVs and macrophage derived sEVs (n=2-4 samples per condition). **b** Concatenated microflow cytometry plots of calibration beads (polystyrene-dark purple and silica-light purple), serum derived sEVs (naïve-blue and SNI-red), and RAW 264.7 macrophage derived sEVs from cells with (Exo+ pink) or without (Exo green) LPS stimulation (n=2-4 samples per condition). **c** Representative gating scheme of CFSE positive sEVs for sEVs labeled by CFDA-SE with or without Triton. **d** Plot demonstrating the linear correlation of all sample protein concentrations with either microflow cytometry (purple) or NTA (yellow) particle concentrations. **e** Detection of surface marker profile of sEVs by flow cytometry using MACSPlex exosome capture beads. sEVs from male naïve control and SNI model mice **f** sEVs from male and female naive control mice. The geoMFI (geometric Mean fluorescence Intensity) of different capture antibody bead types after incubation with 15µg EVs, followed by staining with a cocktail of CD9-APC, CD63-APC, and CD81-APC detection antibodies is reported. The geoMFI of individual surface marker was normalized against the highest calculated geoMFI. sEVs used were pooled from serum of 5-6 mice.

### Semi-quantitative analysis of differential surface epitopes on sEVs

Having identified macrophage specific markers on serum-derived sEVs, we further investigated surface expression of a panel of immune markers using exosome capture beads with standard flow cytometry. Enrichment of exosomal marker CD63 in sEVs from naïve male mice and CD4 (T cell), CD24 (B cell), CD40, CD41, MHC-II and (activation markers), and CD31 (endothelial) were observed in sEVs from male SNI model (**Fig. 10e**). Comparison of sEVs from naïve male and female mice (**Fig. 10f**) showed higher CD63 and CD146, CD49e (NK cell), CD105 (endothelial) and MHC-II in male, and CD19 (B cell), CD40 and CD31 in sEVs from naïve female mice. Our data suggests some of the immune markers present in sEVs could impart immunomodulatory functions that may partially mimic the biological properties of their donor cell.

## Discussion

Our studies show that serum-derived sEVs and their cargo molecules have both a short-term analgesic capacity and promote chronic inflammatory pain resolution in recipient mice. Short-term analgesic properties of sEVs were dependent upon opiate signaling, while inflammatory pain resolution was associated with immunomodulatory properties of sEVs through alteration of immune cell populations in the spinal cord and DRG. These findings imply that various sEV cargo molecules and the pain model being investigated in sEV recipient mice can govern the behavioral outcome. Though naïve and SNI model mice differentially package miRNA cargo into sEVs, our behavior studies showed that both sEVs resolved pain comparably. sEV peptide and protein cargo likely act immediately while the miRNA mediated gene regulation could contribute to sEVs long-term effects. Thus, long-term pain resolving effects are likely driven by common miRNAs enriched in all serum-derived sEVs.

Endogenous opioid peptide leu-enkephalin was present in sEVs from both SNI model and naïve mice. Leu-enkephalin can induce short-term analgesia^38, 39^. It was difficult to quantify the peptides by LC-MS-MS since the peak area was too low. However, our analysis clearly detected these peptides in all the sEV samples. We confirmed the presence of leu-enkephalin by a quantitative immunoassay and observed sEVs had elevated levels compared to serum. Leu-enkephalin in sEVs from female mice was higher than male. Enkephalins can activate both mu and delta-opioid receptors but their affinity is 10-fold lower for mu-opioid receptors and negligible for kappa-opioid receptors^40^. In mouse models of inflammatory pain, the analgesic effects of exogenous and endogenous opioids are mediated by mu-opioid receptors expressed in distinct subpopulations of neurons^41^. Naltrexone is a competitive antagonist at mu opioid receptor and antagonizes antinociception of 30 mg/kg morphine with an IC50 of 0.007 mg/kg in mice^42^. Our data confirms that sEV analgesia can be reversed by the non-selective opioid antagonist naltrexone, suggesting leu-enkephalin in sEVs contributes to a short-term analgesia. Endogenous opioid peptides can also be released by immune cells^43^. For example, specific T cell subsets release endogenous opioid peptides that can suppress or resolve pain^44^. Macrophages also generate and release opioid peptides under inflammatory and neuropathic pain states^45^, and spinal microglial β-endorphin expression produces antinociception^46^. Thus, we cannot rule out that sEVs released by immune cells in recipient mice indirectly increase their basal pain threshold. The rapid loss of enkephalin analgesia has been attributed to its rapid hydrolysis in the body^47^. However, sEVs can extend the half-life of leu-enkephalin by protecting it from being degraded, and this could be another mode by which leu-enkephalin exerts its effect *in vivo*.

SNI model mice that prophylactically received 10 µg serum-derived sEVs from naïve or SNI model donors showed increased mechanical threshold 1- and 3-days post-surgery. Pretreatment with sEVs did not prevent mechanical hypersensitivity after SNI surgery, but they delayed SNI-induced pain hypersensitivity in the initial stages. Prophylactic serum-derived sEVs resolved mechanical allodynia faster than vehicle after CFA injection, but the dose of sEVs required was ten times higher than the effective dose of RAW 264.7 macrophage derived sEVs^13^. Thus, heterogenous sEVs in serum have a much lower potency than sEVs secreted by macrophage cells. Serum-derived sEVs are produced by many different types of donor cells and can harbor cargo molecules with varying efficacy against pain sensitivity. Our studies using macrophage-derived sEVs^13^, suggest that monocyte/macrophage-derived sEVs in serum could be one of the contributing factors that resolve hypersensitivity in CFA model mice.

The DWB system measures weight distribution on all four paws and provides a way to evaluate non-reflexive pain in mice^48^. Compared to evoked measures, DWB is semi-automated, objective, aligned with clinically observed behavioral outcomes of pain, and therefore a more reliable method for assessing pain^49^. Our DWB studies showed that mice administered sEVs from naïve donors differed from those mice that received sEVs from SNI donor mice or PBS control at 3h post-injection. Administering naïve sEVs significantly increased weight distribution of hind paw compared to the SNI sEV treatment or control group. The differences between the two sEV treated groups at 3 hours could be due to enriched proinflammatory proteins such as C5a^14^ in sEVs from SNI model mice. Interestingly, basal mechanical threshold assessed by von Frey test was also elevated at this time point but all sEV treated mice responded similarly. In mice that received prophylactic sEV treatment, CFA injection two weeks later induced a much smaller change in weight distribution of hind paws compared to the control group at 3 days post CFA indicating alleviation of pain by sEVs.

Neuroimmune interactions contribute to the development or maintenance of chronic pain^50, 51, 52^. Macrophages and T cells enhance pain via proinflammatory mediators and resolve pain via anti-inflammatory mediators such as IL-10 and endogenous opioids^53^. Immune cells can also stimulate the release of endogenous opioid peptides or enhance their synthesis^43^, which can then bind opioid receptors on these cells and modulate their function^54^. Further, macrophage-nociceptor interactions are mediated by miRNAs and miRNA-containing exosomes in chronic pain^55^. Exosomes released by cultured sensory neurons upon capsaicin activation are readily phagocytosed by macrophages, which increases their expression of miR-21-5p and promotes a pro-inflammatory phenotype^56^. A single intrathecal injection of human conditioned serum produced long-lasting inhibition of paclitaxel chemotherapy-induced mechanical allodynia in mice, which was significantly diminished upon removal of exosomes^57^. A recent study reported that plasma EVs (pEVs) of healthy human donors are endogenous homeostatic modulators of macrophages that control macrophage activation, decrease inflammatory cytokines, and promote tissue repair. pEVs induce prostaglandin PGE2 expression by macrophages, which in turn critically mediates pEVs anti-inflammatory activity^58^. The sEVs from naïve mice could have comparable functions and need further investigations.

High-plex ChipCytometry enabled us to identify changes in immune cell populations in spinal cord and DRG tissues simultaneously in sEV treated mice. We found that temporal and spatial alterations in immune cell number and subpopulations could attenuate inflammatory pain. Specifically prophylactic sEV treatment reduced the number of NK and NKT cells in spinal cord and increased CD206^+^ anti-inflammatory macrophages in DRG after CFA injection. A recent study showed upregulation of IL-10 in spinal macrophages resolved neuroinflammation and neuropathic pain^59^. IL-10-producing monocytes/macrophages in DRG promote resolution of transient inflammatory hyperalgesia^60^. Though nerve injury can increase the number and activation of sensory neuron-associated macrophages in the sciatic nerve and sensory ganglia, it is unclear if this occurs via infiltration of blood monocytes or local proliferation^61^. One study reported that increase in the number of macrophages in the sensory ganglia after SNI is a consequence of the proliferation of resident CX3CR1^+^ macrophages and not due to infiltration of peripheral blood monocytes^62^. In yet another study, a limited number of neutrophils and macrophages/monocytes infiltrate the spinal cord after peripheral injury but their impact on pain persistence or resolution remains unknown^63^. Neutrophils and macrophages are innate immune cells that can induce a transition from acute to chronic pain and maintain pain. Adaptive immune cells like B cells and T cells mainly initiate chronic pain, though T cells can also act to resolve pain^64^. This creates an interesting dichotomy in T cell regulation of pain. Local depletion of macrophages in CFA model mice attenuates inflammatory pain, but after persistent inflammatory pain is established, local macrophage depletion no longer impacts pain^65^. Targeted depletion of T cells or neutrophils also did not affect pain behaviors in CFA model mice^66^. Further, neutrophils secrete proinflammatory mediators but also produce opioid peptides that might reduce pain. Neutrophil mediated inflammatory responses in individuals experiencing pain relief indicate that active inflammatory responses underlie pain resolution^67^. NK cells are traditionally viewed as pro-inflammatory, but they are also functionally important pain resolvers. B lymphocytes produce antibodies involved in humoral immunity and also present antigens^64^. Thus, temporal patterns of immune cell recruitment and activation confer distinct roles in regulating different types of pain. The accelerated resolution of inflammatory pain hypersensitivity mediated by sEVs could be coordinated by multiple types of immune cells.

sEVs from SNI model male donor mice showed enrichment of immune markers such as CD4, CD24 and activation markers like CD40 and CD44 compared to sEVs from naïve male mice. Based on the immune markers expressed on sEVs in serum, these different populations of sEVs are likely released by specific immune cells. sEV cargo reflect a subset of the biological functions of the parent cell releasing these vesicles. Several studies show that immune cell sEVs can be therapeutic, including sEVs released by APCs that present peptide loaded MHC and costimulatory molecules such as CD80 and CD86 to T cells^68^. The majority of sEV immune marker expression was similar between naive male and female mice, with a few notable exceptions including CD40, whose ligand CD40L is located on the X chromosome. Additional studies will determine if this explains lower prophylactic efficacy of female derived sEVs in CFA model female mice. sEVs isolated two weeks after SNI surgery showed no difference in size by both NTA and microflow cytometry, consistent with SNI model mice four weeks post injury^14^. Size measurements by ExoView however, showed an increase in size of CD81, CD45, and F4/80 captured populations. This suggests cell dependent effects on sEV size in context of SNI, which may be similar to what has been observed with neutrophil derived sEVs in context of PMA^69^. To determine how SNI surgery impacts particle size, additional future studies are needed to determine the major source of sEVs in circulation and how nerve injury influences genes involved in sEV biogenesis. Insights could be gained by a systematic analysis of sEVs isolated from various primary cells and elucidating the molecular and behavioral consequences upon uptake.

## Acknowledgements

We thank Dr. Shaye Hagler, Dr. Xenia Meshik, Dr. Adam Northcutt, Spencer Schwarz and Tim Sindelar from Canopy Biosciences for their technical assistance with ChipCytometry, Dr. Luca Musante from the University of Pennsylvania Extracellular Vesicle Core Facility for his help with ExoView studies, Dr. Pat Loll from Drexel University College of Medicine for his expertise in sample preparation for determining Leu-enkephalin, Dr. Bradley Nash for editing the manuscript. This study was funded by NIH NINDS R01 NS102836 and partially supported by NINDS R01NS129191 and NINDS 1RF1NS130481 to Seena Ajit. We would like to acknowledge the support provided by the Proteomics and Metabolomics Shared Resource of the Wake Forest School of Medicine and Wake Forest Baptist Comprehensive Cancer Center (NIH/NCI P30 CA12197). Xuan Luo, Richa Pande, and Jason Wickman received Dean’s Fellowship for Excellence in Collaborative or Themed Research from Drexel University College of Medicine.

## Author contributions

Conceived the study S.K.A; designed and performed experiments: Z.L., X.L., J.R.W., D.R., R.P., Y.T., V.T. J.L.; analyzed data: Z.L., X.L., J.R.W., D.R., R.P., J.L., D.P., A.S., S.K.A.; supervision: C.M.F., D.P., A.S., S.K.A.; writing: Z.L., X.L., S.K.A.; editing: J.R.W., D.R., R.P., J.L. C.M.F. D.P., A.S.; funding acquisition: S.K.A.

## Supplementary Figures

**Supp Fig. 1 Uptake of intrathecally injected sEVs in spinal cord and DRG.** sEVs were labelled with membrane labelling PKH26 dye (red). Ten µg labelled sEVs were intrathecally injected into each naïve mouse. After 6h, mice were scarified and the spinal cord and L4-L5 DRG tissues were collected. Uptake of sEVs were observed by Olympus FV3000 microscope (60 X magnification). Neurons were stained with MAP2A (green), astrocytes were stained with GFAP (green), and microglia/macrophage were stained with IBA1 (green), while cell nuclei were stained with DAPI (blue). The sEVs could be visualized in different cells (solid white arrows). PKH26 dye alone served as a negative control for sEV staining (not shown). Scale bar = 20 µm.

**Supp Fig. 2 Low dose (1 µg) sEVs did not change the basal mechanical and thermal threshold in recipient mice. a** Schematic of *in vivo* experiment. sEVs purified from serum of male C57BL/6 mice 2 weeks post SNI surgery or naïve mice were administrated intrathecally into another group of naïve male recipient mice followed by behavior tests. **b** Mechanical and **c** thermal threshold was not significantly different before and after the single intrathecal injection of sEVs (n=5). Data shown are mean ± SEM. Statistical analysis was determined by two-way repeated-measures ANOVA.

**Supp Fig. 3 sEVs from female mice did not alter basal mechanical or thermal threshold in female recipient mice. a** Schematic of *in vivo* experiment. sEVs purified from serum of female C57BL/6 mice 2 weeks post SNI surgery or naïve mice were administrated intrathecally into another group of naïve female recipient mice followed by behavior tests. **b** There was significant difference in mechanical and **c** thermal threshold in recipient mice before and after the single intrathecal injection of sEVs (n=5).

**Supp Fig. 4 Detection of endogenous opioid peptides as cargo in serum sEVs.** Liquid chromatography with tandem mass spectrometry of serum sEVs samples from male mice showing representative MS/MS spectrum of nociceptin in serum-derived sEVs.

**Supp Fig. 5 Confirmation of naltrexone efficacy at 3h and 24h post-treatment using morphine as positive control for the two time points that were significant in sEV experiments. a** Schematic of *in vivo* experiment to test if naltrexone can reverse morphine-induced analgesia. Naltrexone (10mg/kg) was administrated intraperitoneally into naïve 9-week-old male C57BL/6 mice, and after 2h 30min or 24h post injection, 0.5 µg morphine was injected intrathecally. Behavior studies were performed at 30 min after morphine injection. **b** Pretreatment with naltrexone for 3 hours reversed analgesic effect of morphine. Naltrexone alone did not affect the mechanical threshold and mice that received only morphine showed an increase in basal mechanical threshold at 3h post PBS injection. Mice treated with both naltrexone and morphine exhibited lower PWT than morphine group (n=5), showing naltrexone could reverse morphine-induced short-term analgesic effect 3h post naltrexone injection. **c** Naltrexone reversed analgesic effect of morphine at 24h. Pretreatment with naltrexone for 24 hours reversed analgesic effect of morphine even at 24h post injection (n=5). Data shown are mean ± SEM. Statistical analysis was determined by two-way repeated-measures ANOVA followed by Bonferroni test * p <0.05, ** p <0.01, *** p <0.001, **** p <0.0001, ## p <0.01, ### p <0.001.

**Supp Fig. 6 Prophylactic intrathecal injections of low dose (1 µg) sEVs did not alter CFA induced mechanical hypersensitivity. a** Schematic representation of experimental design. One µg sEVs from male C57BL/6 naïve and SNI model donor mice two weeks after injury were injected intrathecally into nine weeks old male C57BL/6 recipient mice. CFA model was established in these recipient mice 2 weeks post sEVs injection. **b** Mechanical allodynia and **c** thermal hyperalgesia in CFA model mice did not show significant differences in sEV treated mice compared to PBS control group (n=5). Data shown are mean ± SEM, two-way repeated-measures ANOVA (n=5).

**Supp Fig. 7 Prophylactic intrathecal injections of 10 µg sEVs from female mice did not contribute to resolution of CFA induced mechanical hypersensitivity in female recipient mice. a** Schematic representation of experimental design. The sEVs from female C57BL/6 naïve and SNI model donor mice two weeks after injury were injected intrathecally into nine weeks old female C57BL/6 recipient mice. CFA model was established in these recipient mice 2 weeks post sEVs injection. **b** Mechanical allodynia and **c** thermal hyperalgesia in CFA model mice did not show significant differences in sEV treated mice compared to PBS control group (n=5). Data shown are mean ± SEM, two-way repeated-measures ANOVA (n=5).

**Supp Fig 8** Immune cells separated as ipsi and contralateral to CFA injection and dorsal and ventral spinal cord sections as determined using ChipCytometry.

**Supp Fig 9a** sEV marker distribution by capture target and **b** detector antibody fluorescence intensity.

**Supp Fig 10a** Representative gating scheme of CFSE positive sEVs for sEVs labeled by CFDA-SE with or without Triton. Concatenated plots of sEV samples gated by MALsLo or MALSHi. **b** Quantification of Fig. 10a showing the percentage of MALSLo or MALSHi events in non-treated or treated sEVS, with a representative histogram of MALS intensity for Exo sEVs and quantification of the MALS gMFI for each sample type below it (n=4 per group). Data shown are mean ± SD. Statistical analysis was determined by one-way or two-way repeated-measures ANOVA followed by Bonferroni test, * p < 0.05 ** p < 0.01 *** p < 0.001 **** p < 0.0001.

## Supplementary Tables

**Supp Table 1** Differentially expressed miRNAs in sEVs from naïve, sham control and SNI model mice two weeks post surgery.

**Supp Table 2** miRNAs highly expressed in all serum derived sEVs.

**Supp Table 3** List of significantly enriched pathways.

**Supp Table 4** Experimental design showing treatment groups in recipient mice. L4-L5 spinal cord and DRG from recipient male mice were obtained at time points that were most efficacious in behavior testing (n=2). sEVs from RAW 264.7 were included as a positive control for homogenous sEV source of macrophage under culture conditions.

**Supp Table 5** The 16-plex panel of markers used to identify the populations in ChipCytometry.

## References

1. Cata JP, Uhelski ML, Gorur A, Dougherty PM. Nociception and Pain: New Roles for Exosomes. The Neuroscientist 28, 349–363 (2022).

2. van Niel G, D’Angelo G, Raposo G. Shedding light on the cell biology of extracellular vesicles. Nature Reviews Molecular Cell Biology 19, 213–228 (2018).

3. Dragovic RA, et al. Sizing and phenotyping of cellular vesicles using Nanoparticle Tracking Analysis. Nanomedicine 7, 780–788 (2011).

4. Théry C, et al. Minimal information for studies of extracellular vesicles 2018 (MISEV2018): a position statement of the International Society for Extracellular Vesicles and update of the MISEV2014 guidelines. Journal of extracellular vesicles 7, 1535750 (2018).

5. Mathieu M, Martin-Jaular L, Lavieu G, Théry C. Specificities of secretion and uptake of exosomes and other extracellular vesicles for cell-to-cell communication. Nat Cell Biol 21, 9–17 (2019).

6. Cheng L, Hill AF. Therapeutically harnessing extracellular vesicles. Nature Reviews Drug Discovery 21, 379–399 (2022).

7. Teng F, Fussenegger M. Shedding Light on Extracellular Vesicle Biogenesis and Bioengineering. Adv Sci (Weinh) 8, 2003505 (2020).

8. Ren J, Liu N, Sun N, Zhang K, Yu L. Mesenchymal Stem Cells and their Exosomes: Promising Therapeutics for Chronic Pain. Current stem cell research & therapy 14, 644–653 (2019).

9. D’Agnelli S, Gerra MC, Bignami E, Arendt-Nielsen L. Exosomes as a new pain biomarker opportunity. Molecular pain 16, 1744806920957800–1744806920957800 (2020).

10. Shiue SJ, et al. Mesenchymal stem cell exosomes as a cell-free therapy for nerve injury-induced pain in rats. Pain 160, 210–223 (2019).

11. He L, et al. Bone marrow mesenchymal stem cell-derived exosomes protect cartilage damage and relieve knee osteoarthritis pain in a rat model of osteoarthritis. Stem Cell Res Ther 11, 276 (2020).

12. McDonald MK, et al. Functional significance of macrophage-derived exosomes in inflammation and pain. PAIN® 155, 1527–1539 (2014).

13. Jean-Toussaint R, et al. Therapeutic and prophylactic effects of macrophage-derived small extracellular vesicles in the attenuation of inflammatory pain. Brain Behav Immun 94, 210–224 (2021).

14. Jean-Toussaint R, Tian Y, Chaudhuri AD, Haughey NJ, Sacan A, Ajit SK. Proteome characterization of small extracellular vesicles from spared nerve injury model of neuropathic pain. J Proteomics 211, 103540 (2020).

15. Sosanya NM, et al. Identifying Plasma Derived Extracellular Vesicle (EV) Contained Biomarkers in the Development of Chronic Neuropathic Pain. The Journal of Pain 21, 82–96 (2020).

16. Jang JH, Clark JD, Li X, Yorek MS, Usachev YM, Brennan TJ. Nociceptive sensitization by complement C5a and C3a in mouse. Pain 148, 343–352 (2010).

17. Decosterd I, Woolf CJ. Spared nerve injury: an animal model of persistent peripheral neuropathic pain. Pain 87, 149–158 (2000).

18. Quadros AU, Pinto LG, Fonseca MM, Kusuda R, Cunha FQ, Cunha TM. Dynamic weight bearing is an efficient and predictable method for evaluation of arthritic nociception and its pathophysiological mechanisms in mice. Scientific reports 5, 14648 (2015).

19. Pitzer C, Kuner R, Tappe-Theodor A. EXPRESS: Voluntary and evoked behavioral correlates in neuropathic pain states under different housing conditions. Mol Pain 12, (2016).

20. Gupta R, Luo X, Lin Z, Tian Y, Ajit SK. Uptake of Fluorescent Labeled Small Extracellular Vesicles In Vitro and in Spinal Cord. Journal of visualized experiments: JoVE, (2021).

21. Agarwal V, Bell GW, Nam JW, Bartel DP. Predicting effective microRNA target sites in mammalian mRNAs. eLife 4, (2015).

22. Daaboul GG, et al. Digital Detection of Exosomes by Interferometric Imaging. Scientific reports 6, 37246 (2016).

23. Kowal J, Tkach M, Thery C. Biogenesis and secretion of exosomes. Curr Opin Cell Biol 29, 116–125 (2014).

24. Luo X, Jean-Toussaint R, Tian Y, Balashov SV, Sacan A, Ajit SK. Small Extracellular Vesicles From Spared Nerve Injury Model and Sham Control Mice Differentially Regulate Gene Expression in Primary Microglia. The journal of pain: official journal of the American Pain Society, (2023).

25. Bryk M, Karnas E, Mlost J, Zuba-Surma E, Starowicz K. Mesenchymal stem cells and extracellular vesicles for the treatment of pain: Current status and perspectives. Br J Pharmacol, (2021).

26. Ting E, et al. Role of complement C5a in mechanical inflammatory hypernociception: potential use of C5a receptor antagonists to control inflammatory pain. Br J Pharmacol 153, 1043–1053 (2008).

27. Bui TM, Wiesolek HL, Sumagin R. ICAM-1: A master regulator of cellular responses in inflammation, injury resolution, and tumorigenesis. J Leukoc Biol 108, 787–799 (2020).

28. Sztáray J, Memboeuf A, Drahos L, Vékey K. Leucine enkephalin--a mass spectrometry standard. Mass Spectrom Rev 30, 298–320 (2011).

29. Pitzer C, Kuner R, Tappe-Theodor A. Voluntary and evoked behavioral correlates in inflammatory pain conditions under different social housing conditions. Pain Rep 1, e564 (2016).

30. Griffioen MA, Dernetz VH, Yang GS, Griffith KA, Dorsey SG, Renn CL. Evaluation of dynamic weight bearing for measuring nonevoked inflammatory hyperalgesia in mice. Nurs Res 64, 81–87 (2015).

31. Jarosch S, Köhlen J, Wagner S, D’Ippolito E, Busch DH. ChipCytometry for multiplexed detection of protein and mRNA markers on human FFPE tissue samples. STAR Protoc 3, 101374 (2022).

32. Thery C, Ostrowski M, Segura E. Membrane vesicles as conveyors of immune responses. Nat Rev Immunol 9, 581–593 (2009).

33. Gardiner C, et al. Measurement of refractive index by nanoparticle tracking analysis reveals heterogeneity in extracellular vesicles. Journal of extracellular vesicles 3, 25361 (2014).

34. Rupert DLM, et al. Effective Refractive Index and Lipid Content of Extracellular Vesicles Revealed Using Optical Waveguide Scattering and Fluorescence Microscopy. Langmuir: the ACS journal of surfaces and colloids 34, 8522–8531 (2018).

35. de Rond L, et al. Refractive index to evaluate staining specificity of extracellular vesicles by flow cytometry. Journal of extracellular vesicles 8, 1643671 (2019).

36. Welsh JA, et al. Towards defining reference materials for measuring extracellular vesicle refractive index, epitope abundance, size and concentration. Journal of extracellular vesicles 9, 1816641 (2020).

37. Pleet ML, et al. Extracellular Vesicle Refractive Index Derivation Utilizing Orthogonal Characterization. Nano letters 23, 9195–9202 (2023).

38. Honda M, et al. Spinorphin, an endogenous inhibitor of enkephalin-degrading enzymes, potentiates leu-enkephalin-induced anti-allodynic and antinociceptive effects in mice. Jpn J Pharmacol 87, 261–267 (2001).

39. Porreca F, Mosberg HI, Hurst R, Hruby VJ, Burks TF. Roles of mu, delta and kappa opioid receptors in spinal and supraspinal mediation of gastrointestinal transit effects and hot-plate analgesia in the mouse. J Pharmacol Exp Ther 230, 341–348 (1984).

40. Janecka A, Fichna J, Janecki T. Opioid receptors and their ligands. Curr Top Med Chem 4, 1–17 (2004).

41. Zhang XY, et al. Different neuronal populations mediate inflammatory pain analgesia by exogenous and endogenous opioids. Elife 9, (2020).

42. Porter SJ, Somogyi AA, White JM. In vivo and in vitro potency studies of 6beta-naltrexol, the major human metabolite of naltrexone. Addict Biol 7, 219–225 (2002).

43. Plein LM, Rittner HL. Opioids and the immune system - friend or foe. Br J Pharmacol 175, 2717–2725 (2018).

44. Laumet G, Ma J, Robison AJ, Kumari S, Heijnen CJ, Kavelaars A. T Cells as an Emerging Target for Chronic Pain Therapy. Front Mol Neurosci 12, 216 (2019).

45. Sauer RS, et al. Toll like receptor (TLR)-4 as a regulator of peripheral endogenous opioid-mediated analgesia in inflammation. Mol Pain 10, 10 (2014).

46. Wu HY, et al. Spinal interleukin-10 produces antinociception in neuropathy through microglial beta-endorphin expression, separated from antineuroinflammation. Brain Behav Immun 73, 504–519 (2018).

47. Hambrook JM, Morgan BA, Rance MJ, Smith CF. Mode of deactivation of the enkephalins by rat and human plasma and rat brain homogenates. Nature 262, 782–783 (1976).

48. Laux-Biehlmann A, Boyken J, Dahllof H, Schmidt N, Zollner TM, Nagel J. Dynamic weight bearing as a non-reflexive method for the measurement of abdominal pain in mice. Eur J Pain 20, 742–752 (2016).

49. Klokker L, et al. Dynamic weight-bearing assessment of pain in knee osteoarthritis: construct validity, responsiveness, and interpretability in a research setting. Health Qual Life Outcomes 14, 91 (2016).

50. Pinho-Ribeiro FA, Verri WA, Jr., Chiu IM. Nociceptor Sensory Neuron-Immune Interactions in Pain and Inflammation. Trends Immunol 38, 5–19 (2017).

51. Fiore NT, Debs SR, Hayes JP, Duffy SS, Moalem-Taylor G. Pain-resolving immune mechanisms in neuropathic pain. Nat Rev Neurol 19, 199–220 (2023).

52. Mustafa S, et al. One immune system plays many parts: The dynamic role of the immune system in chronic pain and opioid pharmacology. Neuropharmacology 228, 109459 (2023).

53. Kavelaars A, Heijnen CJ. Immune regulation of pain: Friend and foe. Sci Transl Med 13, eabj7152 (2021).

54. Machelska H, Celik MO. Opioid Receptors in Immune and Glial Cells-Implications for Pain Control. Front Immunol 11, 300 (2020).

55. Chen O, Donnelly CR, Ji RR. Regulation of pain by neuro-immune interactions between macrophages and nociceptor sensory neurons. Curr Opin Neurobiol 62, 17–25 (2020).

56. Simeoli R, et al. Exosomal cargo including microRNA regulates sensory neuron to macrophage communication after nerve trauma. Nature communications 8, 1778 (2017).

57. Buchheit T, et al. Intrathecal administration of conditioned serum from different species resolves Chemotherapy-Induced neuropathic pain in mice via secretory exosomes. Brain Behav Immun 111, 298–311 (2023).

58. Adamczyk AM, et al. Extracellular vesicles from human plasma dampen inflammation and promote tissue repair functions in macrophages. Journal of extracellular vesicles 12, 12331 (2023).

59. Niehaus JK, Taylor-Blake B, Loo L, Simon JM, Zylka MJ. Spinal macrophages resolve nociceptive hypersensitivity after peripheral injury. Neuron, (2021).

60. Willemen HL, et al. Monocytes/Macrophages control resolution of transient inflammatory pain. The journal of pain: official journal of the American Pain Society 15, 496–506 (2014).

61. Silva CEA, Guimarães RM, Cunha TM. Sensory neuron-associated macrophages as novel modulators of neuropathic pain. Pain Rep 6, e873 (2021).

62. Guimarães RM, et al. Neuron-associated macrophage proliferation in the sensory ganglia is associated with peripheral nerve injury-induced neuropathic pain involving CX3CR1 signaling. eLife 12, e78515 (2023).

63. Huck NA, et al. Sex-distinct microglial activation and myeloid cell infiltration in the spinal cord after painful peripheral injury. Neurobiol Pain 12, 100106 (2022).

64. Yang J-X, et al. Potential Neuroimmune Interaction in Chronic Pain: A Review on Immune Cells in Peripheral and Central Sensitization. Frontiers in Pain Research 3, (2022).

65. Raoof R, Willemen HLDM, Eijkelkamp N. Divergent roles of immune cells and their mediators in pain. Rheumatology 57, 429–440 (2017).

66. Ghasemlou N, Chiu IM, Julien J-P, Woolf CJ. CD11b+Ly6G− myeloid cells mediate mechanical inflammatory pain hypersensitivity. Proceedings of the National Academy of Sciences 112, E6808–E6817 (2015).

67. Parisien M, et al. Acute inflammatory response via neutrophil activation protects against the development of chronic pain. Science Translational Medicine 14, eabj9954 (2022).

68. Robbins PD, Dorronsoro A, Booker CN. Regulation of chronic inflammatory and immune processes by extracellular vesicles. Journal of Clinical Investigation 126, 1173+ (2016).

69. Alvarez-Jiménez VD, et al. Extracellular Vesicles Released from Mycobacterium tuberculosis-Infected Neutrophils Promote Macrophage Autophagy and Decrease Intracellular Mycobacterial Survival. Front Immunol 9, 272 (2018).

