## Supplemental Figures for "Inflammatory pain resolution by mouse serum-derived small extracellular vesicles"

Supplementary Fig. 1

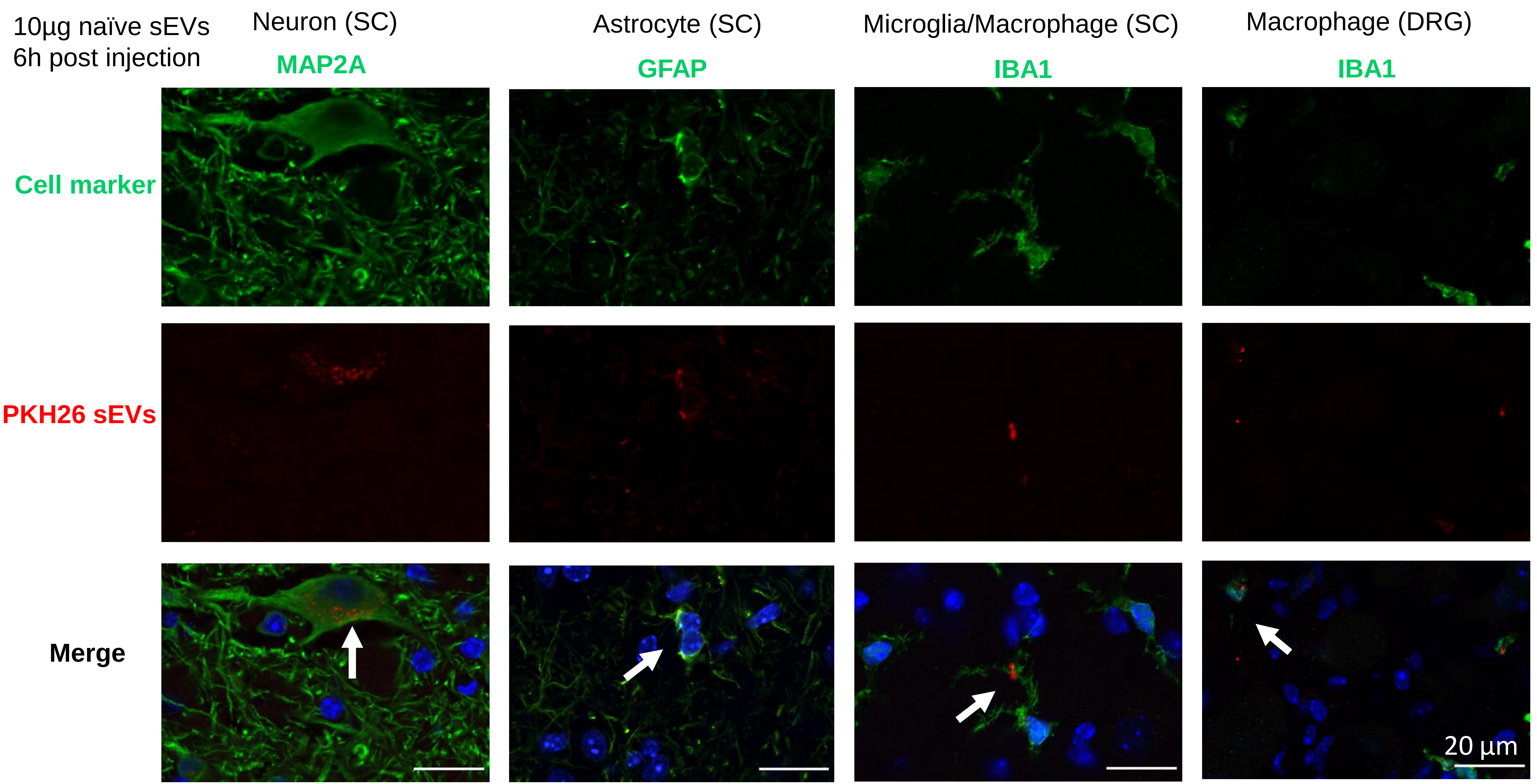

Supplementary Fig. 2

a

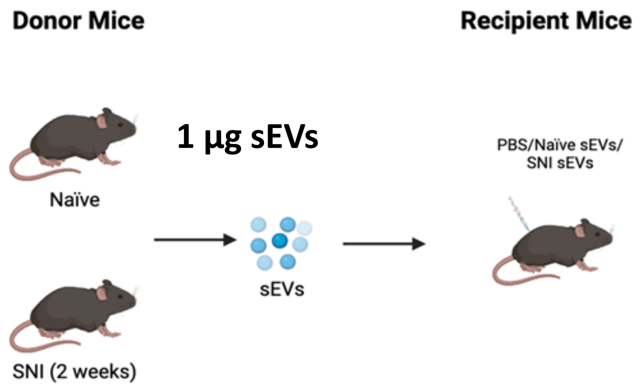


b

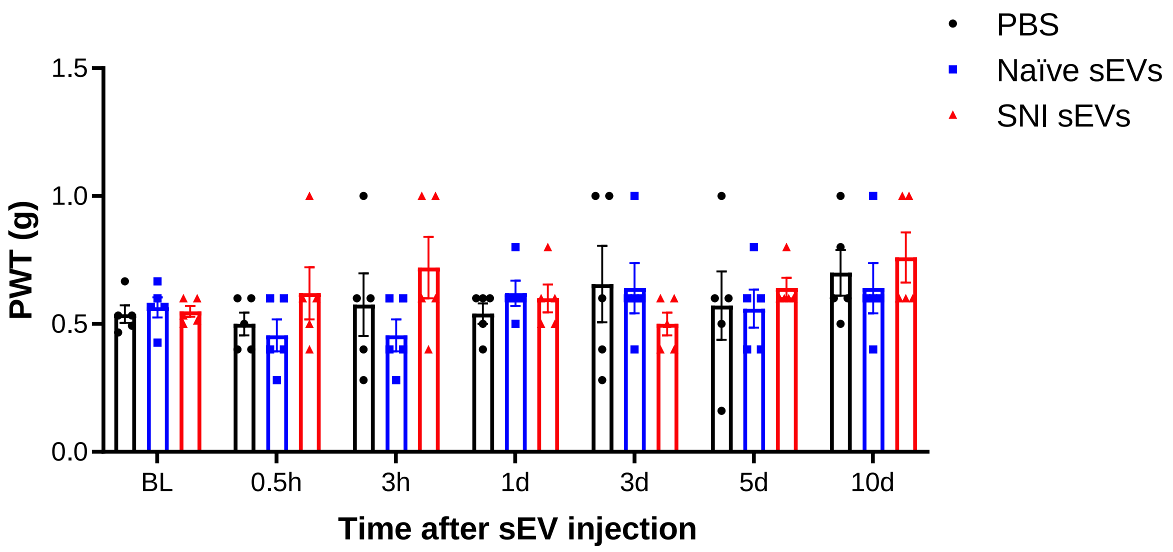


c

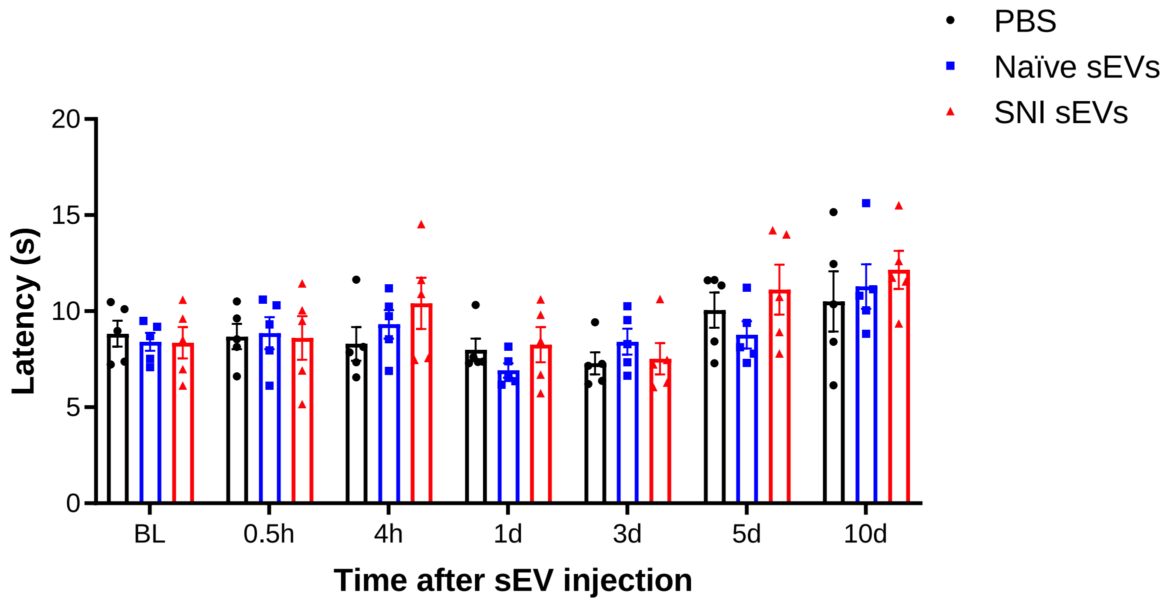

### Supplementary Fig. 3

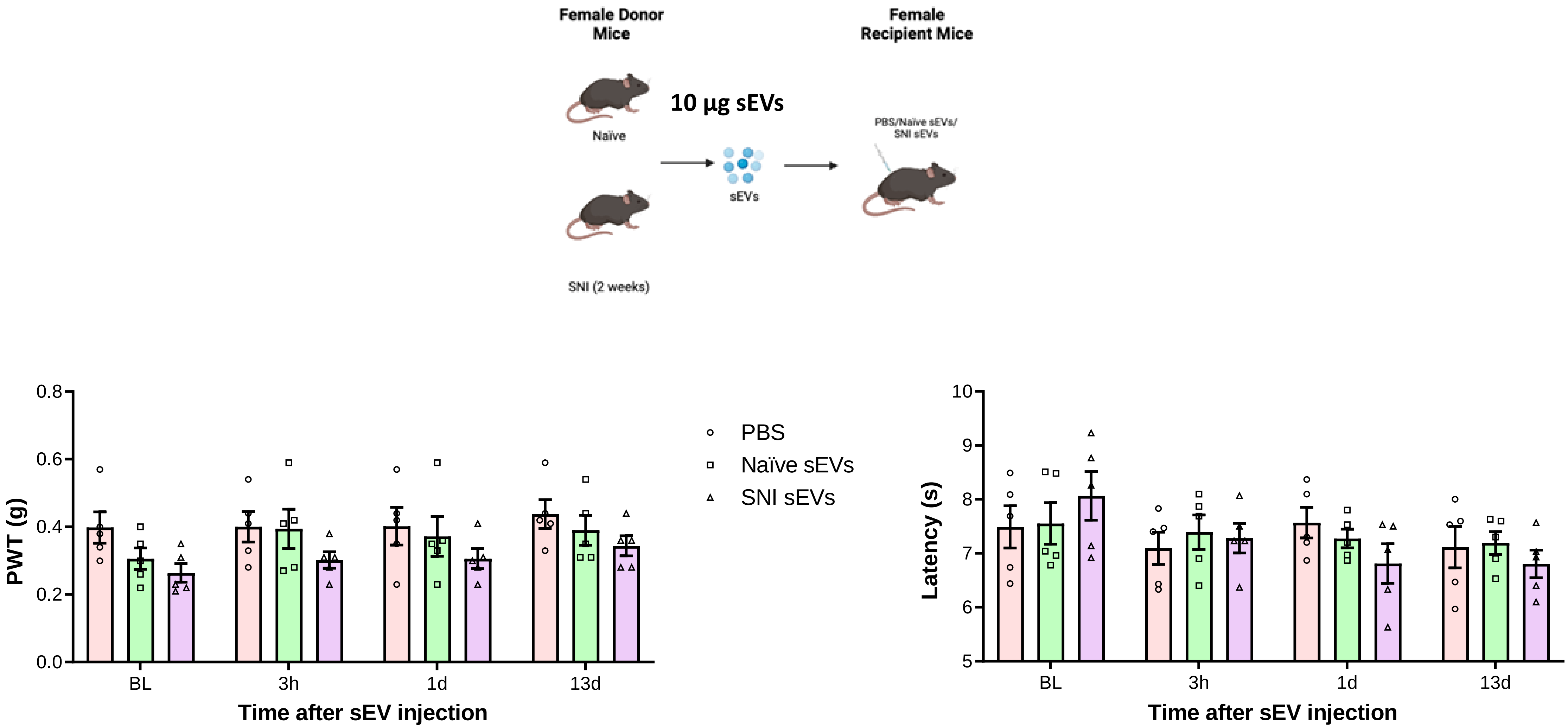

Supplementary Fig. 4

a

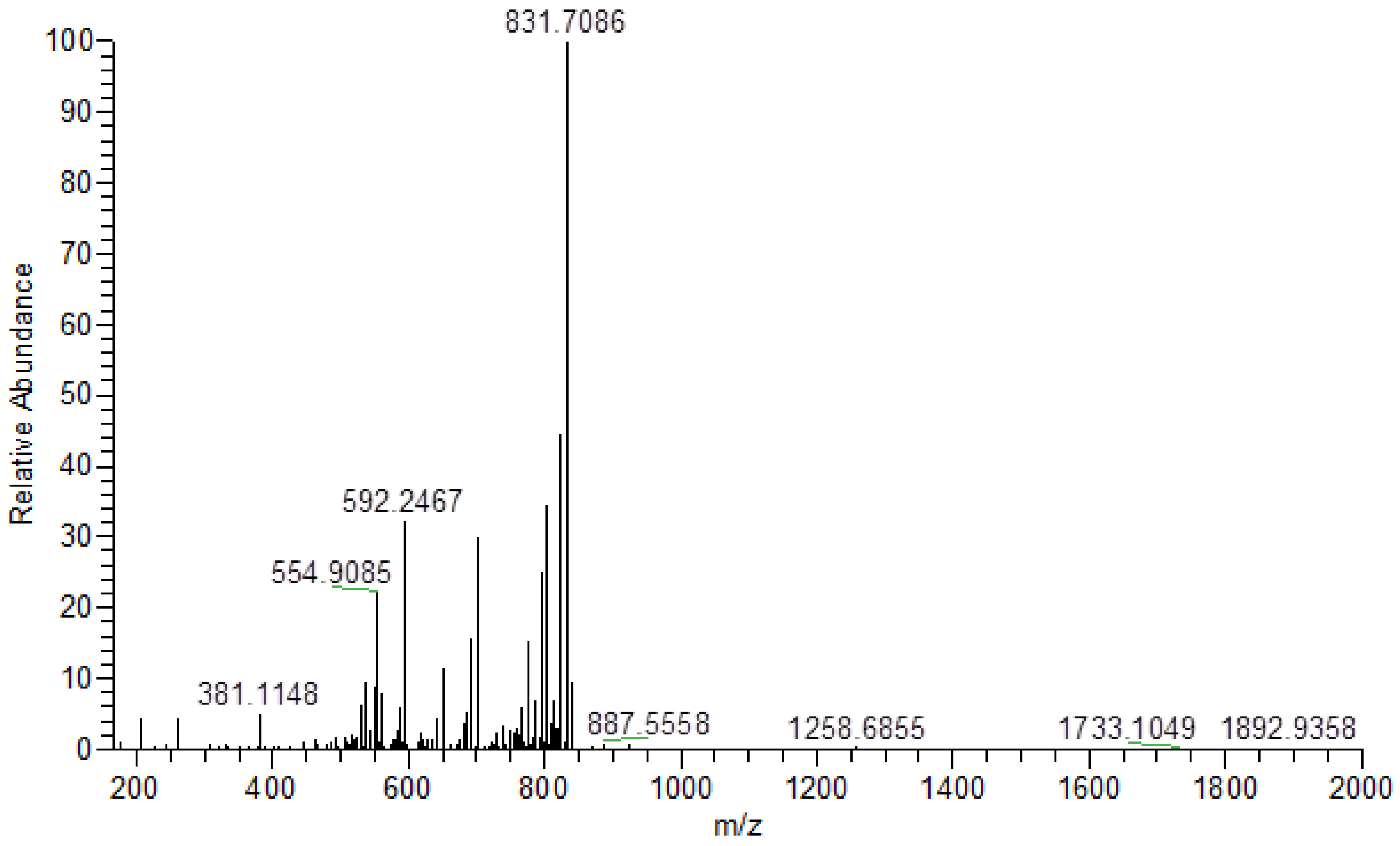

Supplementary Fig. 5a

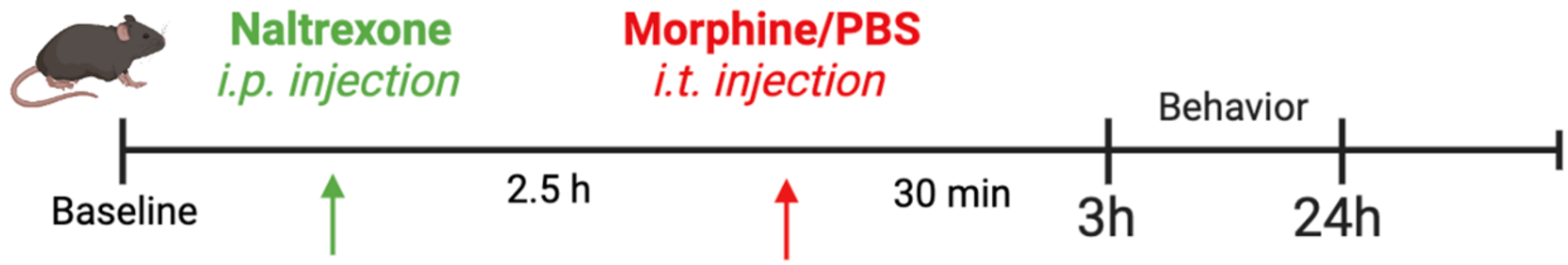


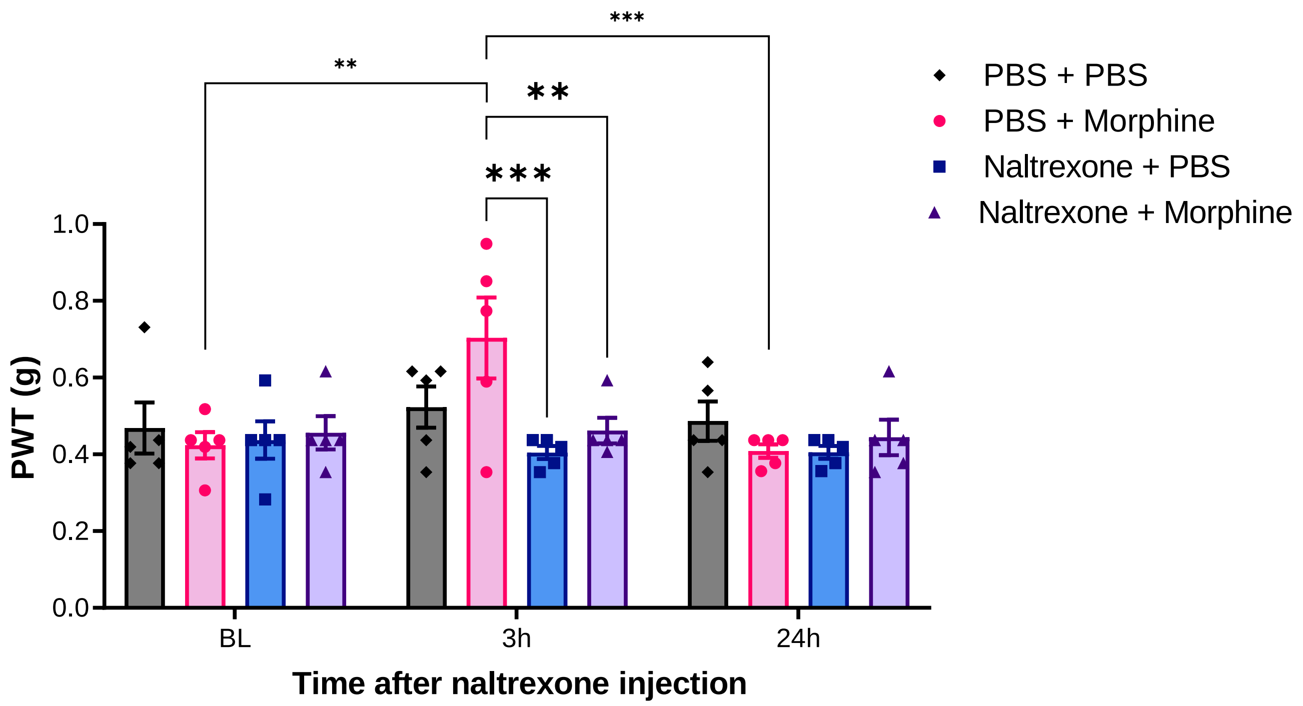

Supplementary Fig. 5b

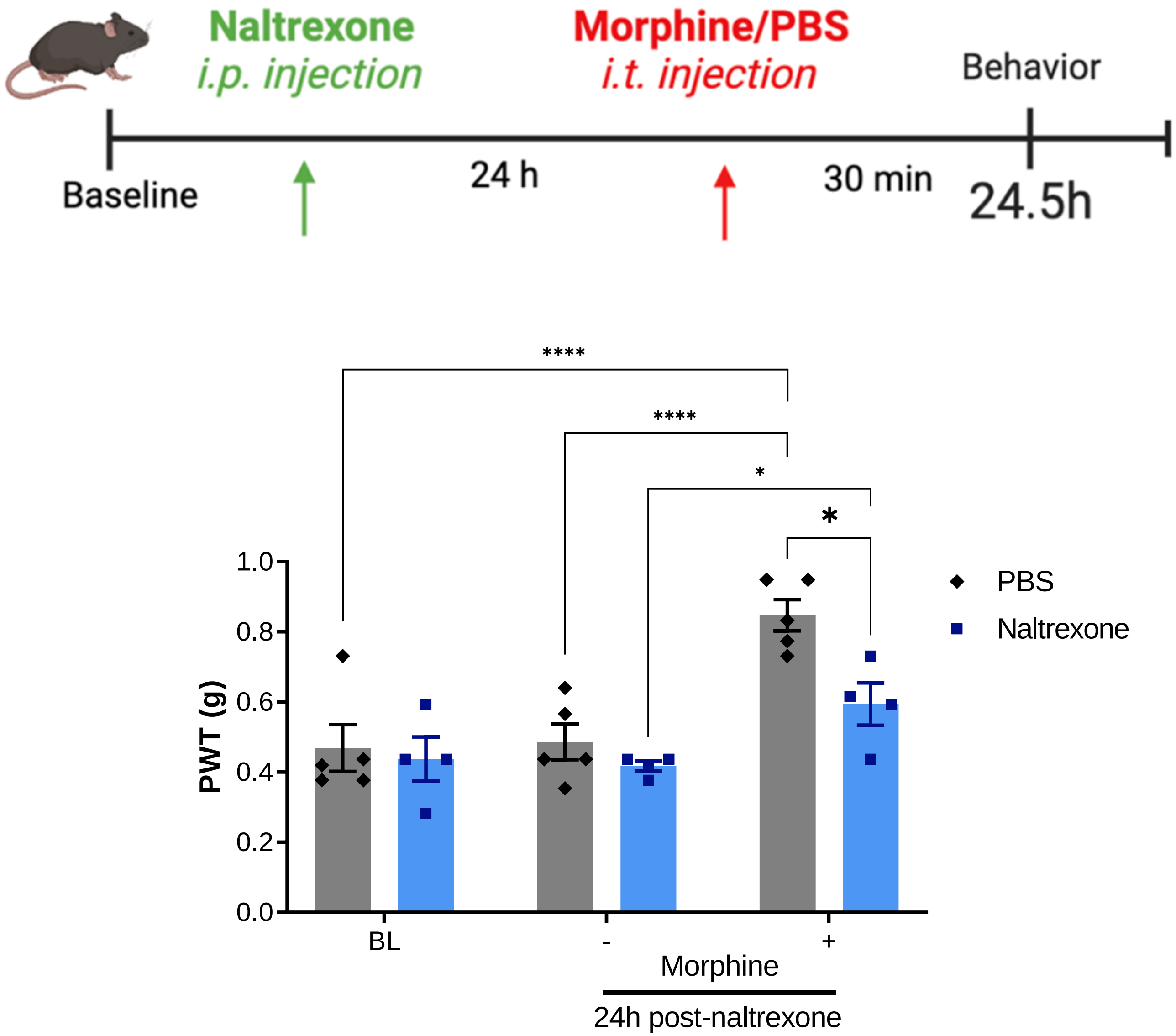

Supplementary Fig. 6

a

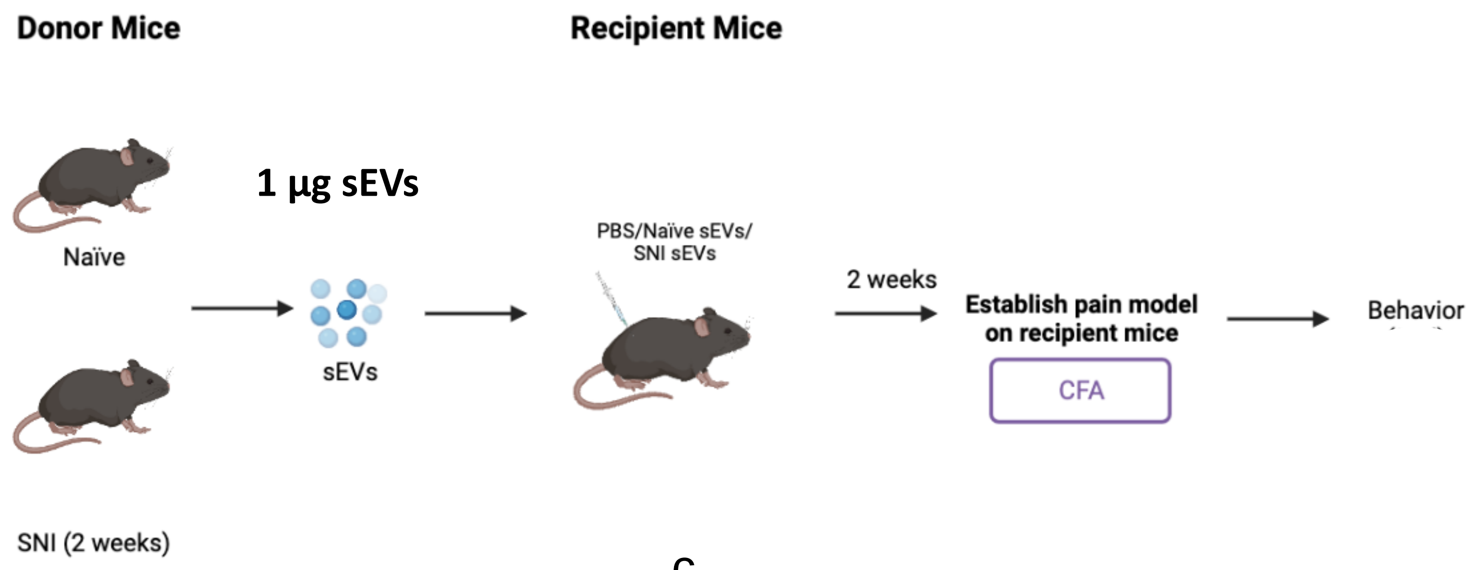


b

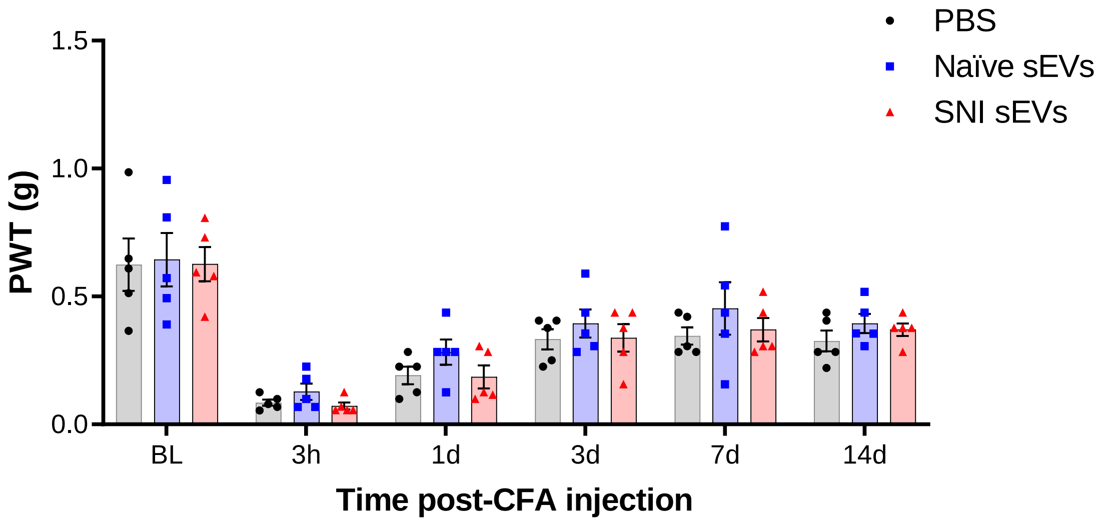


c

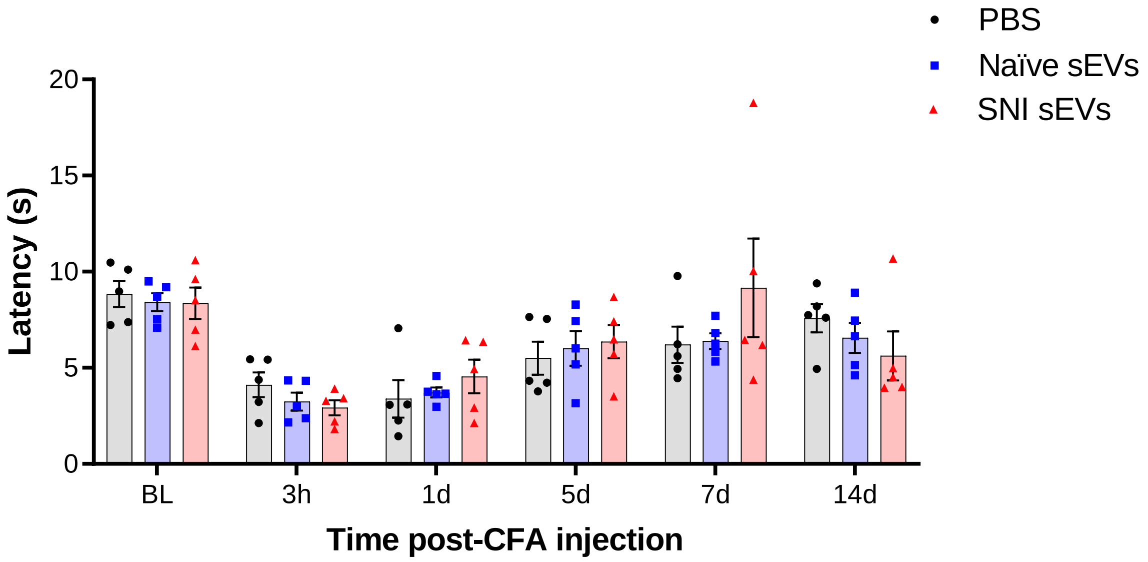

Supplementary Fig. 7

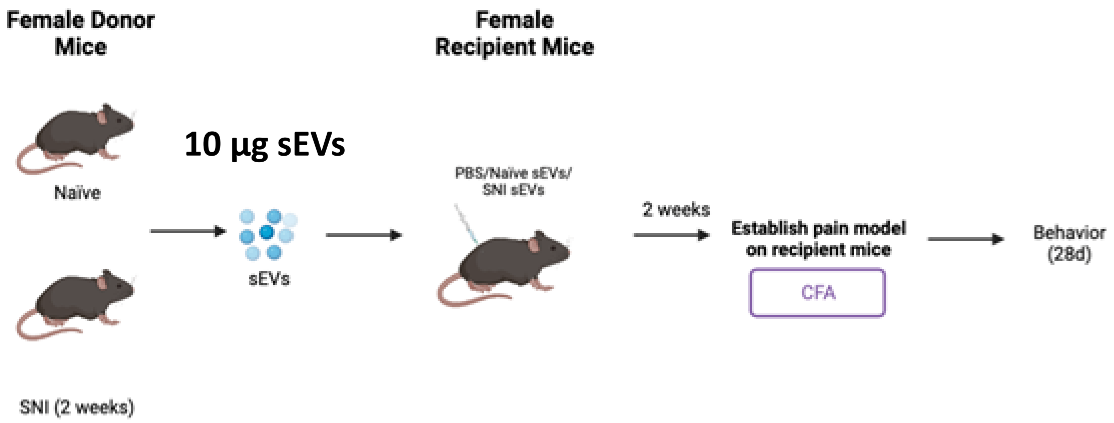


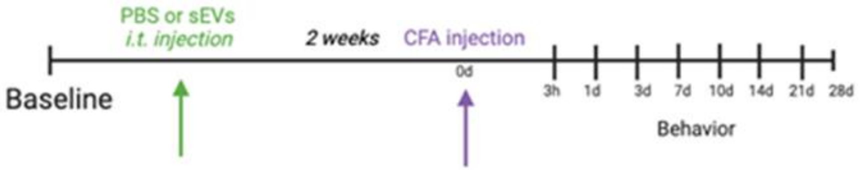


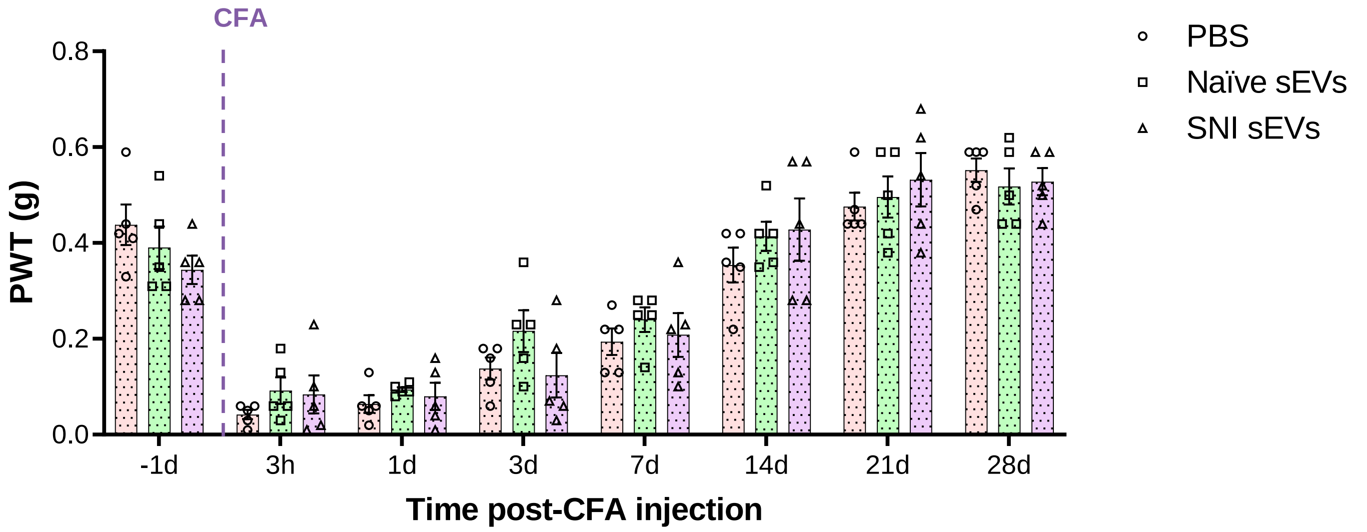


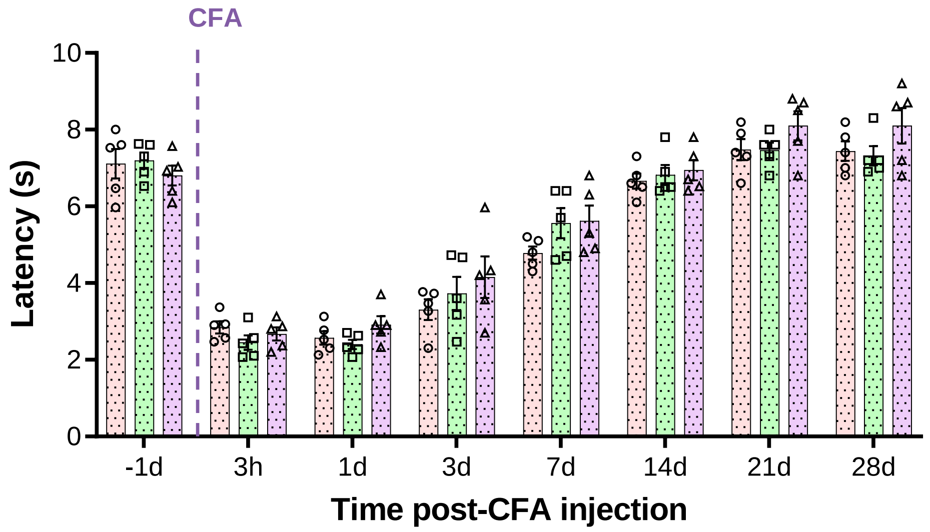

Supplementary Fig. 8

SC ipsi/contra

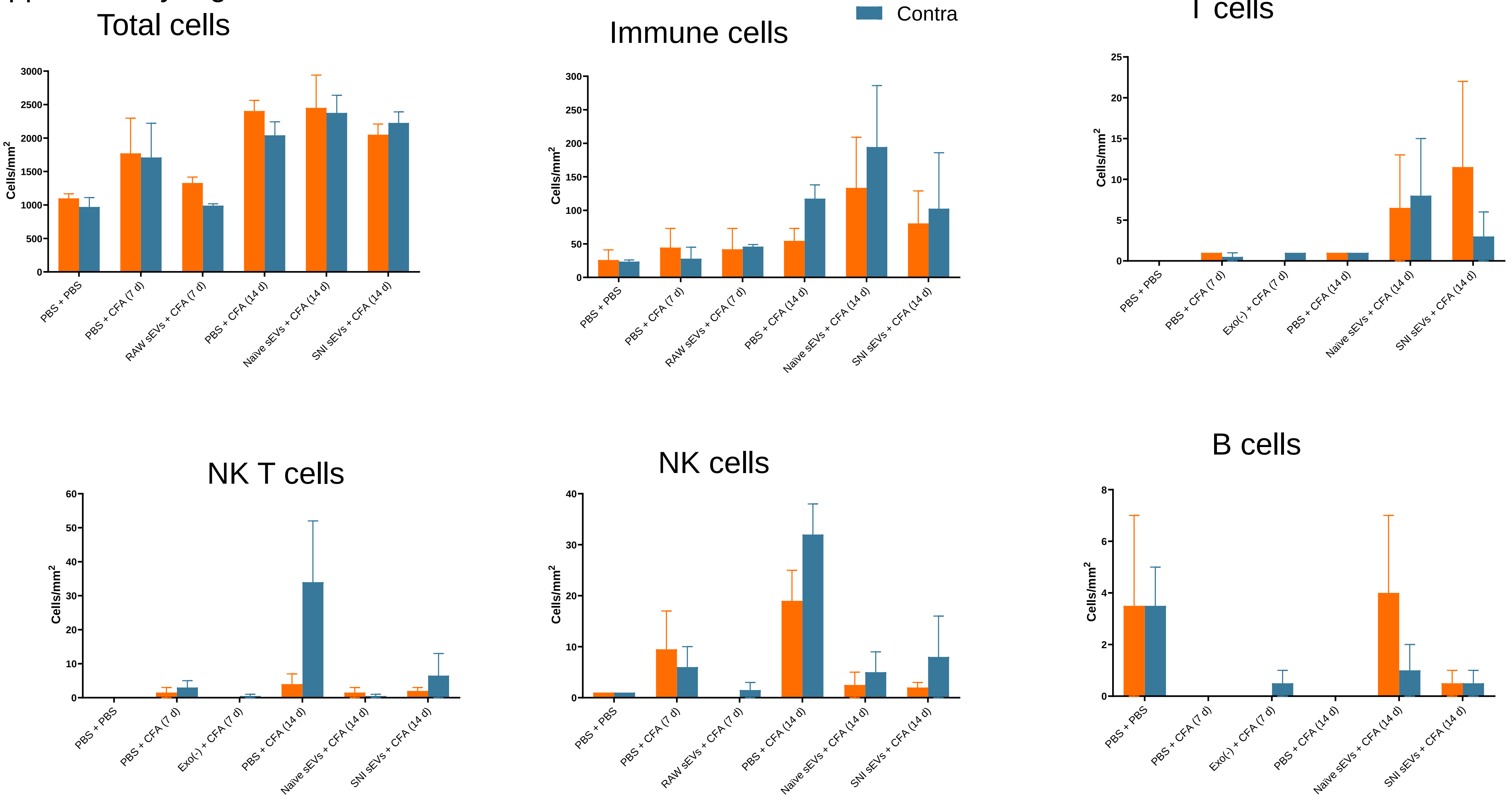

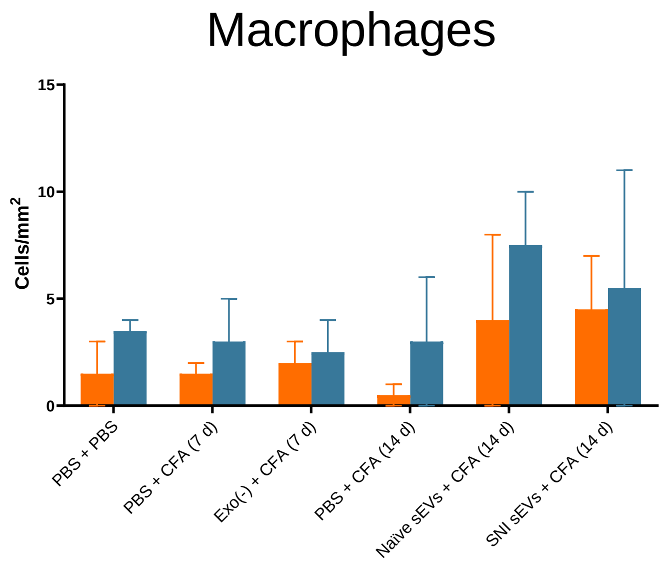
Macrophages
Cells/mm$^2$
0
5
10
15
PBS + PBS
PBS + CFA (7 d)
Exo(-) + CFA (7 d)
PBS + CFA (14 d)
Naïve sEVs + CFA (14 d)
SNI sEVs + CFA (14 d)

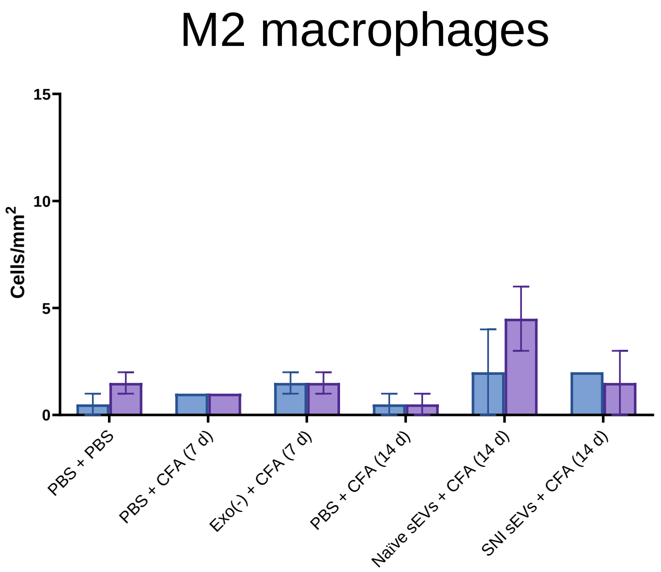
M2 macrophages
Cells/mm$^2$
0
5
10
15
PBS + PBS
PBS + CFA (7 d)
Exo(-) + CFA (7 d)
PBS + CFA (14 d)
Naïve sEVs + CFA (14 d)
SNI sEVs + CFA (14 d)

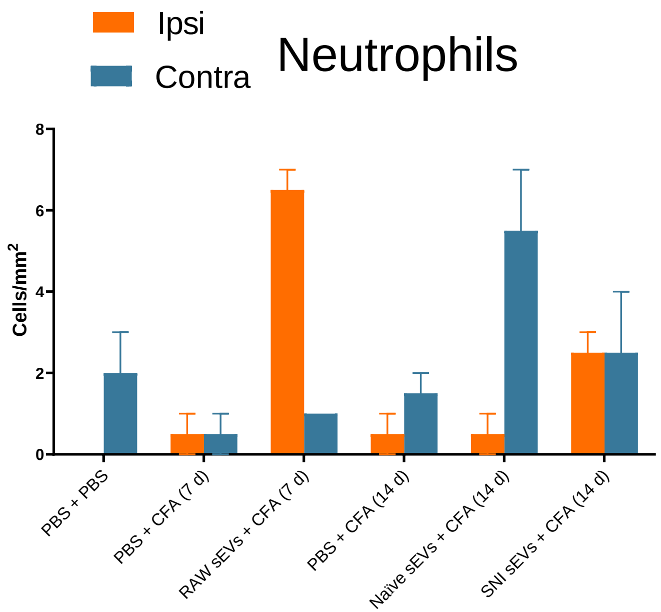
Ipsi
Contra
Neutrophils
Cells/mm$^2$
0
2
4
6
8
PBS + PBS
PBS + CFA (7 d)
RAW sEVs + CFA (7 d)
PBS + CFA (14 d)
Naïve sEVs + CFA (14 d)
SNI sEVs + CFA (14 d)

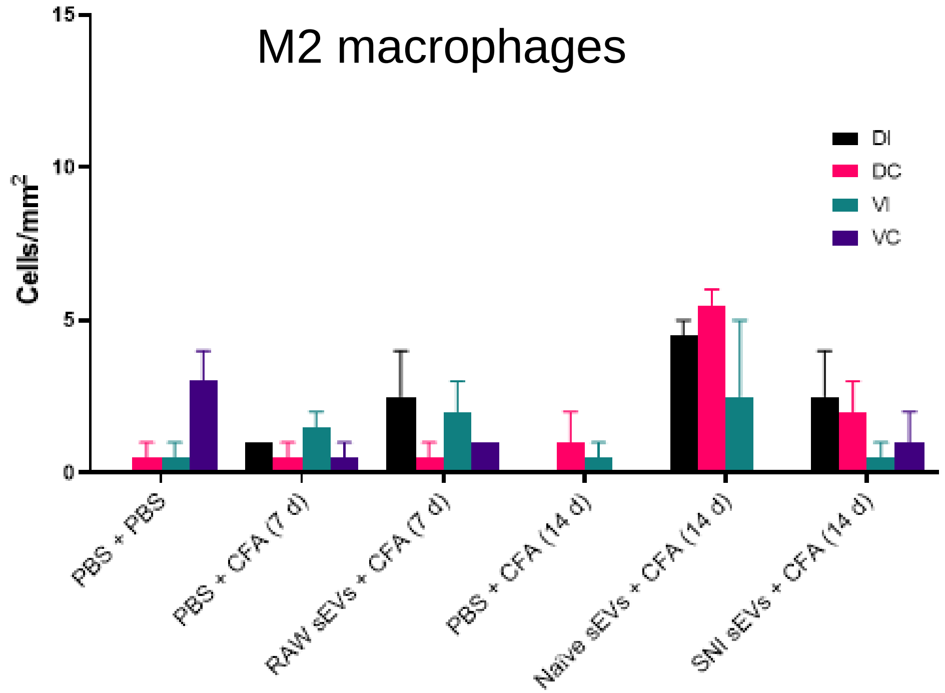
M2 macrophages
Cells/mm$^2$
0
5
10
15
DI
DC
VI
VC
PBS + PBS
PBS + CFA (7 d)
RAW sEVs + CFA (7 d)
PBS + CFA (14 d)
Naïve sEVs + CFA (14 d)
SNI sEVs + CFA (14 d)

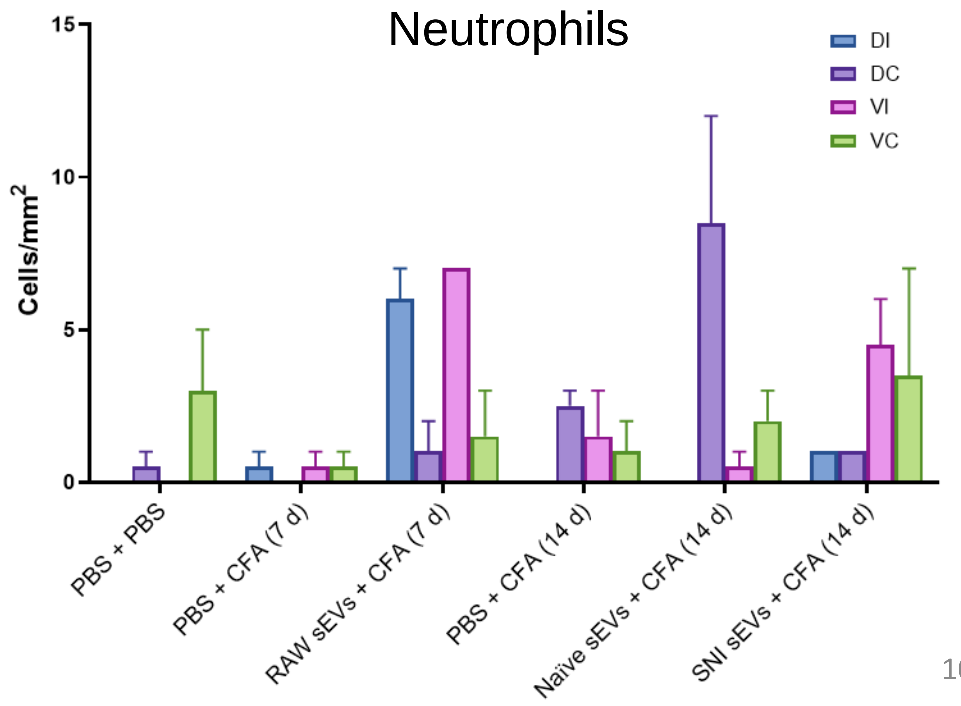
Neutrophils
Cells/mm$^2$
0
5
10
15
DI
DC
VI
VC
PBS + PBS
PBS + CFA (7 d)
RAW sEVs + CFA (7 d)
PBS + CFA (14 d)
Naïve sEVs + CFA (14 d)
SNI sEVs + CFA (14 d)

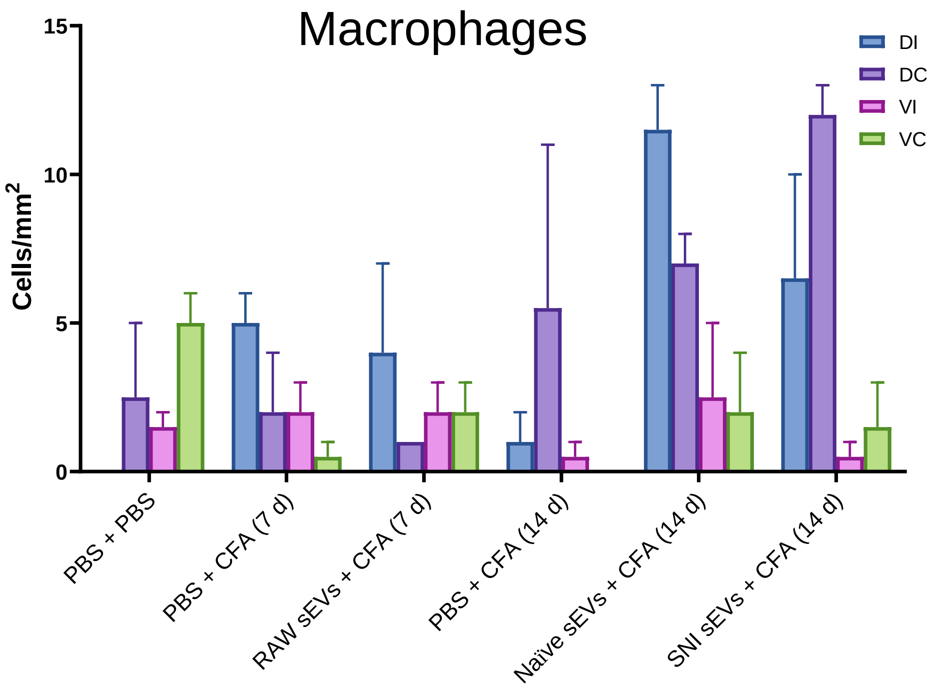
Macrophages
DI
DC
VI
VC
Cells/mm²
15
10
5
0
PBS + PBS
PBS + CFA (7 d)
RAW sEVs + CFA (7 d)
PBS + CFA (14 d)
Naïve sEVs + CFA (14 d)
SNI sEVs + CFA (14 d)

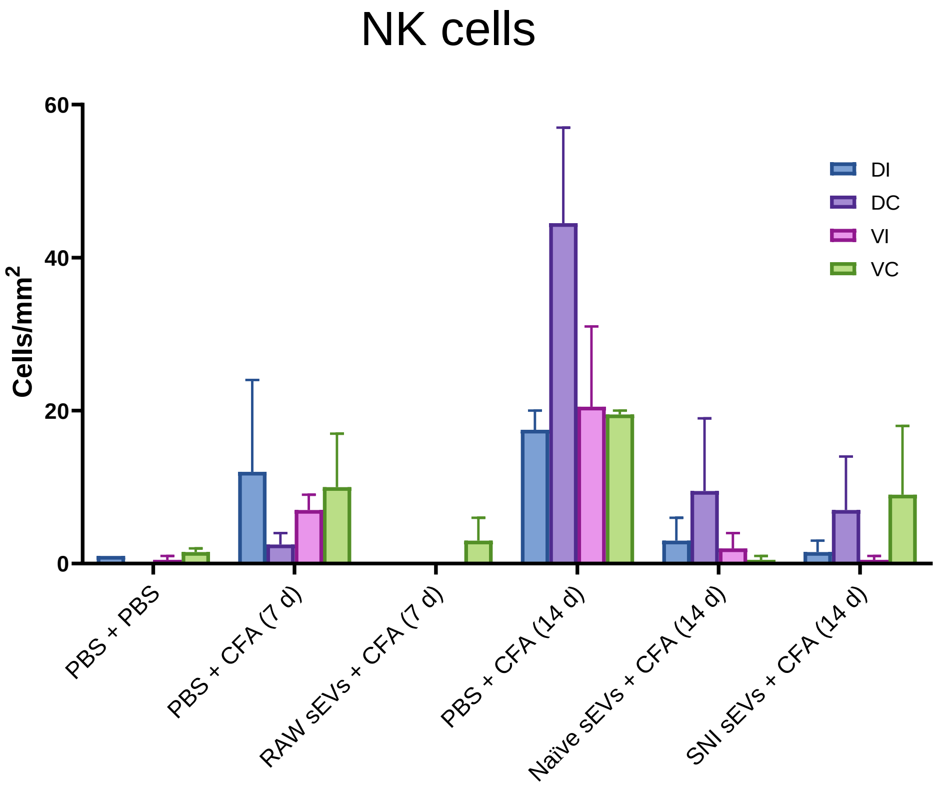
NK cells
DI
DC
VI
VC
Cells/mm²
60
40
20
0
PBS + PBS
PBS + CFA (7 d)
RAW sEVs + CFA (7 d)
PBS + CFA (14 d)
Naïve sEVs + CFA (14 d)
SNI sEVs + CFA (14 d)

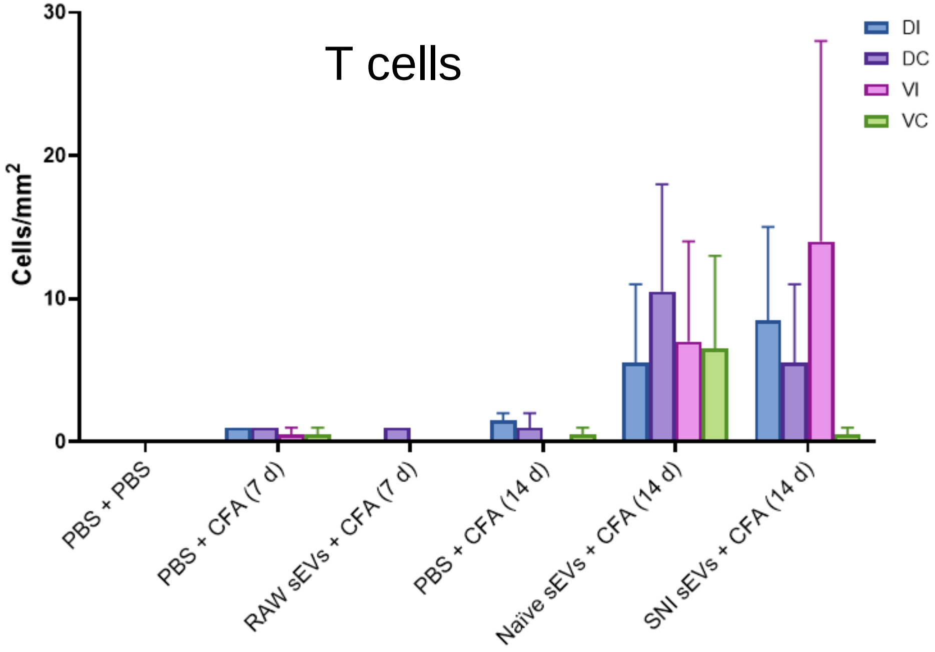
T cells
DI
DC
VI
VC
Cells/mm²
30
20
10
0
PBS + PBS
PBS + CFA (7 d)
RAW sEVs + CFA (7 d)
PBS + CFA (14 d)
Naïve sEVs + CFA (14 d)
SNI sEVs + CFA (14 d)

Supplementary Fig. 9

a

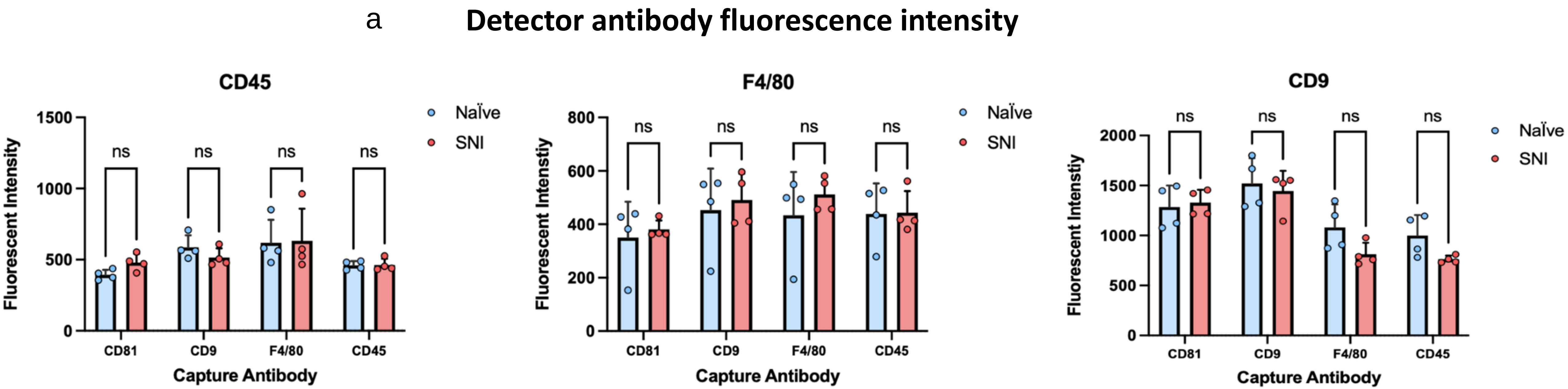


b

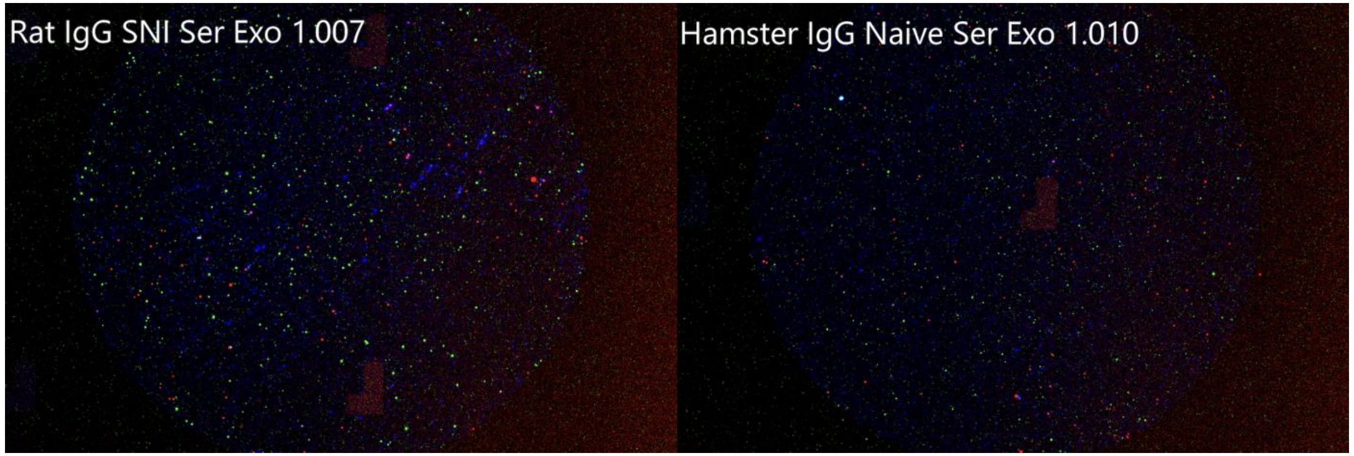

Supplementary Fig. 10

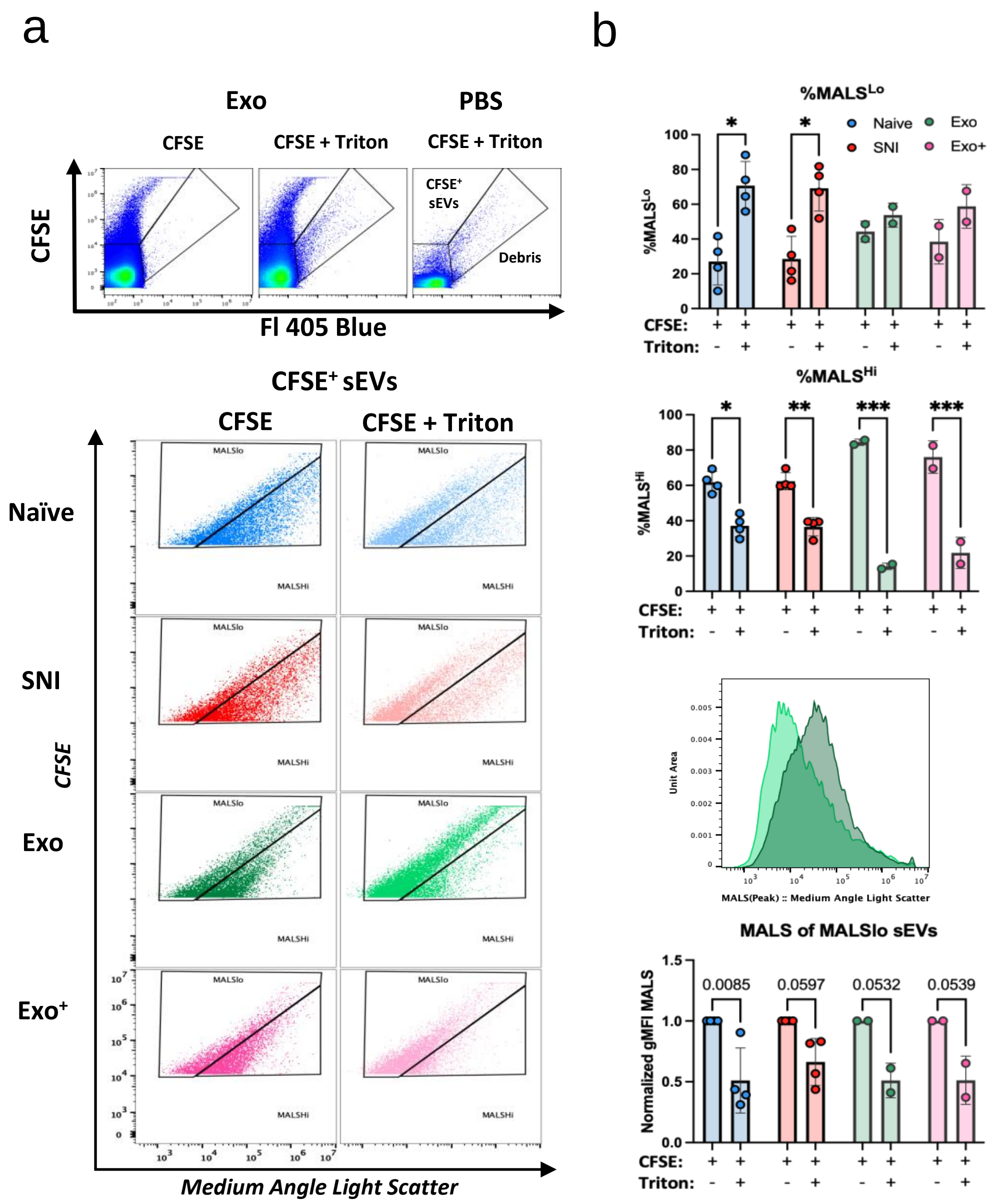
