## Supplemental Tables 4 and 5 for "Inflammatory pain resolution by mouse serum-derived small extracellular vesicles"

Supplementary Table 4

| # | **L4-L5 spinal cord and DRG from recipient male mice** |
|---|---|
| 1 | Naïve control |
| 2 | PBS (*i.t.)* + 7 days post CFA |
| 3 | sEVs from RAW 264.7 macrophage cells (*i.t.)* + 7 days post CFA |
| 4 | PBS (*i.t.)* + 14 days post CFA |
| 5 | sEVs from naïve donor mice (*i.t.)* + 14 days post CFA |
| 6 | sEVs from SNI model (2 wks post SNI; *i.t.*) + 14 days post CFA |

**Supp Table 4** Experimental design showing treatment groups in recipient mice. L4-L5 spinal cord and DRG from recipient male mice were obtained at time points that were most efficacious in behavior testing (n=2). sEVs from RAW 264.7 were included as a positive control for homogenous sEV source of macrophage under culture conditions.

### Supplementary Table 5

| Population | Marker definition |
|---|---|
| Immune cells | CD45+ |
| T cells | CD45+CD3+CD335- |
| CD4+T cells | CD45+CD3+CD335-CD4+CD8- |
| CD8+T cells | CD45+CD3+CD335-CD4-CD8+ |
| T reg | CD45+CD3+CD335-CD4+CD8-FOXP3+ |
| Activated T cells | CD45+CD3+CD335-MHCII+ |
| NKT cells | CD45+CD3+CD335+ |
| NK cells | CD45+CD3-CD335+ |
| Activated NK cells | CD45+CD3-CD335+MHCII+ |
| B cells | CD45-CD3-CD335-B220+CD19+ |
| Monocytes /Macrophages | CD45+CD3-CD335-CD11c-F4/80+CD11b+ |
| M2 macrophages | CD45+CD3-CD335-CD11c-F4/80+CD11b+CD206+ |
| Neutrophils | CD45+CD3-CD335-F4/80-CD11b+Gr1+ |
| Proliferating immune cells | CD45+Ki67+ |
| Neurons | NeuN+ |
| Microglia | CD45+CD3-CD335-F4/80-CD11b+TMEM119+ |

**Supp Table 5** The 16-plex panel of markers used to identify the populations in ChipCytometry
